# Contrasting epigenetic dynamics in chromosome arms and pericentromeres during ROS1-induced plant immune memory

**DOI:** 10.1101/2025.08.19.671022

**Authors:** Adam Hannan Parker, Peijun Zhang, Kwok Yin Man, Louis Tirot, Stephen A Rolfe, Lisa M Smith, Samuel W Wilkinson, Jurriaan Ton

## Abstract

Immune memory enables plants to respond more effectively to recurrent pathogen infections. Epigenetic changes in DNA methylation and small RNAs have been implicated, yet how their temporal dynamics govern the establishment and erasure of immune memory remains unclear. Using a chemically inducible system in Arabidopsis to activate the DNA demethylase REPRESSOR OF SILENCING 1 (ROS1), we establish immune memory against biotrophic pathogens that persists for up to two weeks. Memory establishment coincides with reduced small RNA accumulation and DNA methylation along chromosome arms, revealing gene targets involved in salicylic acid–dependent immunity and DNA damage repair. These changes are progressively reversed during immune memory erasure. In contrast, transposon-rich pericentromeres retain elevated small RNA and DNA methylation levels, consistent with a CLASSY3-associated redistribution of RNA-directed DNA methylation. Our findings show that transient ROS1 activity drives contrasting epigenetic responses across chromosomal regions, shaping both the establishment and erasure of plant immune memory.

## Introduction

After recovering from stress by pests or pathogens, plants can acquire an enhanced defensive capacity to better resist similar challenges later in life. This induced resistance is based on a form of immune memory, which spans the period between recovery and secondary challenge^1,2^. Depending on the plant species and the intensity of the stress experienced, this immune memory can persist in newly formed tissues for extended periods after the initial exposure to stress^3–8^. A growing body of evidence indicates an important role for epigenetic mechanisms in regulating plant immune memory^6,9^. In particular, cytosine DNA methylation has emerged as a key regulator due to its dynamic response to environmental stress and its capacity for stable transmission across both mitotic and meiotic cell divisions^6,9–13^. Because this memory can incur a fitness cost, epigenetically controlled immune memory is also transient, often reversing in the absence of stress^7,14,15^. The mechanisms underpinning the erasure of immune memory remain unknown.

In plants, cytosine DNA methylation is established by RNA-dependent DNA methylation (RdDM) pathways, which depend on non-coding small RNAs (sRNAs) of 21-24 nucleotides (nt) in length originating from RNA POLYMERASE-II (Pol-II) or Pol-IV transcripts^16,17^. Pol-IV is targeted to specific tissues and genomic loci by the chromatin remodelling factors CLASSY 1-4 (CLSY1-4)^18,19^. These sRNAs associate with ARGONAUTE proteins and guide them to nascent transcripts produced by Pol-V that share sequence similarity, leading to the recruitment of DOMAINS REARRANGED METHYLTRANSFERASE 1 and/or 2 (DRM1/2) and the *de novo* methylation of adjacent cytosines^16,20^. Once established, DNA methylation is maintained via different pathways depending on the cytosine context (CG, CHG, or CHH; H = A, T, or C). METHYLTRANSFERASE 1 (MET1) and CHROMOMETHYLASE 3 (CMT3) maintain DNA methylation in CG and CHG contexts, respectively, while CMT2 and RdDM machinery maintain DNA methylation at CHH sites^16,21^. DNA methylation is crucial for the silencing of transposable elements (TEs) – mobile genetic elements that cause large-scale modifications in the genome when activated^9^. In non-stressed vegetative tissues of *Arabidopsis thaliana* (Arabidopsis), RdDM activity is largely confined to gene-transposon boundaries in chromosome arms, whereas CMT-dependent DNA methylation primarily targets TE-rich heterochromatic regions near the centromeres (pericentromeres)^18,21,22^.

The removal of DNA methylation is mediated by bifunctional 5-methylcytosine DNA glycosylase/lyases that act through the base excision repair pathway^23^. In Arabidopsis, there are four of these DNA demethylases: REPRESSOR OF SILENCING 1 (ROS1), DEMETER (DME), DEMETER-LIKE 2 (DML2), and DML3^23^. Of these, ROS1 is the most abundant in leaves, where it actively antagonises RdDM at gene-transposon boundaries^23–26^. Numerous studies have demonstrated that mutation of *ROS1* (*ros1* mutants) causes elevated DNA methylation at TEs in the proximity of protein coding genes, resulting in reduced activation of environmentally responsive genes following exposure to biotic and abiotic stress stimuli^5,23,25^. Furthermore, *ros1* mutants cannot establish long-term immune memory against *Spodoptera littoralis* caterpillars upon seedling treatment with jasmonic acid (JA)^7^, and fail to develop heritable immune memory against the oomycete pathogen *Hyaloperonospora arabidopsidis* (*Hpa*) after parental infection with the bacterium *Pseudomonas syringae* pv. *tomato* DC3000 (*Pst*)^5^. Hence, ROS1-dependent removal of DNA methylation is emerging as a key feature of adaptive immune responses to biotic stress. However, *ros1* mutant lines exhibit constitutive DNA hypermethylation at loci that are typically lowly or moderately methylated in wild-type plants, which means the role of highly methylated ROS1 targets during biotic stress remains unexplored^6,9^. Moreover, the constitutive nature of *ros1* mutation limits the ability to study the dynamic and reversible epigenetic changes associated with ROS1-induced immune memory.

In this study, we have used a chemically inducible gene construct to transiently activate ROS1 in Arabidopsis, thereby establishing epigenetic immune memory against pathogens. We show that the establishment of this memory is associated with loss of DNA methylation and RdDM-associated sRNAs in chromosome arms alongside transcriptional activation of key defence genes, consistent with *cis*-regulation by ROS1. By contrast, transient ROS1 induction has opposing effects on sRNA accumulation and DNA methylation in transposon-rich pericentromeric regions, involving CLSY3-dependent redistribution of RdDM. Together, these findings reveal the dynamic epigenomic responses underlying ROS1-driven immune memory and identify molecular targets and mechanisms that may be leveraged for crop protection strategies.

## Results

### The *XVE:ROS1-YFP* system allows for transient and dose-dependent activation of ROS1

To induce ROS1 activity in a dose-dependent manner, we transformed Arabidopsis with a 17β-estradiol (E2)-inducible ROS1-YFP construct, which is based on the XVE transactivating system (Figure 1a)^27,28^. Treatment of *XVE:ROS1-YFP* plants with E2 induces a conformational change in the XVE receptor, enabling it to bind to LexA operator sequences which activates *ROS1-YFP* expression (Figure 1a, 1b). We transformed wild-type Col-0 to avoid the pleiotropic effects on epigenetic homeostasis and plant immunity of constitutive ROS1 loss in *ros1* mutants^24,25^.

**Figure 1.**
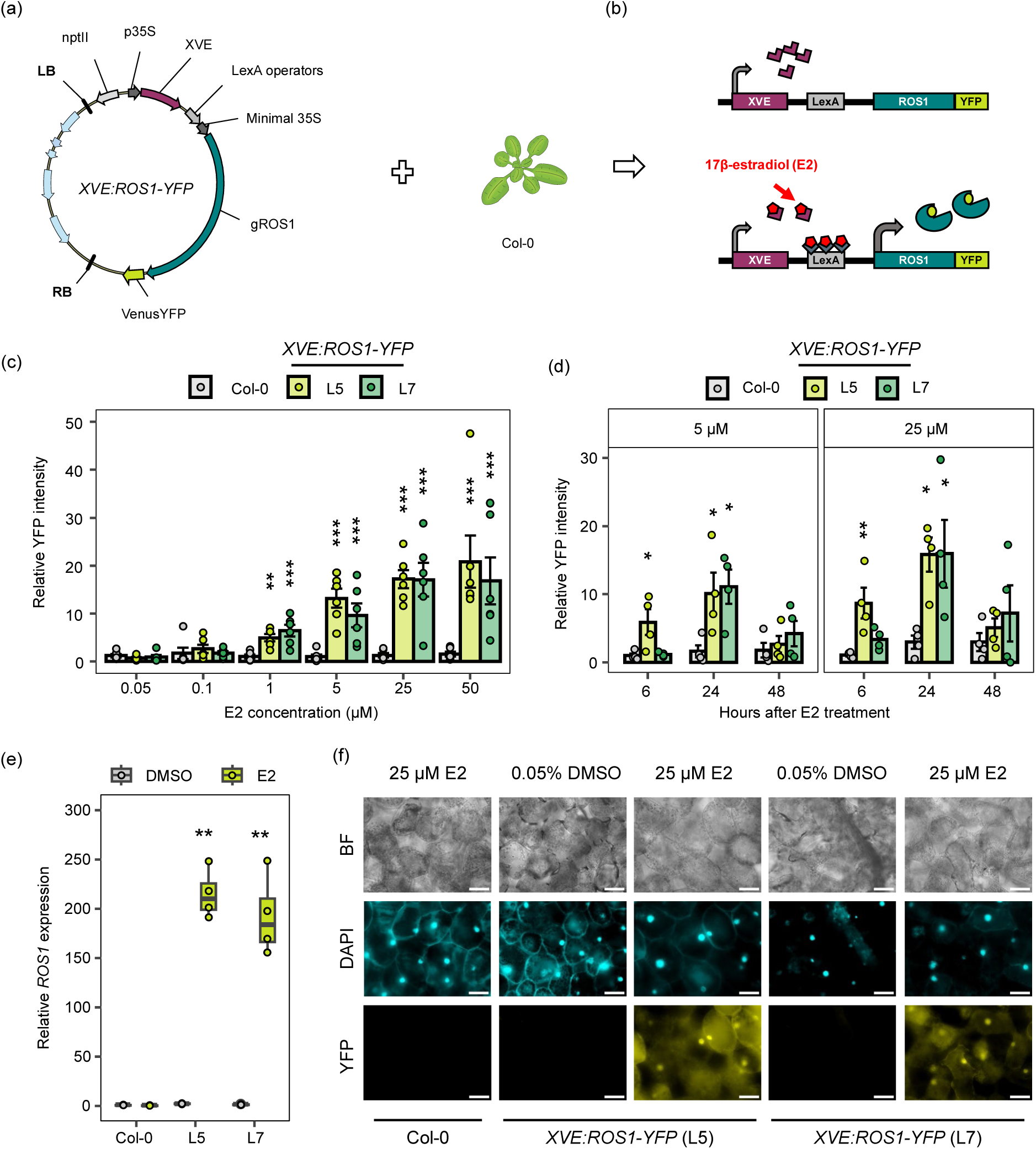
Characterisation of *XVE:ROS1-YFP* lines. **(a)** Plasmid map of estradiol (E2)-inducible *XVE:ROS1-YFP* construct used to transform Col-0. **(b)** Schematic of the XVE E2-inducible system. **(c)** ROS1-YFP protein accumulation at 24 hours after treatment of 7-day-old hydroponically grown seedlings (Col-0, *XVE:ROS1-YFP* line 5; L5, and *XVE:ROS1-YFP* line 7; L7) with increasing E2 concentrations. Data points represent relative YFP fluorescence intensities, normalised by plant area, from biologically replicated seedlings (n = 6). Shown are data relative to the mean YFP fluorescence intensity value from DMSO-treated Col-0 seedlings (background fluorescence). Asterisks indicate a statistically significant difference in relative YFP fluorescence intensity compared to Col-0 plant (Tukey; * *p_adj_* < 0.05, ** *p_adj_* < 0.01, *** *p_adj_* < 0.001). **(d)** ROS1-YFP protein accumulation at 6-, 24-, and 48-hours after treatment (hpt) with 5 μM or 25 μM E2 in 7-day-old hydroponically grown seedlings. Data points represent YFP intensity values from biological replicates (n = 4) relative to the mean YFP intensity value from DMSO-treated controls within a given genotype and timepoint. For each time-point, pairwise two-sided t-tests against Col-0 were performed and *p*-values were adjusted by false discovery rate (FDR; * *q* < 0.05, ** *q* < 0.01, *** *q* < 0.001). In (c) and (d), bars represent mean values with error bars representing standard error of the mean. **(e)** Quantification of *ROS1* transcripts by RT-qPCR at 24 hpt after spraying soil-grown Col-0, L5, and L7 plants with 25 μM E2 or 0.05% DMSO at 14 days after sowing (DAS). Data points represent relative expression values of individual biological replicates (n=4) normalised to the mean relative expression value of DMSO-treated Col-0 plants. Asterisks indicate a statistically significant difference between treatments within a genotype (two-sided t-test; ** *p* < 0.01, *** *p* < 0.001). **(f)** ROS1-YFP protein accumulation was imaged at 24 hours by epi-fluorescence microscopy after the same treatment as (e). Brightness and contrast of all images were adjusted as follows: bright-field (BF) images, +20% and +10%; DAPI images, +60% and +70%; YFP images, +40% and +70%, respectively. DAPI and YFP images were pseudo-coloured cyan and yellow, respectively. Scale bars, 50 μm

To validate the inducibility of the *XVE:ROS1-YFP* system, we quantified E2-induced ROS1-YFP accumulation by epifluorescence microscopy in hydroponically grown seedlings. Relative YFP fluorescence was measured 24 hours after E2 treatment across a range of E2 concentrations in two independent *XVE:ROS1-YFP* lines – line 5 (L5) and line 7 (L7) – and in Col-0 (negative control; Figure 1c). A two-way ANOVA revealed a significant genotype × treatment interaction for relative YFP fluorescence (F = 6.2; df = 10,90; *p* < 0.001). Both L5 and L7 exhibited fluorescence levels significantly above background (Col-0) at E2 concentrations ≥1 µM (Tukey post-hoc, *p_adj_* < 0.05).

To assess the persistence of E2-induced ROS1-YFP accumulation, we quantified YFP fluorescence in L5, L7, and Col-0 plants relative to DMSO-treated control plants at 6-, 24-, and 48-hours post treatment (hpt) with 5 or 25 µM E2 (Figure 1d). We performed a repeated-measures ANOVA with genotype and E2 dose as between-subject factors, and hpt as a within-subject factor. The three-way interaction between genotype, E2 dose, and hpt was not significant (*F* = 0.1; df = 4, 36; *p* > 0.05), but there was a significant genotype × hpt interaction (*F* = 6.4; df = 4, 36; *p* < 0.001), indicating that ROS1-YFP dynamics varied by genotype over time, independent of E2 dose. YFP fluorescence in L5 and L7 significantly increased at 24 hpt (*q* < 0.05) before returning to Col-0 levels by 48 hpt (*q* > 0.05). Only L5 showed significantly higher YFP fluorescence than Col-0 at 6 hpt (*q* < 0.05; Figure 1d). These results indicate that while early induction kinetics differ slightly between the two *XVE:ROS1-YFP* lines, E2 consistently induces transient ROS1-YFP accumulation, peaking at 24 hpt, with expression ceasing or the protein degrading by 48 hpt, regardless of dose.

To confirm the inducibility of the *XVE:ROS1-YFP* construct under conditions used for plant-pathogen assays, we sprayed soil-grown plants with 25 μM E2 at 14 days after sowing (DAS). At 24 hpt, *ROS1* transcript levels were quantified by RT-qPCR, revealing a ∼200-fold increase in *XVE:ROS1-YFP* L5 and L7 (two-sided t-test, *p* < 0.05; Figure 1e). In contrast, Col-0 plants showed no change in *ROS1* expression or YFP signal upon this E2 treatment (two-sided t-test, *p* < 0.05; Figures 1e-f). DMSO-treated L5 and L7 showed a minor but statistically significant increase in *ROS1* transcripts (1.7x and 2.3x, respectively) relative to Col-0 (two-sided t-test, *p* <0.05; Figure 1e), but this did not correspond with increased YFP signal in leaves (Figure 1f). By comparison, E2-treated L5 and L7 displayed strong YFP signals at 24 hpt that co-localised with nuclear DAPI staining, confirming nuclear localisation of ROS1-YFP (Figure 1f). Together, these results indicate that despite mild leakiness, the *XVE:ROS1-YFP* system is highly E2-dependent, enabling robust and transient accumulation of ROS1-YFP in Arabidopsis under different growth conditions.

### Ectopic ROS1 induction results in transient immune memory against (hemi-)biotrophic pathogens

The *ros1-4* mutant of Arabidopsis is more susceptible to the biotrophic oomycete *Hpa* and the hemi-biotrophic bacterium *Pst*^5,25^. To examine whether ROS1 overexpression induces resistance to these pathogens, we treated Col-0, L5, and L7 seedlings twice with E2 or DMSO before spray-inoculating 2-days-post-treatment (2 dpt) with either *Hpa* (Waco9) or bioluminescent *Pst luxCDABE*^29,30^ (*Pst-Lux*; Figure 2a). Colonisation by *Hpa* was analysed microscopically at 5 days post inoculation (dpi) by assigning trypan blue-stained leaves to *Hpa* colonisation classes, ranging from I (no colonisation) to IV (severe colonisation and sporulation)^31^, whereas bacterial colonisation by *Pst-Lux* was quantified non-destructively over multiple days, using bioluminescence intensity as a quantitative measure of colonisation^29,30^ (Figures 2c, 2d).

**Figure 2.**
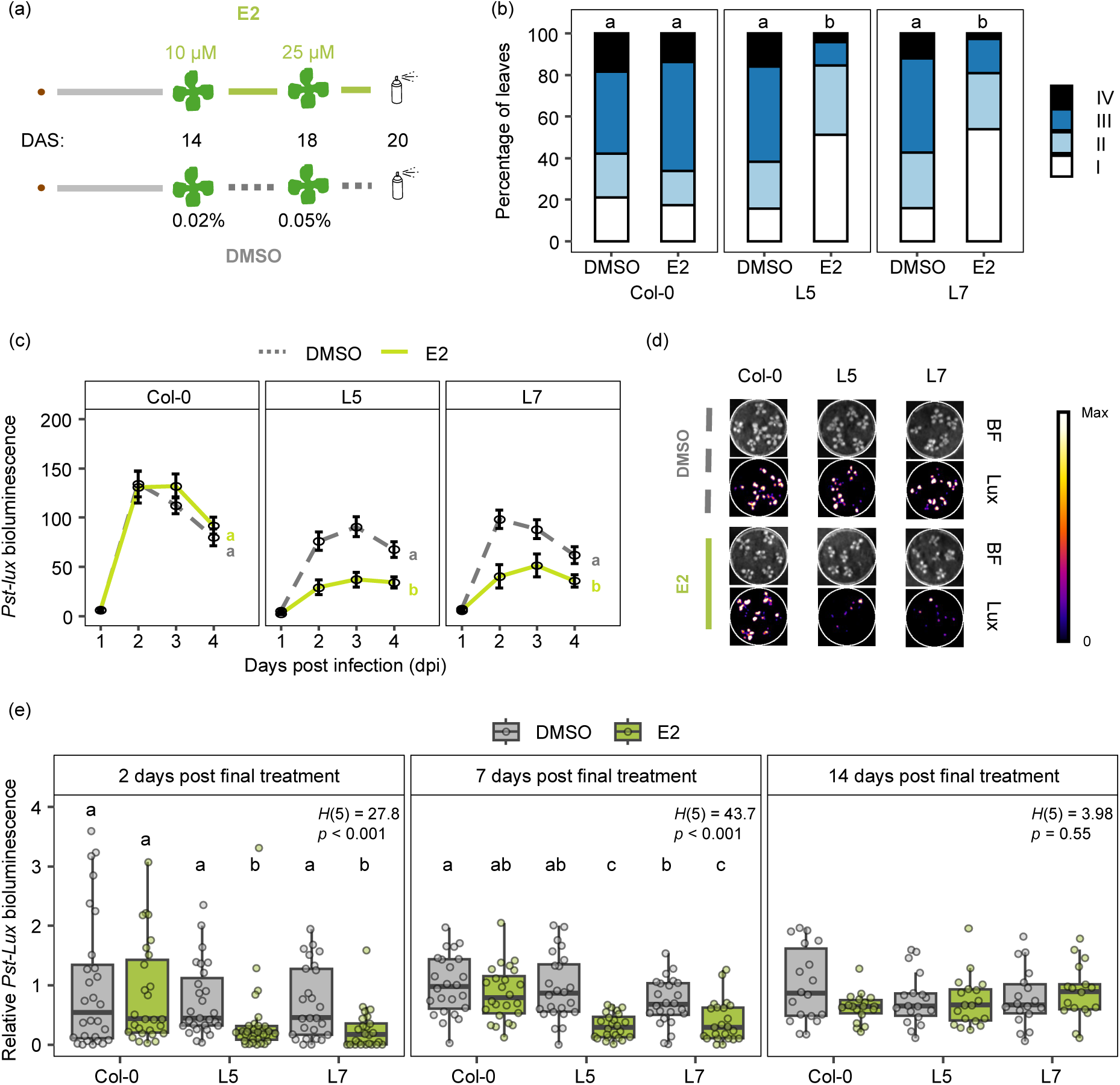
Quantification of ROS1-induced immune memory against (hemi-)biotrophic pathogens. **(a)** Experimental setup to quantify induced resistance by E2-induced ROS1 against the (hemi)biotrophic pathogens *Hpa* and *Pst-Lux*. Col-0, L5, and L7 plants were treated twice with E2 or DMSO at 14 days after sowing (DAS) (10 µM E2 or 0.02% DMSO) and 18 DAS (25 µM E2 or 0.05% DMSO). 2 days post the final treatment (20 DAS; 2 dpt), plants were inoculated with *Hpa* or *Pst-Lux*. **(b)** At 5 dpi with *Hpa*, individual leaves (72-80 leaves from 4 pots containing 3-5 individual plants) were stained by trypan-blue and microscopically assigned to 4 different *Hpa* colonisation classes: I: no hyphal colonisation; II: limited hyphal colonisation, ≤ 8 conidiophores; III: extensive hyphal colonisation, > 8 conidiophores; class IV: extensive hyphal colonisation with conidiophores and oospore(s)^31^. Shown are relative frequencies of leaves across the *Hpa* colonisation classes. Different letters indicate statistically significant differences in frequency distribution (Fisher’s exact test; FDR-correction; *q* < 0.05). **(c)** Colonisation by *Pst-Lux* was quantified 1-4 dpi by bioluminescence intensity. Points with error bars represent mean *Pst-Lux* bioluminescence values (n = 31-41) ± standard error of the mean. Different letters indicate statistically significant differences in the progression of *Pst-Lux* colonisation between 2-4 dpi (linear mixed effects model; estimated marginal means post-hoc; Tukey *p_adj_* < 0.05). **(d)** Representative examples of *Pst-Lux* bioluminescence Col-0, L5, and L7 at 2 dpi. Bright field (BF) images were used to normalise *Pst-Lux* bioluminescence intensities by plant area. **(e)** Duration of ROS1-induced immune memory against *Pst-Lux*. Col-0, L5, and L7 plants were treated with DMSO or E2, as shown in (a), and inoculated with *Pst-Lux* 2, 7, or 14 days post final treatment (2 dpt, 7 dpt, 14 dpt). *Pst-Lux* colonisation was quantified by bioluminescence 2- or 3-dpi and expressed relative to the *Pst-Lux* bioluminescence of DMSO-treated Col-0 within each timepoint. If a significant treatment effect on relative *Pst-Lux* colonisation was detected (Kruskal-Wallis, *p* < 0.05), Conover-Iman post-hoc tests with a Benjamini-Hochberg correction were performed. Different letters indicate significant differences (*p_adj_* < 0.05).

Treatment of Col-0 with either DMSO or E2 resulted in similar levels of colonisation by *Hpa* (Fisher’s exact-test, *q* > 0.05), indicating that E2 itself has no direct effect on *Hpa* (Figure 2b). In contrast, E2 treatment of both *XVE:ROS1-YFP* lines resulted in strongly reduced *Hpa* colonisation compared to DMSO-treated plants (Fisher’s exact tests: L5: *q* < 0.001; L7: *q* < 0.001), while DMSO-treated *XVE:ROS1-YFP* lines were as susceptible as DMSO- and E2-treated Col-0 plants (Fisher’s exact tests, *q* > 0.05; Figure 2b). Hence, ectopic induction of ROS1 by two successive E2 treatments induces resistance against *Hpa*.

A linear mixed-effects model, which accounted for dpi and the random effect of pot was fitted to our *Pst-Lux* bioluminescence data from two to four dpi. Likelihood ratio tests between the full model and reduced models revealed statistically significant effects of genotype (Col-0, L5, L7; *Χ* ^2^ = 46.6, df = 4, *p* < 0.001), treatment (DMSO, E2; *Χ* ^2^ = 31.7, df = 3, *p* < 0.001), and the interaction between genotype and treatment (*Χ* ^2^ = 15.9, df = 2, *p* < 0.001). Pairwise post-hoc comparisons on the estimated marginal means confirmed that E2-treated *XVE:ROS1-YFP* lines sustained significantly lower levels of *Pst-Lux* colonisation compared to the corresponding DMSO controls (*p_adj_* < 0.05; Figure 2c). In contrast, DMSO- and E2-treated Col-0 plants sustained similar *Pst-Lux* colonisation dynamics (*p* > 0.05; Figures 2c, 2d). Characterisation of three additional independent *XVE:ROS1-YFP* lines (L6, L30 and L32) which showed varying levels of E2-induced YFP fluorescence confirmed that reduced ROS1 induction results in lower resistance to *Pst-Lux* (Supplementary Figure 1a-b). Thus, ectopic induction of ROS1 induces resistance against *Pst-Lux*.

Our initial bioassays used two successive E2 treatments with increasing concentrations, reasoning that sustained ROS1 activation would better mimic the progressive epigenetic changes triggered by escalating stress from a virulent pathogen. Single treatments – either 10 µM E2 at 14 DAS or 25 µM E2 at 18 DAS – did not consistently induce resistance against *Pst*-*Lux* at either 7- or 2-dpt (Supplementary Figure 1d-g). In contrast, both a repeated high-dose induction regime (four applications of 25 µM E2 between 14-20 DAS) and an incremental dual-dose regime (10 µM at 14 DAS followed by 25 µM at 18 DAS) reproducibly enhanced resistance in both lines at 2 dpt (Supplementary Figure 1d-g). Therefore, we adopted the dual-treatment protocol for all subsequent experiments to ensure a reliable resistance phenotype for downstream analyses.

To assess the durability of ROS1-induced immune memory, we challenged Col-0, L5, and L7 plants with *Pst-Lux* at 2 dpt, 7 dpt, or 14 dpt after the final treatment (Figure 2e). Resistance against *Pst-Lux* was evident in *XVE:ROS1-YFP* lines at both 2 dpt and 7 dpt, but not at 14 dpt (Figure 2e). Furthermore, progeny of E2-treated *XVE:ROS1-YFP* lines did not show enhanced resistance to either *Pst-Lux* or *Hpa* relative to progeny from DMSO-treated control plants (Supplementary Figure 2). These results indicate that, while transient ROS1 induction is sufficient to induce resistance against biotrophic pathogens, this immune memory is lost within 7-14 days. Thus, the *XVE:ROS1-YFP* system offers a tractable model to study the establishment, maintenance, and often-overlooked erasure of plant immune memory.

### The establishment of ROS1-induced immune memory is associated with major changes in the transcriptome and DNA methylome

To investigate the molecular drivers of ROS1-induced immune memory, Col-0, L5, and L7 plants were treated twice with E2 or DMSO (Figure 2a) and biologically replicated (n=3) aerial tissues were harvested 48 hours after the final treatment. DNA and RNA were extracted from the same pool of tissues before sequencing for DNA methylation (whole-genome bisulfite sequencing; WGBS), ribosome-depleted total RNA, and small RNAs (sRNAs). Principal component analysis (PCA) of all three datasets revealed that ROS1-induction was the primary source of variance (Supplementary Figure 3b-d). Crucially, E2- and DMSO-treated Col-0 samples clustered together in all three datasets, demonstrating that the global shifts in DNA methylation, gene expression, and sRNA abundance were driven by ectopic ROS1 induction rather than by E2 treatment itself.

E2-treated *XVE:ROS1-YFP* lines, but not Col-0, exhibited reduced DNA methylation across chromosome arms in all cytosine contexts (allC, CG, CHG, CHH) relative to DMSO-treated controls (Figure 3a; Supplementary Figure 4). In contrast, pericentromeric regions were less affected, with some loci displaying CHG hypermethylation in E2-treated L5 and L7 (Figure 3a; Supplementary Figure 4). DNA hypomethylation by ROS1 was less pronounced within gene bodies than in their flanking regions (Supplementary Figure 4a-b), whereas hypomethylation was stronger within TE bodies compared to their flanking regions (Supplementary Figure 4b). Analysis of differentially methylated regions (DMRs) identified thousands of hypomethylated DMRs in E2-treated L5 (11,067) and L7 (12,158) plants, but only 25 in E2-treated Col-0 plants. Few hypermethylated DMRs were detected across all genotypes: 22 in L5, 18 in L7, and 30 in Col-0 (Supplementary Figure 4c; Supplementary Data 1). After merging across all cytosine contexts, 4,971 overlapping hypomethylated DMRs were identified in E2-treated L5 and L7 plants (Supplementary Figure 5a). These consistent hypo-DMRs were highly enriched at gene-TE boundary regions, with the strongest enrichment occurring at TEs in the promoter regions of protein-coding genes (Supplementary Figure 5b). Analysis of two recently published ChIP-seq datasets for ROS1^26,32^ revealed significant enrichment of endogenous ROS1 binding within our hypo-DMRs relative to randomly selected genomic regions, in both independent datasets (Kruskal–Wallis *p* < 0.001; Conover–Iman *q* < 0.05; Supplementary Figure 5c-d). Thus, our results support previous characterisations of ROS1 in Arabidopsis^24,25,32^ and indicate that the recombinant ROS1-YFP protein is functionally active as a DNA demethylase.

**Figure 3.**
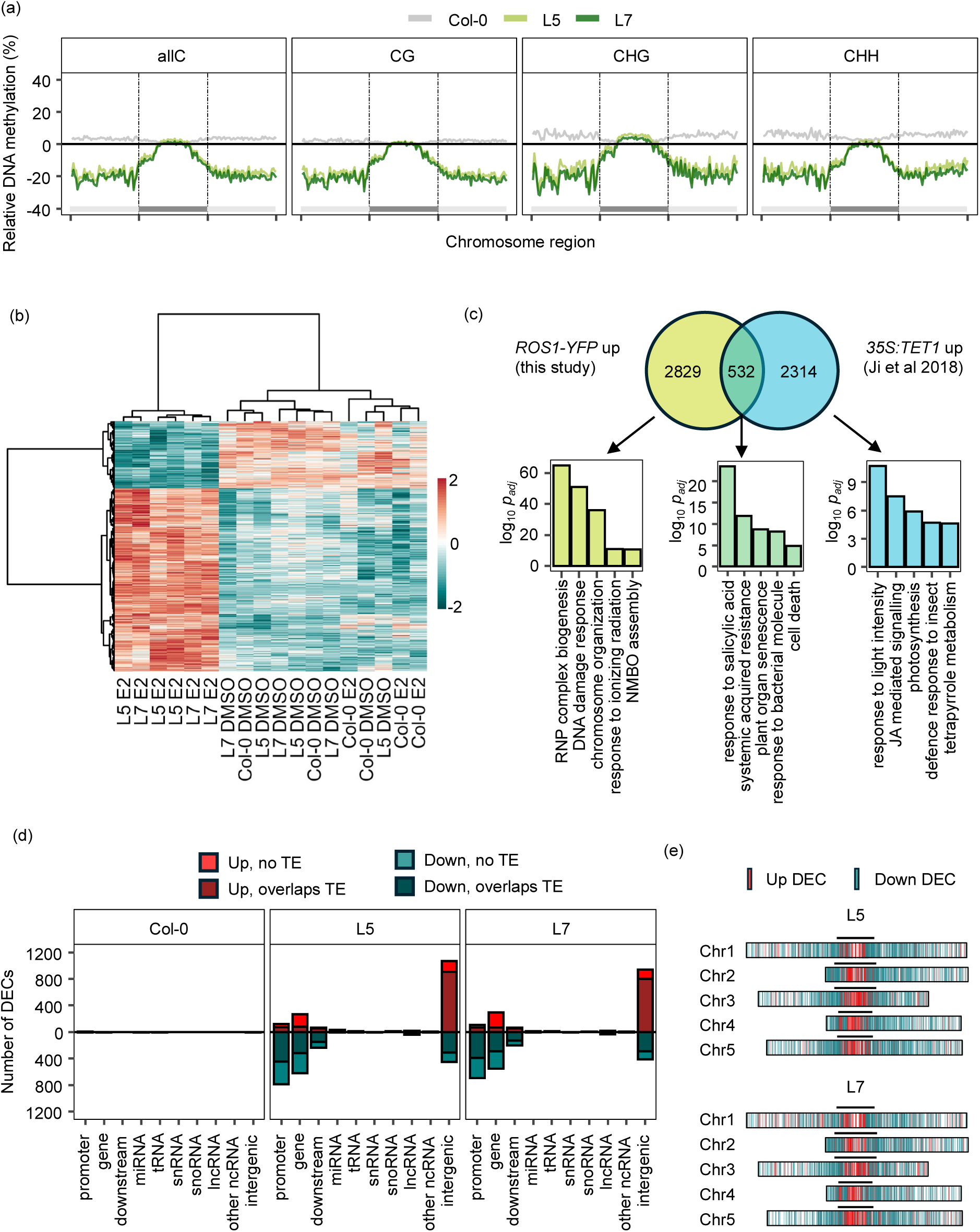
Genome-wide characterisation of the methylome, transcriptome, and sRNAome following transient ROS1 induction. **(a)** Chromosome-level metaplots of DNA methylation in E2-treated plants. Shown are the mean percentage differences in cytosine methylation relative to DMSO-treated plants in Col-0 (grey), L5 (light green), and L7 (dark green). For each chromosome, average DNA methylation was calculated for 50 equally sized bins in the left arm, the pericentromeric region, and the right arm before averaging across replicates (n=3) and chromosomes. **(b)** Heatmap projection of all 4,724 differentially expressed loci/genes (DEGs) between E2- and DMSO-treated plants within a genotype. Heatmap-projected values represent per gene Z-scores of transformed normalised read counts from 3 biological replicates for each genotype-treatment combination. Scores were clustered by both DEG (Euclidean, Ward) and sample (Pearson correlation, Ward). **(c)** Venn diagram showing the overlap of upregulated DEGs (*p_adj_* < 0.05) in E2-treated *XVE:ROS1-YFP* L5 and L7 (derived from figure b) and upregulated DEGs in *35S:TET1* lines (*p_adj_* < 0.05)^33^. Shown are the top 5 GO enrichment terms in order of statistical significance for biological process for the overlapping DEG set (green) and the unique DEG sets (yellow and blue *for XVE:ROS1-YFP* and *35S:TET1*, respectively). RNP = ribonucleoprotein; JA = jasmonic acid. **(d)** Annotation counts of differentially expressed sRNA clusters (DECs) from E2-treated L5 and L7, as compared to DMSO-treated L5 and L7, respectively. DECs were annotated using Araport11^37^, and TAIR10 TE annotations. Counts of up- and downregulated DECs are shown above and below the y = 0 line, respectively. **(e)** Distribution of DECs across the five Arabidopsis chromosomes for L5 and L7. Red and blue vertical rectangles superimposed on the chromosomes represent up- and downregulated DECs, respectively. Black lines above the chromosomes mark pericentromeric regions.

### The ROS1-induced transcriptome is enriched with genes controlling salicylic acid-dependent immunity and DNA damage response

E2-treatment of L5 and L7, but not Col-0, profoundly altered the transcriptional profile, with a strong correlation between both lines (*r* = 0.94, n = 4724, *p* < 0.001; Supplementary Figure 6a; Supplementary Data 2). Hierarchical clustering of all differentially expressed genes (DEGs) separated replicate samples of E2-treated *XVE:ROS1-YFP* lines from all other treatment groups (Figure 3b). This analysis identified two distinct DEG clusters: genes upregulated in E2-treated *XVE:ROS1-YFP* lines (‘up’ cluster; n = 3,460) and genes downregulated in these lines (‘down’ cluster; n = 1264). The majority of DEGs were protein-coding genes (95% and 96% in the up- and downregulated clusters, respectively), with the remainder consisting of TE-derived transcripts and non-coding RNAs (Supplementary Figure 6c; Supplementary Data 2). Gene ontology (GO) enrichment analysis showed that upregulated DEGs were significantly associated with biological processes related to chromatin organisation, DNA damage response, and immunity, whereas downregulated DEGs were primarily enriched for GO terms linked to photosynthesis (Supplementary Figure 6b; Supplementary Data 3).

Activation of the DNA damage repair pathway by ectopically induced ROS1 likely results from the enzyme’s DNA glycosylase/lyase activity^23^. To separate the transcriptional effects of ROS1-induced DNA hypomethylation from those caused by DNA damage, we compared the upregulated DEGs (‘up’ cluster; Figure 3b) with upregulated DEGs by TEN-ELEVEN TRANSLOCATION 1 (TET1), based on a previous characterisation of *35S:TET1* overexpression lines^33^ (Supplementary Data 2). Since TET1 converts 5mC to oxidised derivatives that are poor substrates for DNA methyltransferases^33,34^, enhanced TET1 activity results in passive loss of DNA methylation without directly inducing DNA damage in plants. Genes uniquely upregulated in E2-treated *XVE:ROS1-YFP* lines were enriched for GO terms linked to DNA damage and chromatin organisation (Figure 3c), whereas the 810 genes commonly upregulated in both E2-treated and *35S:TET1* lines were significantly enriched with genes related to SA-dependent immunity (Figure 3c; Supplementary Data 3). Accordingly, we conclude that the epigenetic DNA demethylation activity of ROS1, rather than its DNA glycosylase/lyase activity, enhances SA-dependent immunity in Arabidopsis.

### Enhanced ROS1 activity induces contrasting patterns of small RNA accumulation between chromosome arms and pericentromeres

To characterise the ROS1-induced changes in sRNA profiles, we mapped sRNA clusters against the Arabidopsis genome using ShortStack^35,36^. E2-treated L5 and L7, but not Col-0, showed thousands of differentially expressed sRNA clusters (DECs) compared to the DMSO-treated controls (Figure 3d; Supplementary Figure 7a). Downregulated DECs primarily occurred at gene-TE boundaries, while upregulated DECs were more abundant at intergenic TEs (Figure 3d). Most of these DECs were composed of 24-nucleotide (nt) sRNAs, which accounted for 97% of downregulated and 82-85% of upregulated DECs in both lines (Supplementary Data 4; Supplementary Figure 7b). Only 17 DECs were identified in E2-treated Col-0, demonstrating a negligible chemical effect of E2 on sRNA expression patterns (Supplementary Figures 3g, 7b; Supplementary Data 4). Plotting the genomic locations of all up- and downregulated DECs across the five nuclear chromosomes revealed a dramatic disparity in their distribution: downregulated DECs were predominantly located along chromosome arms, whereas upregulated DECs mostly mapped to the pericentromeric regions (Figure 3e). This pattern correlates with the spatial distribution of DNA methylation changes in E2-treated *XVE:ROS1-YFP* lines (Figure 3a; Supplementary Figure 7d), highlighting opposing epigenetic dynamics in chromosome arms and pericentromeric regions following transient increases in ROS1 activity.

### Epigenetic *cis*-regulation of immune and DNA damage response genes via ROS1–RdDM antagonism

DNA methylation has been proposed as a *cis*-regulatory mechanism that suppresses defence gene expression in Arabidopsis^5,25^. For example, in *ros1* mutants, hypermethylated cytosines in the promoters of defence genes, such as *RLP43*, can inhibit transcription factor binding and subsequently repress gene activation^25,38,39^. By contrast, plants impaired in RdDM machinery display enhanced transcriptional responsiveness to pathogens^5,39,40^. Therefore, ROS1 and RdDM are thought to act antagonistically to regulate DNA methylation and the expression of environmentally responsive genes^5,24,26^. Consistent with this antagonism between ROS1 and RdDM, hypomethylated DMRs in L5 and L7 show marked decreases in 23- and 24-nt sRNAs (Supplementary Figure 8a).

Therefore, to identify putative ROS1-RdDM *cis*-regulated targets, we performed a genome-wide screen for protein-coding genes with increased expression in E2-treated *XVE:ROS1-YFP* lines (*p_adj_* < 0.05) that also exhibit promoter hypomethylation (hypo-DMR, *p_adj_* < 0.05) and/or downregulation of 24 nt DECs (*p_adj_* < 0.05; Figure 4d). Overlap analysis identified 253 and 271 putative *cis*-target genes in L5 and L7, respectively, with 173 shared between both lines (Figure 4e; Supplementary Data 5). Notably, shared targets included *RLP43*, a previously reported defence gene and *cis*-target of ROS1^25^, supporting the validity of our approach. Furthermore, shared targets were significantly upregulated in the globally hypomethylated *mddcc* mutant (Supplementary Figure 8b), independently supporting a positive association between DNA hypomethylation and transcriptional activation of these genes^41^.

**Figure 4.**
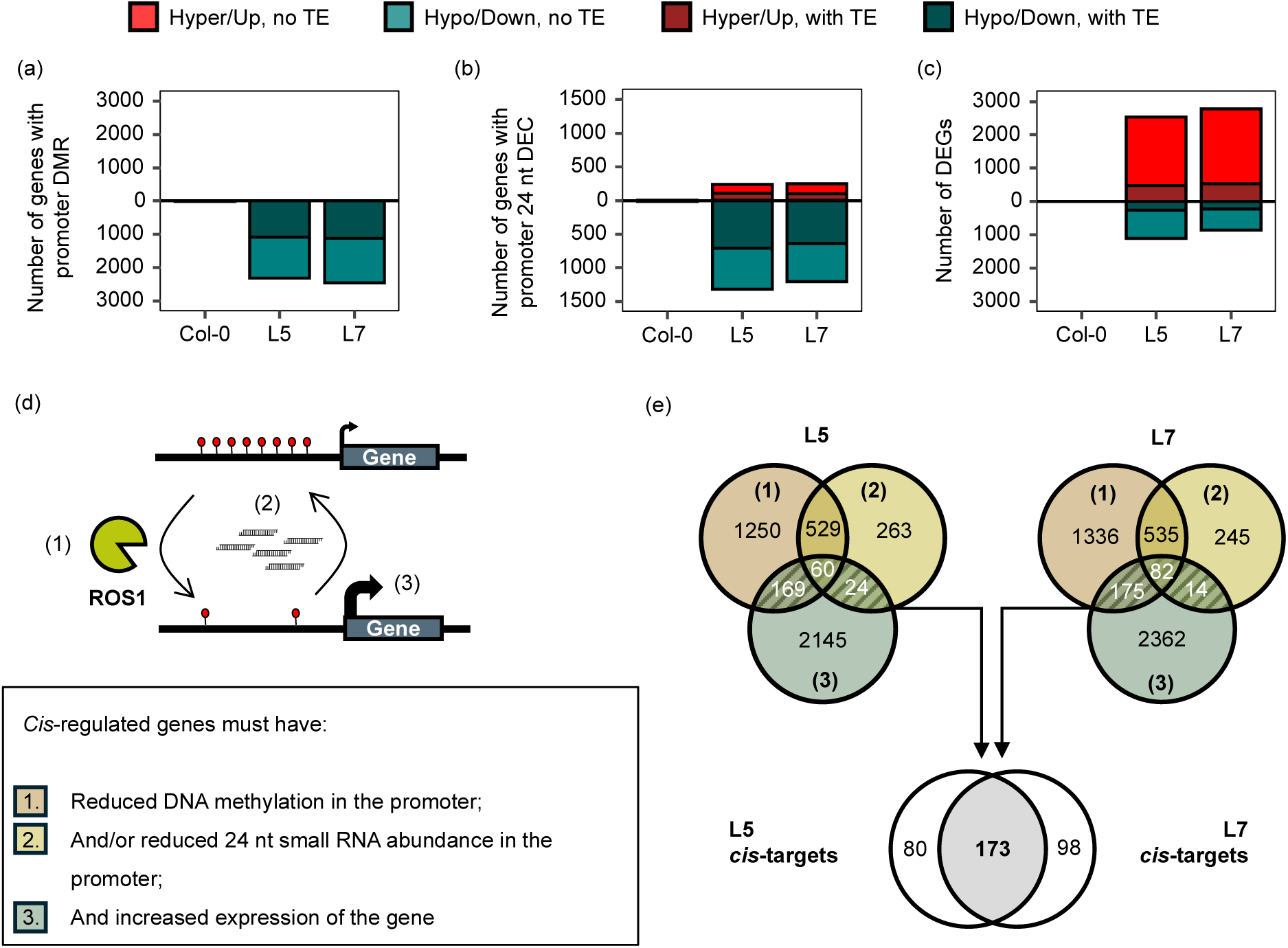
Identification of *cis*-regulatory targets of ROS1. **(a)** Counts of differentially methylated regions (DMRs) within promoter regions (1 kb upstream of TSS or until the next upstream gene, whichever was shorter). Hyper- and hypomethylated DMR counts are shown above and below the y = 0 line, respectively. Dark colours indicate counts overlapping with transposable elements (TE). **(b)** Counts of differentially expressed clusters (DECs) within promoter regions, considering only those classed as 24 nt in size by ShortStack^35,36^. Upregulated and downregulated counts are shown above and below the y = 0 line, respectively. Dark colours indicate counts overlapping with TEs. **(c)** Counts of protein-coding differentially expressed genes (DEGs). Upregulated and downregulated counts are shown above and below the y = 0 line, respectively. Dark colours indicate genes containing one or more TEs within their promoter. **(d)** Schematic demonstrating *cis*-regulatory model under consideration. (1) ROS1 removes DNA methylation from the promoter region of a gene, (2) and/or antagonises RdDM in the region driven by 24 nt sRNAs, (3) which leads to increased transcriptional activity of the gene. Box defines what is considered a *cis*-regulated gene under this model. **(e)** Venn diagram showing numbers of cis-regulated genes by ROS1 in L5 and L7, as defined in (d). Hashed regions represent candidate genes.

In L5, *cis*-regulated candidate targets were enriched for GO terms including ‘DNA damage response’ and ‘response to systemic acquired resistance’, whereas L7 and the shared targets showed enrichment only for ‘DNA damage response’ (*p_adj_* < 0.05; Supplementary Data 3). Notably, *NON-EXPRESSOR OF PATHOGENESIS-RELATED GENES 1* (*NPR1*), which is a master regulator of SA-induced immunity^42^, and *THIOREDOXIN HOMOLOG 5 (TRXH5)*, which promotes the monomerization and nuclear translocation of NPR1^43,44^, were both identified as *cis*-targets in L5 and L7 (Supplementary Figure 9). Likewise, *ATAXIA TELANGIECTASIA AND RAD3-RELATED* (*ATR*) and the transcription factor *E2FC*, which facilitate DNA damage responses following single-stranded breaks^45^, were amongst the consistent targets (Supplementary Data 5). Our data support a model in which ROS1 and RdDM act antagonistically at a subset of loci to regulate key components of plant immunity and genome stability. It is important to note that promoter hypomethylation was not globally associated with increased gene expression (Supplementary Figure 8c), indicating that only a subset of DNA methylation changes *cis-*regulates gene expression. This finding also suggests that most ROS1-responsive DEGs are indirectly regulated, consistent with previous reports^5,25,46,47^.

### Long-read direct DNA sequencing reveals increased non-CG methylation at the (peri)centromeres after ROS1 induction

Our WGBS analysis indicated that pericentromeric regions are less affected by ROS1-induced DNA hypomethylation (Figure 3). Since the reduced DNA methylation by ROS1 anticorrelated with the abundance of sRNAs (Figures 3a, 3e), we hypothesised that the limited hypomethylation in pericentromeric regions is not caused by reduced ROS1 targeting but rather by increased RdDM activity in these regions that counteracts ROS1 activity. However, the highly repetitive nature of pericentromeres limits epigenetic characterisation of these regions, since short-read sequencing approaches map poorly to these regions^48–50^.

To obtain a more reliable analysis of pericentromeric and centromeric (hereafter referred to as (peri)centromeric) DNA methylation, we leveraged long-read sequencing technology and the centromere-complete Col-CEN reference genome assembly^49^. We restricted this analysis to L5, as our previous phenotypic and molecular analyses demonstrated that L5 and L7 show nearly identical responses to E2, while E2 itself had negligible effects on untransformed Col-0 plants (Figures 1-4, Supplementary Figures 1-9). L5 plants were treated twice with either DMSO or E2 (Figure 2a), after which biologically replicated samples (n=3 per treatment) were sequenced using Oxford Nanopore Technology (ONT-seq)^7^. Reads were mapped to the Col-CEN genome assembly and cytosine methylation was called using DeepSignal-plant^51^. Like the WGBS results (Supplementary Figure 3b), PCA of DNA methylation revealed a distinct separation between DMSO- and E2-treated L5 plants (Supplementary Figure 10). Moreover, chromosome-level metaplots of DNA methylation confirmed hypomethylation in chromosome arms, as was also clear from the WGBS data (Figure 5a, Figure 3a). However, the improved resolution by ONT-seq and the Col-CEN genome at (peri)centromeres enabled us to detect a much clearer DNA hypermethylation response in these highly repetitive regions after ROS1 induction (Figure 5a).

**Figure 5.**
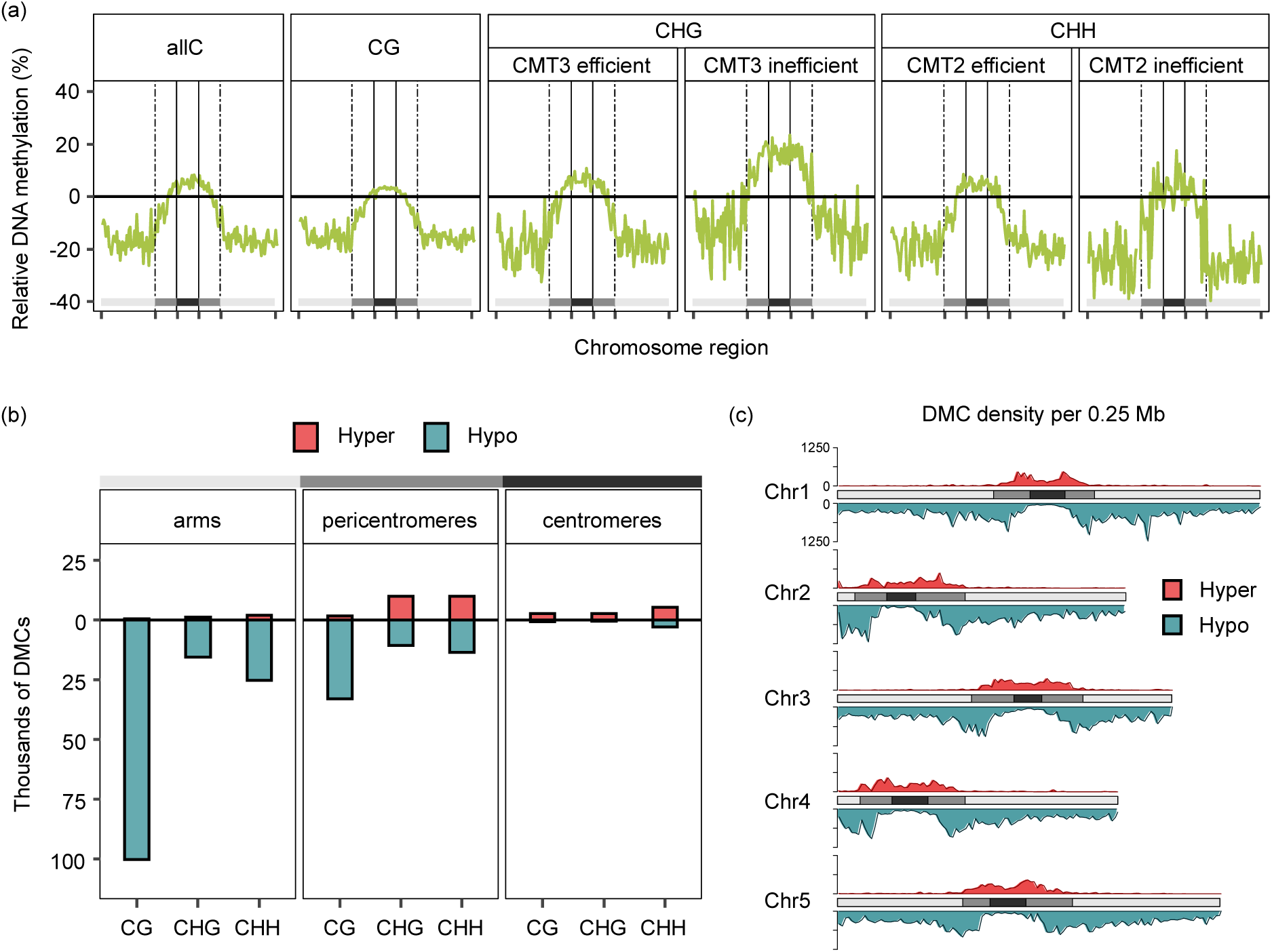
Long-read ONT-sequencing analysis of global DNA methylation responses to induced ROS1 activity. **(a)** Chromosome-level metaplots showing mean percentage differences in cytosine DNA methylation in E2-treated *XVE:ROS1-YFP* L5 plants relative to DMSO-treated plants, determined by long-read Oxford Nanopore Technology sequencing (ONT-seq) and DeepSignal-Plant^51^. For each chromosome in the Col-CEN genome assembly^50^, metaplots were generated by calculating DNA methylation in fixed genomic bins per chromosome: 50 bins for the left arm, 50 bins for the right arm, 20 bins for the centromeric region, and 20 bins for each pericentromeric flank (i.e., 40 bins total for the pericentromeres). DNA methylation was then averaged across biological replicates (n = 3), bins, and chromosomes. For CHG and CHH contexts, DNA methylation was calculated separately for CMT2/3-efficient and -inefficient subcontexts^21^. **(b)** Total counts of differentially methylated cytosines (DMCs; *p* < 0.05, absolute change > 10%)^52,53^ in chromosome arms, pericentromeres, and centromeres. Counts of hyper- and hypo-DMCs are shown above and below the y=0 line, respectively. **(c)** Density of DMCs within 0.25 mega-base (Mb) bins across the 5 chromosomes of the Col-CEN assembly. Light grey, dark grey, and black regions represent chromosome arms, pericentromeres, and centromeres, respectively.

The sequence subcontexts of CHG and CHH methylation sites serve as indicators of the mechanisms driving their establishment and/or maintenance^21^. CHG methylation in the CTG and CAG sub-contexts is typical for CMT3-dependent DNA methylation (‘CMT3 efficient’), while CHH methylation in the CAA and CTA sub-contexts is typical of CMT2-dependent DNA methylation (‘CMT2 efficient’). All other sub-contexts in CHG (CCG) and CHH (CAC, CAT, CCA, CCC, CCT, CTC, CTT) contexts are CMT3- and CMT2-inefficient, and are more likely to be maintained by RdDM^21^. The increased CHH methylation in centromeric and pericentromeric regions did not show a statistical difference between CMT2-efficient and CMT2-inefficient sub-contexts (two-sided Wilcoxon rank-sum test, *p* > 0.05; Figure 5a). However, for CHG contexts, the relative increase in DNA methylation was significantly greater for CMT3-inefficient contexts compared to CMT3-efficient contexts in (peri)centromeric regions (*p* < 0.001; Figure 5a). Hence, the increase in non-CG DNA methylation in the (peri)centromeres is not confined to CMT-dependent sub-contexts, indicating involvement of other pathways, such as RdDM^21^.

To confirm the contrasting patterns of DNA methylation between chromosome arms and (peri)centromeres, we used DSS^52,53^ to select for differentially methylated cytosines (DMCs) between E2- and DMSO-treated L5 plants (*p* < 0.05, absolute change > 10%; Supplementary Data 6). In support of the results depicted in the metaplots (Figure 5a), this analysis identified tens of thousands of hypomethylated DMCs across all contexts (CG, CHG, CHH) in the chromosome arms, with few hypermethylated DMCs. In contrast, several thousand hypermethylated DMCs were detected in the (peri)centromeres, particularly in the CHG and CHH contexts (Figure 5b, 5c). Collectively, our ONT-seq data suggest that transient ROS1 activity triggers a DNA hypermethylation response in (peri)centromeric regions which may act to prevent widespread chromatin de-condensation in these TE-rich regions. Supported by our complementary finding that transient ROS1 activity increases (peri)centromeric sRNA accumulation (Figure 3), we propose that this protective response is driven by RdDM, rather than by CMT2/3-dependent methylation.

### Loss of CLSY3 attenuates ROS1-induced sRNA accumulation in the pericentromeres

In non-stressed vegetative tissues of Arabidopsis, RdDM activity is largely confined to the chromosome arms^18,22^. The genomic distribution of Pol-IV-associated 24 nt sRNAs is coordinated by competing CLSY proteins, which recruit Pol-IV to distinct genomic regions^18,19,54^. CLSY1 predominates in the vegetative tissues, directing Pol-IV RdDM to the chromosome arms, whereas CLSY3/4 are more active in reproductive tissues, directing RdDM towards pericentromeric regions^18,19^. Strikingly, *CLSY3* transcript levels increased approximately sevenfold after E2 induction in *XVE:ROS1-YFP* plants (Figure 6a; Supplementary Data 2), which we verified by RT-qPCR in an independent experiment (Supplementary Figure 11a). Since transient ROS1 activation enhanced sRNA accumulation and DNA methylation in (peri)centromeric regions (Figure 3; Figure 5), we hypothesised that the elevated *CLSY3* expression contributes to a redistribution of RdDM machinery to these regions. To test this, we introduced the previously characterised *clsy3-1* mutation^19^ into our inducible *XVE:ROS1-YFP* line (L5) and compared genome-wide sRNA, DNA methylation, and mRNA profiles between L5 and *clsy3*/L5 plants.

**Figure 6.**
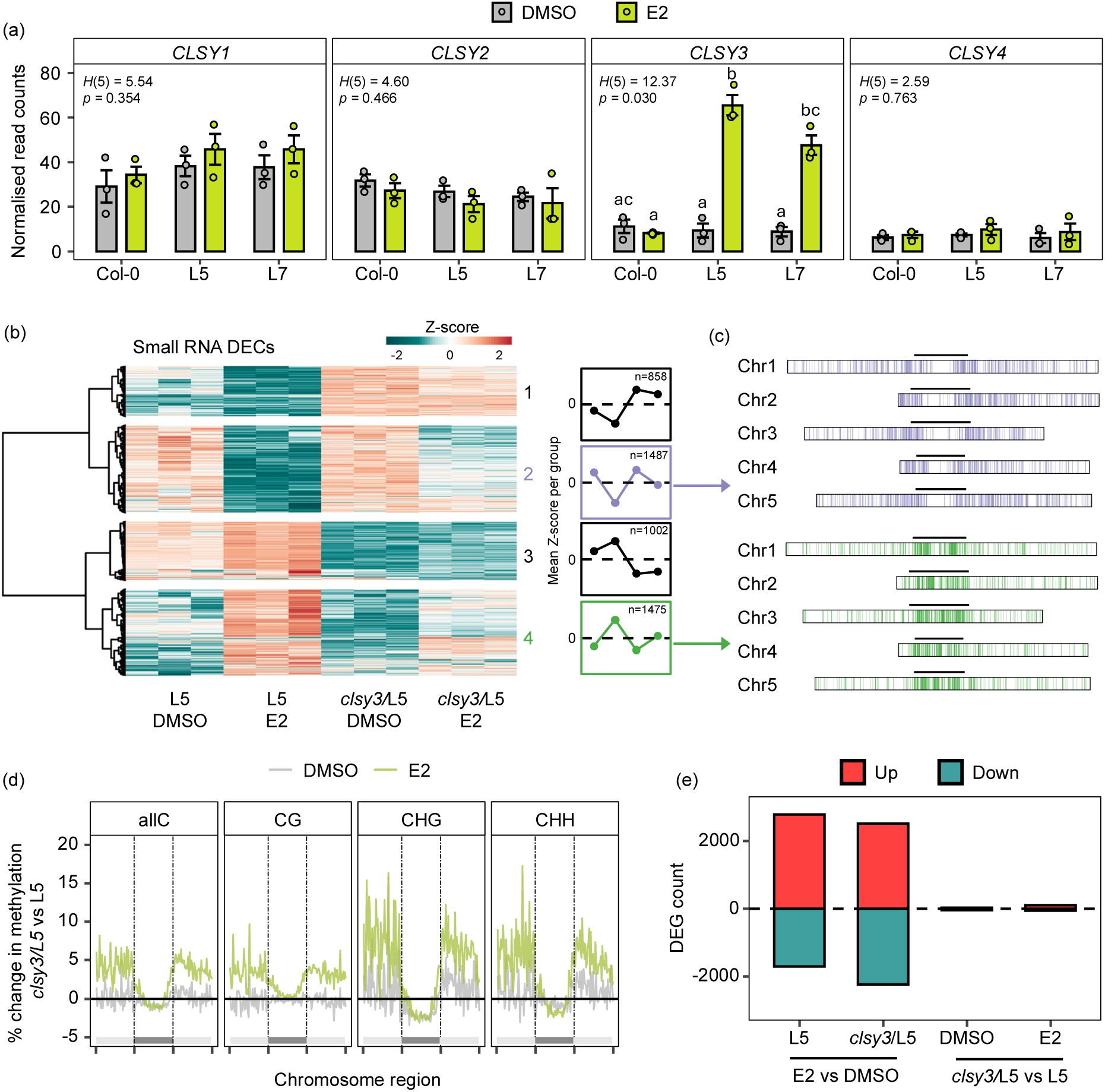
Role of CLSY3 in the epigenomic and transcriptomic response to ROS1 induction. **(a)** DESeq2-normalised read counts of *CLSY* genes in Col-0 and *XVE:ROS1-YFP* lines (L5, L7) 48 hours after treatments with two successive DMSO (0.02% and 0.05%) or E2 (10μM and 25 μM) treatments at 14 and 18 days after sowing (DAS; Figure 2a). For each gene, groups which do not share a letter are statistically different (Kruskal-Wallis *p* < 0.05; Conover-Iman post-hoc *q*-value < 0.05)^55^. **(b)** Heatmap showing Z-scores of normalised and transformed sRNA read counts in all identified DECs between genotypes (*clsy3*/L5 vs L5) and treatments (E2 vs DMSO). DECs were identified using ShortStack^35,36^ and DESeq2^56^. Each genotype-treatment combination is based on 3 biological replicates. Hierarchical clustering was performed for DECs (Euclidean, Ward), which resolved 4 DEC clusters based on expression patterns. Line graphs show mean Z-scores of DECs within each of these four cluster groups. **(c)** Genomic location of DECs in group 2 (purple) and group 4 (green) across the five nuclear chromosomes of TAIR10 Arabidopsis genome assembly. Black lines above box mark pericentromeres. **(d)** Chromosome-level metaplots showing mean percentage difference in DNA methylation in *clsy3*/L5 relative to L5 for DMSO- (grey) and E2-treated (green) plants. Average DNA methylation was calculated for 50 equally sized bins in the left arm, the pericentromeric region, and the right arm before averaging across replicates (n=3) and chromosomes. Dashed lines and dark grey rectangles represent pericentromeres. **(e)** Shown are total number of all DEGs (DESeq2, *p_adj_* < 0.05) identified between treatments (E2 vs DMSO) and between genotypes (*clsy3*/L5 vs L5). Up- and down-regulated DEGs are plotted above and below the y=0 line, respectively.

PCA of sRNA profiles revealed clear global differences between *clsy3*/L5 and L5 under both basal (DMSO) and induced (E2) ROS1 conditions (Supplementary Figure 11c). DECs across all treatment-genotype combinations grouped into four major clusters (Figure 6b). The two largest clusters (groups 2 and 4; n=1,487 and n=1,475) displayed a markedly reduced response to E2 in *clsy3*/L5 relative to L5, despite the slightly higher *ROS1* induction in the mutant background (Figure 6b; Supplementary Figure 11b). Furthermore, groups 2 and 4 showed clear spatial biases: DECs which were less repressed by ROS1 in *clsy3-1* mutants (group 4) were enriched in the pericentromeres, whereas DECs that were less induced by ROS1 in *clsy3-1* mutants were mostly located in chromosome arms (Figure 6c). The two smaller clusters, groups 1 (n=858) and 3 (n=1,002), representing constitutive increases and decreases in sRNAs by *clsy3-1*, irrespective of treatment, were likewise partitioned between chromosome arms and pericentromeric regions, respectively (Supplementary Figure 11f-i).

Consistent with these sRNA patterns, PCA of genome-wide DNA methylation revealed global differences between *clsy3*/L5 and L5 plants (Supplementary Figure 11d). Chromosome-level metaplots indicate that *clsy3*/L5 plants exhibit reduced DNA methylation in the pericentromeric regions and a concomitant increase along chromosome arms, which was more pronounced following E2 treatment (Figure 6d). The modest magnitude of these DNA methylation changes suggests that additional mechanisms alongside CLSY3 participate in the redistribution of DNA methylation activity. Nonetheless, our combined sRNA and methylome data indicate that *CLSY3* contributes to the redistribution of RdDM from chromosome arms to pericentromeric regions after transient increases in ROS1 activity.

Despite the global impacts of the *clsy3-1* mutation on epigenomic responses to ROS1 activity, PCA of the corresponding polyA-enriched RNA-seq (mRNA-seq) data showed no clear separation between L5 and *clsy3*/L5 under either treatment condition (Supplementary Figure 11e). Although E2 treatment induced thousands of DEGs in both genotypes (Fig. 6e; Supplementary Data 2), direct comparisons between *clsy3/L5* and L5 identified only a few DEGs under either basal (n= 80) or induced (n=164) conditions (Figure 6e). These results indicate that, despite its role in shaping genome-wide RdDM dynamics, CLSY3 exerts only minor effects on gene expression in leaf tissues, both in the absence and presence of increased ROS1 activity.

### DNA re-methylation in the chromosome arms resets ROS1-induced immune memory

Transient increases in ROS1 activity induces resistance against (hemi-)biotrophic pathogens, but this resistance is lost between 7- and 14-days post treatment (Figure 2e; Supplementary Figure 2). Since the establishment of this transient immune memory coincided with substantial reductions in DNA methylation and sRNAs along the chromosome arms (Figures 3, 5, 6; Supplementary Figures 4, 7), we next investigated whether the reversal of these epigenetic marks drives the erasure of immune memory.

To address this, we introduced the *XVE:ROS1-YFP* L5 transgene into mutant backgrounds impaired in non-CG DNA methylation and/or sRNA production, and assessed the durability of ROS1-induced immune memory against *Pst-Lux*. In addition to the *clsy3*/L5 line described above, we generated *cmt3-11*/L5 and *nrpe1-11*/L5 crosses. In non-stressed conditions, the *cmt3-11* mutation primarily affects CHG methylation in pericentromeric regions^57,58^, whereas the *nrpe1-11* mutation disrupts RdDM^5,59^, leading to non-CG hypomethylation along the chromosome arms^22,58^ (Supplementary Figure 12). None of these mutations prolonged ROS1-induced immune memory against *Pst-Lux* (Supplementary Figure 13). However, while *cmt3-11* and *clsy3-1* showed no statistically significant effects on *Pst-Lux* colonisation across all timepoints (Tukey *p_adj_* > 0.05), E2-treated *clsy3*/L5 plants showed a modest increase in *Pst-Lux* colonisation at 7 dpt (Welch’s t=2.89, df=51.3, *p*=0.005) and 14 dpt (Welch’s t=2.00, df=60.4, *p*=0.0495) relative to E2-treated L5 plants. This observation is consistent with the attenuated ROS1-induced DNA demethylation in chromosome arms in the *clsy3-1* background (Figure 6d), suggesting a minor contribution of CLSY3 to immune memory maintenance. In contrast, the *nrpe1-11* mutation increased basal resistance at later timepoints (Supplementary Figure 13) consistent with previous reports linking reduced RdDM activity to enhanced immunity^5,40^. This pleiotropic phenotype, however, confounded our assessment of RdDM in the erasure of ROS1-induced immune memory.

To test whether the recovery of DNA methylation at ROS1-demethylated sites (DNA re-methylation) contributes to the erasure of immune memory, we treated L5 seedlings with 5-Azacytidine (5-Aza), a DNA methyltransferase inhibitor that causes genome-wide reductions in DNA methylation^60^ (Supplementary Figure 12). Small droplets of water or 100 μM 5-Aza were applied to the meristems of Col-0 or L5 seedlings starting 7 days before the first DMSO/E2 treatment and ending 1 day before the final DMSO/E2 treatment (Figure 7a). Aside from mild growth reductions, 5-Aza caused no visible phytotoxicity (Supplementary Figure 14). Plants were then challenged with *Pst-Lux* at 1 dpt, 7 dpt, and 14 dpt after the second E2 treatment, corresponding to the establishment, maintenance, and erasure of the immune memory, respectively (Figure 7a). In the absence of ROS1 induction, 5-Aza treatment had a negligible effect on pathogen colonisation, with only a slight increase in resistance in DMSO-treated Col-0 at 1 dpt (Figure 7b). In contrast, combining E2 and 5-Aza in L5 consistently and synergistically reduced *Pst-Lux* colonisation at all time points relative to E2 treatment alone (*p_adj_* < 0.05), including at 14 dpt, when ROS1-induced immune memory had otherwise disappeared (Figure 7, Figure 2e, Supplementary Figure 13). These results indicate that inhibiting DNA methyltransferase activity with 5-Aza slows the re-methylation of DNA in chromosome arms, which is associated with prolonged ROS1-induced immune memory.

**Figure 7.**
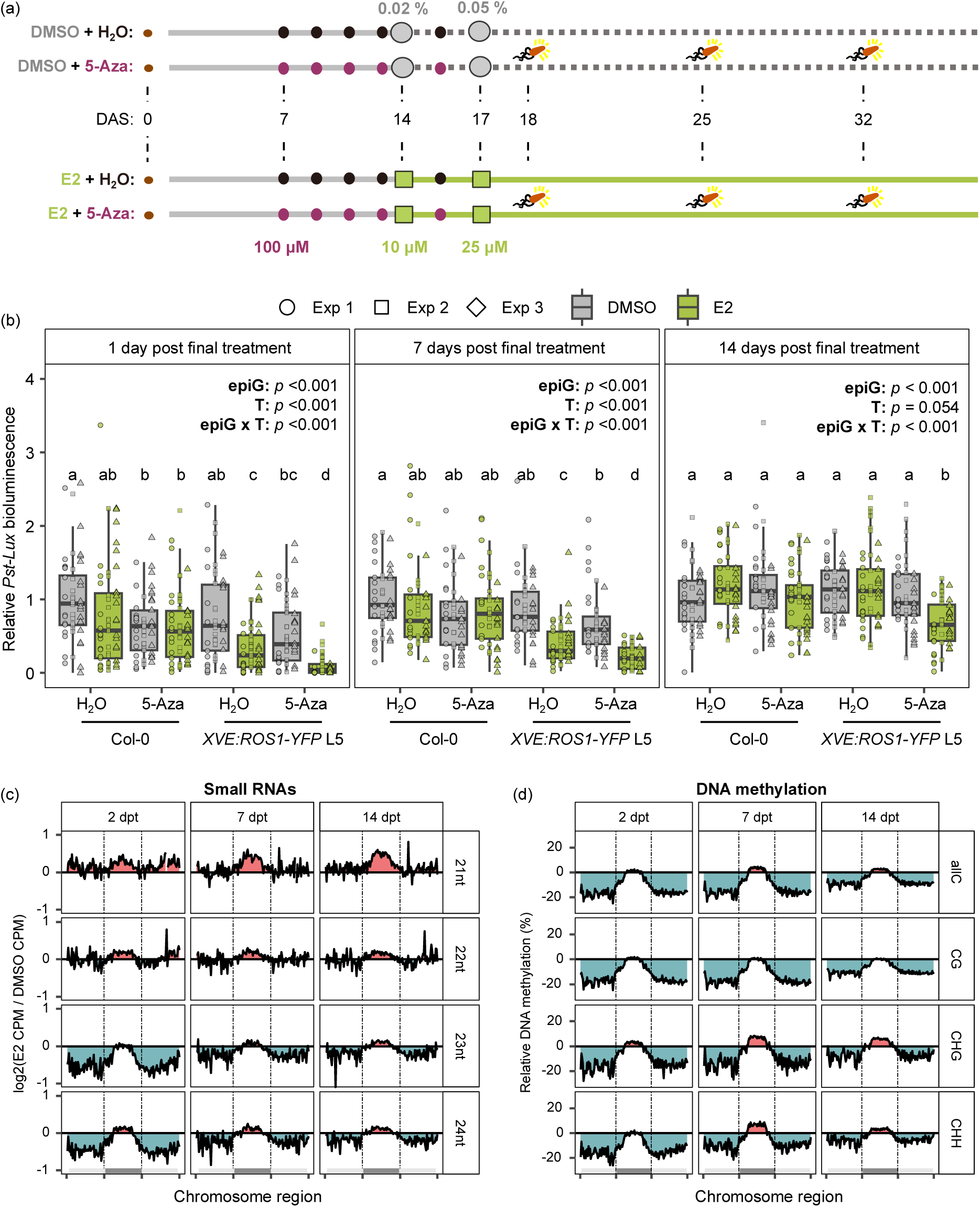
DNA re-methylation of the chromosome arms drives the loss of ROS1-induced immune memory. **(a)** Experimental design to assess the strength and duration of ROS1-induced immune memory against *Pst-Lux* in the presence and absence of 5-Aza. Droplets of 100 μM 5-Aza (purple) or H_2_O (black; mock) were applied 5 successive times at 7, 9, 11, 13, and 16 DAS onto the leaves of Col-0 and *XVE:ROS1-YFP* seedlings (L5). In addition, plants were sprayed twice with DMSO (0.02% and 0.05%) or E2 (10 μM and 25 μM), at 14 and 17 DAS. Plants were challenged with *Pst-Lux* at 1 day (18 DAS), 7 days (25 DAS), or 14 days (32 DAS) after the final E2/DMSO treatment. **(b)** Quantification of *Pst-Lux* bioluminescence at 2-3 days post-inoculation (dpi)^46^. Plants (Col-0, L5, L7) were challenged at 1, 7, or 14 days after the final DMSO (grey) or E2 (green) treatment. For each time point, data from three independent experiments (Exp1, Exp2, Exp3) were normalised to the average *Pst-Lux* colonisation values of H_2_O-DMSO-treated Col-0 plants before pooling. Two-way ANOVA was used to assess the statistical significance of the (epi)genotype (epiG; genotype and 5-Aza/H_2_O treatment), E2 treatment (T), and their interaction (epiG x T) on relative *Pst-Lux* colonisation. For each timepoint, different letters indicate statistically significant differences between treatment-genotype combinations (Tukey, *p_adj_* < 0.05). (**c-d**) Chromosome-level metaplots of (c) small RNAs (sRNAs) and (d) DNA methylation in *XVE:ROS1-YFP* L5 at 2, 7 or 14 days post final treatment (2 dpt, 7 dpt, 14 dpt). E2- and DMSO-treatments were performed as shown in Figure 2a: 10 μM and 25 μM E2, or 0.02% and 0.05% DMSO, at 14 and 18 DAS, respectively. For each chromosome in the TAIR10 genome assembly, (c) sRNA counts, or (d) mean DNA methylation were calculated in 50 bins for each chromosomal region (left arm, pericentromeres, right arm) and then averaged across replicates (n=3 per timepoint) and chromosomes. For each timepoint, plots show (c) mean log_2_ fold-changes in sRNA counts per mapped-million (CPM) for 21-24 nt reads, and (d) mean percentage differences in DNA methylation for each context (allC, CG, CHG, CHH), relative to DMSO-treated controls. Dashed lines indicate pericentromeric regions. Relative increases and decreases have been shaded in red and blue, respectively.

Because the *clsy3-1* and *cmt3-11* mutations, which reduce pericentromeric DNA methylation, did not prolong ROS1-induced immune memory to *Pst-Lux* (Supplementary Figures 11-13), we hypothesised that the erasure of immune memory via DNA re-methylation occurs in the chromosome arms. To confirm this, we performed time-resolved epigenomic profiling, tracking sRNA abundance and global DNA methylation from memory establishment (2 dpt) through maintenance (7 dpt) and erasure (14 dpt). Consistent with a role for RdDM in memory erasure, we observed a gradual recovery of sRNAs and non-CG DNA methylation along the chromosome arms that coincided with the loss of ROS1-induced immune memory (Figures 7c-d). The promoters of our 173 *cis*-regulated gene targets (Figure 4) remained hypomethylated from 2 dpt to 7 dpt but exhibited significant recovery by 14 dpt (Supplementary Figure 15). In contrast, ROS1-induced increases in pericentromeric sRNAs and DNA methylation were maintained or even enhanced over the same period, indicating that this response does not control immune memory (Figures 7c–d). Together, our results demonstrate that ROS1-induced immune memory is epigenetically associated with transient DNA hypomethylation in chromosome arms and is reset through progressive re-methylation in these regions.

## Discussion

Here, we demonstrate that transient increases in ROS1 activity can establish immune memory against (hemi-)biotrophic pathogens in Arabidopsis. Previous research linked loss of ROS1 function to impaired immunity and transcriptional responsiveness^5,7,25^. However, these studies are confounded by pleiotropic effects associated with constitutive loss of ROS1 function in *ros1* mutants and cannot address the role of loci that are already highly methylated under stress-free conditions. Using an inducible *XVE:ROS1-YFP* system, we have overcome these limitations and provide a framework to dissect the mechanisms underlying the establishment, maintenance, and erasure of epigenetic immune memory in plants. We also identify new putative ROS1 targets, including the promoter regions of the immune-regulatory genes *NPR1* and *TRXH5*^42–44^ (Supplementary Figure 9; Supplementary Data 5), which likely contribute to ROS1-induced immune memory (Figures 2 and 7; Supplementary Figures 1 and 13) and transcriptional activation of SA-dependent immunity (Figure 3; Supplementary Figure 6). Although hypomethylated DMRs were significantly enriched for ROS1 occupancy in two independent ChIP-seq datasets (Supplementary Figure 5c-d)^26,32^, direct assessment of ROS1-YFP recruitment at individual loci during induction (Figure 2a) would require time-resolved ChIP-seq analyses in the *XVE:ROS1-YFP* system.

Ectopic induction of ROS1 caused significant reductions in DNA methylation along the chromosome arms (Figure 3; Supplementary Figure 4) but was accompanied by an unexpected increase in DNA methylation across the (peri)centromeric regions that persisted at least fourteen days (Figures 3a, 5a, 7b). This hypermethylation coincided with increased pericentromeric sRNA accumulation (Figures 3e, 7c) and an approximate sevenfold increase in *CLSY3* expression (Figure 6a; Supplementary Figure 11), suggesting that enhanced ROS1 activity drives CLSY3-associated redistribution of RdDM towards the pericentromeres (Figure 6)^18,19,54^. Consistent with this model, ROS1 induction in the *clsy3-1* background (*clsy3*/L5) attenuated these patterns: pericentromeric sRNA accumulation was reduced, and depletion of sRNAs from chromosome arms was less pronounced (Figure 6). Correspondingly, ROS1-induced DNA hypomethylation in chromosome arms and DNA hypermethylation in pericentromeric regions were both reduced in *clsy3*/L5 (Figure 6d). These findings support a model in which CLSY3 competes with other CLSY proteins to guide Pol-IV to pericentromeric regions in response to elevated ROS1 activity^19,54^. Thus, while RdDM in unstressed leaf tissues is largely confined to the chromosome arms^18,22^, its spatial distribution appears more dynamic under periods of extensive DNA demethylation, such as those triggered by biotic stress^6,9^. Under these conditions, increased pericentromeric RdDM activity may help limit widespread TE activation and preserve genome integrity.

As ROS1-induced immune memory only persisted for seven days before fading (Figure 2e, 7b; Supplementary Figure 13), the *XVE:ROS1-YFP* system enabled us to investigate the often-overlooked erasure of immune memory. To address this, we introduced the *XVE:ROS1-YFP* construct into select genetic mutant backgrounds (*clsy3-1*, *cmt3-11*, *nrpe1-11*), and assessed their effects on the establishment, maintenance, and erasure of this memory (Supplementary Figure 13). While the *cmt3-11* mutation, which primarily affects pericentromeric DNA methylation (Supplementary Figure 12), had no detectable effect on the strength or duration of the memory, the RdDM-associated mutations, *nrpe1-11* and *clsy3-1*, had notable effects on the resistance phenotype against *Pst-Lux*. Specifically, the *nrpe1-11* mutation, which reduces non-CG methylation along the chromosome arms (Supplementary Figure 12), resulted in elevated levels of basal resistance to *Pst-Lux* at later timepoints, regardless of ROS1 induction. By contrast, the *clsy3-1* mutation did not enhance nor prolong ROS1-induced immune memory significantly, but it did weakly suppress the resistance from 7 days post treatment (Supplementary Figure 13). Given that *clsy3/*L5 lines displayed reduced loss of DNA methylation along chromosome arms following ROS1 induction (Figure 6), these findings collectively suggest that ROS1-induced DNA demethylation followed by re-methylation of chromosome arms drives the establishment and erasure of immune memory, respectively. Complementary experiments with the DNA methyltransferase inhibitor 5-Aza^60^ supported this interpretation: 5-Aza not only enhanced ROS1-induced immune memory but also extended its duration by at least 7 days (Figure 7b), suggesting that DNA re-methylation drives the erasure of immune memory. Indeed, our time-course experiment revealed the ROS1-induced reduction in DNA methylation and 23-24 nt sRNAs along the chromosome arms gradually recovered in parallel with the erasure of ROS1-induced immune memory (Figure 7c-d; Supplementary Figure 15), whereas pericentromeric regions showed sustained and, in some cases, enhanced accumulation of sRNAs and DNA methylation at later timepoints (Figure 7d). Taken together, our findings indicate that transient ROS1-induced immune memory is established through DNA demethylation within chromosome arms and subsequently erased through re-methylation of these regions.

Collectively, our results uncover roles for DNA methylation in both the establishment and erasure of plant immune memory, providing a plausible molecular mechanism underlying the survival and adaptive plasticity of plants in variably stressful environments^14,15^. Beyond advancing our fundamental understanding of ROS1-mediated DNA demethylation, the *XVE:ROS1-YFP* system and the methodologies developed here offer a framework for the controlled introduction of epigenetic variation in plants. More broadly, this opens new opportunities for epigenetic crop breeding and plant protection technologies.

## Methods

### Plant materials and growth conditions

All experiments were conducted using *Arabidopsis thaliana* (Arabidopsis) lines in the genetic background of Columbia (Col-0). For pot-based experiments, seeds were stratified in 1 mL deionised water (dH_2_O) at 4°C in the dark for 2-4 days. Seeds were then sown in 70 mL pots containing a mix of horticultural sand and peat-based compost (either Levington M3 or Levington F2+S) in a 2:1 ratio. Plants were grown under short day conditions (8.5:15.5 h day:night at 21°C at 45–70% RH; Valoya NS1 LED bulbs providing ∼150 μmol photons m^-2^ s^- 1^). To initiate flowering and the passage of generations, plants were grown in long-day conditions (16:8 h day:night at 21-23°C; 45–70% RH; Sylvania GroLux T8 36W bulbs providing ∼ 150 μmol photons m^-2^ s^-1^). Pots were watered from the bottom 2-3 times a week with reverse osmosis water and trays were frequently rotated within the chamber to minimise positional effects.

For hydroponic experiments (Figure 1c, 1d), seeds were vapour-phased sterilised^61^ before sowing into 12-well plates. For each well, ∼5 seeds were sown in 2 mL of ½ strength Murashige and Skoog (MS) basal salt mixture without vitamins (MS; Sigma-Aldrich M5524) supplemented with 0.5% sucrose, adjusted to pH 5.7. Plates were sealed with micropore tape and stratified at 4°C in the dark for 2-4 days, after which the plates were moved to long-day conditions for germination and growth. MS media was replaced after 7 days of growth.

Generation of *XVE:ROS1-YFP* plasmids and transgenic lines is described in detail in the supplementary methods of Wilkinson et al. (2023). Five independent *XVE:ROS1-YFP* T3 lines were used in this study: L5, L6, L7, L30, and L32. The *clsy3-1* (SALK_040366), *cmt3-11* (SALK_148381), *nrpe1-11* (SALK_029919) mutants used in this study have been characterised previously^5,18,19,24,57–59^. Homozygous genetic crosses between *clsy3-1*, *cmt3-11* or *nrpe1-11* and L5 were confirmed using primers listed in Supplementary Table 1.

### Pathogen cultivation, inoculation, and quantification

All plant inoculations with *Pst-Lux* and *Hpa* were performed within the first 4 hours of the light period to minimise the influence of diurnal variation on plant innate immunity^62^.

*Pseudomonas syringae* pv. *tomato* DC3000 *luxCDABE*^29,30^ (*Pst-Lux*) was stored in 15-25% (v/v) glycerol stocks at -70°C. Overnight cultures (16-20 hours; 28°C; shaking at 200 rpm) were grown in King’s B (KB) solution supplemented with rifampicin (50 μg/mL) and kanamycin (50 μg/mL). Prior to infection, plants were covered with transparent lids to increase relative humidity and promote stomatal opening. *Pst-Lux* cells were isolated by centrifugation (4000 x *g*; 3 minutes), washed (10 mM MgSO_4_), and re-centrifuged (4000 x *g*; 3 minutes) before resuspending in 10 mM MgSO_4_. For inoculation, the cell suspension was adjusted to an optical density at 600 nm (OD_600_) of 0.2 before adding 0.015% v/v silwet L-77 (silwet; LEHLE SEEDS, VIS-30)^30^. For each tray, containing 16-18 pots of plants, 35 mL of inoculum was sprayed onto the surface of leaves, resulting in run-off. Inoculated plants were covered with transparent lids and sealed with parafilm. *Pst-Lux* colonisation was quantified by bioluminescence intensity between 1 and 4 days post infection, using a G:BOX Chemi XRQ gel doc system and ImageJ^30^. Briefly, plants were quenched in darkness for 8 minutes before taking a long-exposure image (4 minutes) in the dark. Bright field images (25 ms exposure, with white light) were used to measure plant area and define plant regions. The mean pixel brightness intensity (‘mean-grey’ value in ImageJ) plus 3x the standard deviation was used to subtract background noise from the bioluminescent images (process > math > subtract). Subsequently, the remaining pixels were used to calculate mean-grey values for each plant-derived region. Thus, the final measurement gives a metric of *Pst-Lux* bioluminescence intensity above background levels, normalised to plant area^30^. Further details can be found in Supplementary Methods.

*Hyaloperonospora arabidopsidis* isolate Waco9 (*Hpa*) was maintained on hypersusceptible *NahG*-expressing Arabidopsis plants of the ecotype *Wassilewskija* (Ws). For inoculations, infected leaves from host plants were washed with dH_2_O, filtered through Miracloth, and quantified using a hemacytometer (Improved Neubauer, Hawksley, UK; Depth 0.1 mm, 1/400 mm2). Spore density was adjusted to 1x10^5^ spores/mL and inoculum sprayed onto the surface of leaves until droplets covered all leaves. Five days after infection, aerial tissue was collected in 100% ethanol. Plant material was then stained with trypan blue, as described by Stassen et al. (2018). Individual leaves were microscopically analysed (Optika Italy Lab-20 binocular stereozoom microscope), based on previously reported hallmarks of *Hpa* colonisation and pathogenesis^31^. Briefly, leaves were assigned to four *Hpa* colonisation classes: I: no hyphal colonisation; II: limited hyphal colonisation with 8 or less conidiophores; III: extensive hyphal colonisation with more than 8 conidiophores; and class IV: extensive hyphal colonisation with conidiophores and oospore(s).

### Chemical treatments

Chemical treatments were performed within the first 4 hours of the light period. 17β-estradiol (E2; Sigma-Aldrich E8875) was dissolved in dimethyl sulfoxide (DMSO; Sigma-Aldrich D4540) to a final stock concentration of 50 mM and stored at -20°C. At the time of treatment, stock aliquots were diluted in dH_2_O to the desired working solution. Mock solutions (DMSO treatments) consisted of dH_2_O plus an equivalent volume of DMSO. In soil-based experiments, E2/DMSO solutions were supplemented with 0.015% (v/v) silwet and sprayed onto the surface of leaves (20 mL per tray; 1.1 – 1.5 mL per pot). Unless stated otherwise, E2-and DMSO-treatment refers to a dual spray treatment of soil-grown plants at 14 and 18 days after sowing (DAS) with 10 μM and 25 μM E2 or 0.02% and 0.05% DMSO (Figure 2a). For hydroponic experiments (Figure 1c, 1d), E2/DMSO was added directly to the wells using stock solutions with no added silwet.

Stock solutions of 5-Azacytidine (5-Aza; Sigma #A2385) were prepared in dH_2_O at a concentration of 24.6 mM (6mg/mL). Working solutions were prepared in ddH_2_O to a final concentration of 100 μM and supplemented with 0.015% (v/v) silwet. Mock treatments consisted of dH_2_O and silwet only. To minimise health risks of spraying 5-Aza solution, a single droplet containing either 5-Aza or mock solution was carefully applied to the apical meristem of every seedling. Droplet volumes were: 6 μL at 7 days after sowing (DAS); 8 μL at 9 and 11 DAS; and 10 μL at 13 and 16 DAS.

### Epifluorescence microscopy

Epi-fluorescence microscopy to quantify accumulation of ROS1-YFP in living plants was performed using a Leica M165 FC fluorescent stereo microscope (Objective: 1x/0.06; ET-GFP filter set: 470/40 nm excitation, 495 nm dichroic and 525/50 nm emission) with a CoolLED pE-300 illumination system, controlled by Leica LAS X software v3.7.4.23463. ROS1-YFP intensity was quantified in ImageJ, using the background-subtracted mean-grey value, normalised by plant area. To account for autofluorescence, ROS1-YFP signal was calculated as a relative fold change compared to DMSO-treated plants.

To validate nuclear localisation of ROS1-YFP signal, leaves were detached from soil-grown plants 24 h after spraying with 25 μM E2 and vacuum infiltrated with DAPI solution (2 μg/mL) for 2-3 minutes. Leaves were mounted on slides and imaged using a Leica DM6B epifluorescence microscope (light source: CoolLED pE-2; Objective: HC PL Fluotar 40x/0.80; ET-DAPI filter set: 350/50 nm excitation, 400 nm dichroic, and 460/50 nm emission; ET-GFP filter set: 470/40 nm excitation, 495 nm dichroic, and 525/50 nm emission) controlled by Leica LAS X v3.7.3.23245 software. Using ImageJ, the brightness and contrast were increased equally for: BF images (20% and 10%, respectively); DAPI images (60% and 70%, respectively); and YFP images (40% and 70%, respectively). DAPI and YFP images were pseudo-coloured cyan and yellow, respectively, using ImageJ.

### Reverse transcriptase-quantitative PCR (RT-qPCR)

For each biological replicate in each experiment (Figure 2e; Supplementary Figure 11a), aerial tissue of ∼10 plants were harvested together, flash frozen in liquid nitrogen, and stored at - 70°C. 100 mg of ground tissue was used for RNA extractions using the RNeasy plant mini kit (Qiagen, #74904) and 800 ng of RNA was used for cDNA synthesis using the ‘Maxima First Strand cDNA Synthesis Kit for RT-qPCR, with dsDNase’ kit (Thermo Scientific, K1672). Diluted cDNA (1:5) was used for RT-qPCR using SsoAdvanced SYBR Green Supermix (BioRad, 1725270), ROS1-specific primers (Table 3.1), and a Rotor-Gene Q (Qiagen) real-time PCR cycler. Take off values and primer efficiencies were calculated using the Rotor-Gene Q Series software (v2.1.0). Relative expression values were calculated as described in Wilkinson et al. (2023), using *MON1* (AT2G28390) and *UBC21* (AT5G25760) as stably-expressed reference genes^63^ and a DMSO-treated sample as the calibrator.

### Next-generation sequencing: material harvest, extractions, and sequencing

For short-read sequencing experiments (Figures 3, 6), 48 hours after treating Col-0, L5 and/or L7 twice with E2 or DMSO (Figure 2a), aerial tissues were collected from ∼6 pots in a single tray to form a single biological replicate (n=3). Material was snap frozen in liquid nitrogen and ground to a fine powder using liquid nitrogen and a mortar and pestle. Several ∼100mg aliquots of tissue were stored at -70°C for each biological replicate. Individual aliquots, derived from the same pool of ground tissue, were used for genomic DNA and total RNA extractions, using the DNeasy Plant Mini Kit (Qiagen 69104), and TRIzol Reagent (Thermo Fisher Scientific 15596026), respectively. Genomic DNA was used for whole-genome bisulfite sequencing (WGBS); RNA was used for polyA-enriched RNA-sequencing (mRNA-seq), ribosome-depleted RNA-sequencing (RNA-seq), and/or small RNA sequencing (sRNA-seq; 18-30 nt). Library preparation and sequencing were performed by BGI Genomics using the DNA nanoball (DNB) sequencing (DNBSEQ^TM^) G400 platform.

For long-read DNA methylation sequencing data of E2- and DMSO-treated *XVE:ROS1-YFP* L5 (as depicted in Figure 2a), we employed Oxford Nanopore Sequencing Technology (ONT-seq), as described by Wilkinson et al. (2023). Briefly, for each biological replicate (n=3) per treatment and at 20 DAS (2 days after the final E2 treatment), ∼100 plants derived from 1 tray (∼500 mg fresh weight) were snap frozen in liquid nitrogen and ground to a fine powder using a pestle and mortar. DNA was extracted using the NucleoBond® HMW DNA kit (Macherey Nagel, 740160.20) and library preparation was done using the ONT Rapid Barcoding kit (ONT, SQK-RBK004). Sequencing was performed on FLO-MIN106 flow cells, in combination with the Flow Cell Wash Kit (ONT, EXP-WSH004). Summary statistics for all sequencing data are provided in Supplementary Data 7.

### Next-generation sequencing analysis: alignment and quantification

Reads from WGBS, RNA-seq, mRNA-seq, and sRNA-seq were analysed for adapters and read quality using FastQC v0.12.1 and MultiQC v1.21^64^. For WGBS data, reads were aligned to the TAIR10 genome and called for cytosine methylation using Bismark v0.24.1^65^. DMCs and DMRs were called using the R package DSS v2.46.0^52,53^. Transcriptomic sequencing (RNA-and mRNA-seq) data were aligned and read counts per feature were quantified using STAR v2.7.9a^66^, the TAIR10 genome, and a custom annotation file that merged the Araport11 annotation^37^ and a gene-like annotation file of TEs^67^. Library normalisation and DEG calling were performed using the R package DEseq2 v1.42.1^56^. Due to a confounding effect related to the default normalisation technique, the counts were normalised using a suite of extremely stably expressed genes in Arabidopsis^63,68^, as recommended by the DESeq2 user guide (v1.42.0) and Love et al. (2014) (Supplementary Methods). sRNA reads with perfect matches to known ribosomal RNA sequences were removed using Bowtie v1.3.1^69^ and the RNAcentral database of non-coding RNA sequences (accessed: 23/01/2024)^70^. Filtered reads were then mapped to the TAIR10 genome and *de novo* clusters defined using ShortStack v.4.0.3^35,36^. Differentially expressed clusters were called using the R package DEseq2 v1.42.1.

ONT-seq reads were base called and filtered using Guppy v6.5.7 and the ‘dna_r9.4.1_450bps_hac.cfg’ model. Methylated cytosines were identified using the ‘model.dp2.CNN.arabnrice2-1_120m_R9.4plus_tem.bn13_sn16.both_bilstm.epoch6.ckpt’ model from DeepSignal-plant v0.1.4^51^ in conjunction with Tombo v1.5.1 and the reference genome Col-CEN v1.2^49^. DMCs were called using the R package DSS v2.46.0. Further information on the parameters used are provided in the Supplementary Methods. Summary statistics are provided in Supplementary Data 7.

### Next-generation sequencing analysis: annotation

For all short-read sequencing data (WGBS, RNA-seq, mRNA-seq, sRNA-seq), features were annotated using a custom annotation file that merged the Araport11 annotation^37^ and a gene-like annotation file of TEs^67^ in the TAIR10 assembly (Supplementary Methods). For DMRs and DECs, overlaps with features were identified using the R packages Genomation v1.34.0^71^ and GenomicRanges v1.54.1^72^. Promoters were defined as -1kb from the TSS, or to the next nearest gene, whichever was shorter. Downstream regions were defined as +1kb from the transcriptional termination site. In cases where a DMR or DEC overlapped more than one feature, a single annotation was chosen based on a hierarchy (Supplementary Methods). For each nuclear chromosome in the TAIR10 assembly, the start of the pericentromere was defined as the midpoint of the first 100 kb bin in a string of at least 5 neighbouring 100 kb bins for which TE density > gene density. The end of the pericentromere was defined as the midpoint of the last bin in a string of at least 5 neighbouring bins where TE density > protein-coding gene density (Chr1: 12450000-17650000, Chr2:1250000-7050000, Chr3: 10450000-16250000, Chr4: 1650000-6350000, Chr5: 9750000-14650000). Genome bins were created using bedtools v2.31.0^73^.

For long-read ONT-seq data, the Araport11-based annotation file for the Col-CEN assembly was used in combination with the intact complete EDTA repeat annotation file (https://github.com/schatzlab/Col-CEN/tree/main/v1.2; accessed 21/09/2023)^49^. Pericentromeres were defined as done for the TAIR10 assembly (Chr1: 12050000-19850000, Chr2: 1350000-9850000, Chr3: 10350000-18950000, Chr4: 1750000-9850000, Chr5: 9650000-18150000), but centromere coordinates were used as defined by Naish et al (2021) (Chr1: 14840000-17560000, Chr2: 3823000-6046000, Chr3: 13597000-15734000, Chr4: 4204000-6978000, Chr5: 11784000-14560000).

### Next-generation sequencing analysis: gene ontology enrichment

Enrichment of biological process gene ontology (GO) terms was performed using R packages clusterProfiler v4.14.6^74^ (function: ‘enrichGO’; options: ‘pvalueCutoff = 0.05’, ‘qvalueCutoff = 0.05’, ‘OrgDb = “org.At.tair.db”’, ‘keyType = “TAIR”’, ‘ont=”BP”’, ‘minGSSize = 10’, ‘maxGSSize = 500’, ‘universe = all_proteinGenes_≥10counts_≥6samples’) and org.At.tair.db v3.20.0. GO terms were classed as enriched if they had *p_adj_*-values < 0.05 (Benjamini and Hochberg correction). GO semantic similarity analysis was performed using the R package GOSemSim v2.28.1^75^ (function: ‘godata’; options: ‘OrgDb = “org.At.tair.db”’, ‘ont = “BP”’). The object resulting from this analysis was used in conjunction with clusterProfiler to create a simplified enriched GO list (function: ‘simplify’; options: ‘cutoff=0.7’, ‘semData=SemSimAnalysis_object’, ‘by=”p.adjust”’, ‘select_fun=min’). All GO terms, raw and simplified, are provided in Supplementary Data 3.

### Next-generation sequencing analysis: genome-wide scan of *cis*-regulated targets

*Cis*-regulated targets in this study refer to protein-coding genes whose expression is negatively regulated by DNA methylation in the promoter region, as previously described for other ROS1 targets^24,25^. Given that ROS1 targets are often antagonistically regulated by Pol-IV-dependent RdDM, which requires 24 nt sRNAs and existing methylated cytosines (Supplementary Figure 8a)^16,17,20,22,24,26^, *cis*-regulated genes were defined as protein-coding genes that were significantly upregulated in E2-treated plants, and contained a hypomethylated DMR, and/or a downregulated 24 nt DEC in their promoter region.

Expression of *cis*-regulated targets in *mddcc* and Col-0 plants (Supplementary Figure 8b) were analysed using FPKM values provided by He et al., (2022)^41^. Of our 173 *cis*-regulated targets, five hypothetical-protein genes did not have reported FPKM values in this study (*AT1G28005, AT2G14825, AT3G29645, AT4G07995, AT5G23115*).

### Statistical analyses and generation of plots

All specific statistical tests have been explicitly mentioned in the main text or figure legends. Significant effects identified by one- or two-way ANOVAs were followed by a Tukey’s post-hoc test; where assumptions of normality and/or homoscedasticity were not met, a Kruskal-Wallis test followed by Conover-Iman^55^ post-hoc comparisons was used. Unless otherwise specified, plots were generated using the R package ggplot2 v3.5.0 and statistical analyses were performed using base R v4.3.2 or the R package rstatix v0.7.2. Plots of DECs along chromosomes were generated using the R package ‘KaryoploteR’ v1.28.0^76^.

### Data availability

ChIP-seq data for Myc-ROS1 was analysed using processed data uploaded to the NCBI Gene Expression Omnibus (samples GSM7852288, GSM7852285, GSM7852318, GSM7852316). Raw data for one biological replicate per treatment has already been published for the ONT-seq^7^ and uploaded to GEO (GSM6463424; GSM6463425). The additional ONT-seq samples are newly available under accession GSE304524. WGBS data of *nrpe1-11* and *cmt3-11* mutants were reanalysed from Wilkinson et al (2023)^7^ (GSE199625) and Briffa et al (2023)^77^ (GSE204837; generation 0 data), respectively. WGBS of 5-Aza-treated Arabidopsis are from Griffin et al (2016)^60^ (GSE80300) and RNA-seq of *35S:TET1* lines are from Ji et al (2018)^33^ (GSE93024). All generated and processed data for RNA-seq, sRNA-seq, and WGBS have been deposited to GEO under accessions GSE304523, GSE304519, and GSE304520, respectively. Source data are provided with this paper. While this manuscript is under review, the confidential data on GEO can be accessed by reviewers using the following tokens for ONT-seq, RNA-seq, sRNA-seq, and WGBS. A preprint of this work is available on bioRxiv: https://doi.org/10.1101/2025.08.19.671022.

## Acknowledgements

We thank H. Hipperson for conducting the ONT-seq, D. Pascual-Pardo for horticultural support, S. Shahid for advice on sRNA-seq analyses, and D. Baulcombe for advice on cytosine sub-context methylation analyses. This work was supported by a European Research Council (ERC) Advanced Grant to JT and SAR (“PlantMemo”; 101199639) and two grants from UK Research and Innovation (UKRI): a Biotechnology and Biological Sciences Research Council (BBSRC) White Rose DTP Studentship (BB/T007222/1: project 2449363), and a BBSRC Industrial Partnership Award to JT, SAR, and LMS (BB/W015250/1).

## Author contributions

A.H.P. and J.T. conceptualised the study, with supervision from S.A.R., L.M.S., S.W.W., and J.T. Funding was acquired by J.T., L.M.S., and S.A.R.. A.H.P. conducted the experiments and collected data, with assistance from P.Z., K.Y.M., and L.T. Data analysis and visualisation were performed by A.H.P. with guidance from S.W.W., S.A.R., L.M.S., and J.T.. A.H.P. and J.T. wrote the original manuscript, with input from all authors.

