## Supplementary Information and Data for "Contrasting epigenetic dynamics in chromosome arms and pericentromeres during ROS1-induced plant immune memory": HannanParker_etal_2026_BioRxiv_SuppInfo_R2.pdf

### Supplementary figures

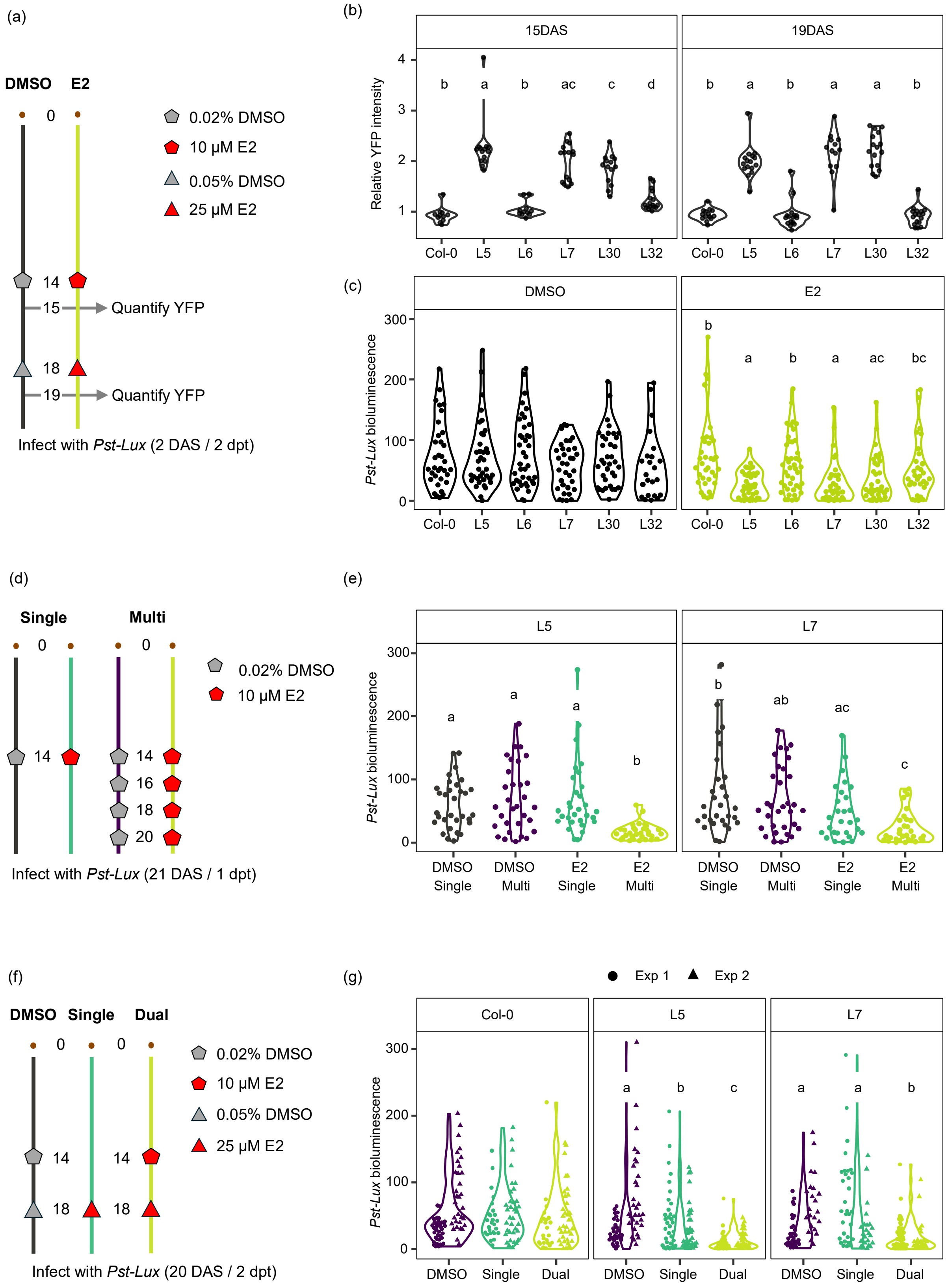


**Supplementary Figure 1. Optimisation and characterisation of 17β-estradiol (E2)-induced immune memory in five independent *XVE:ROS1-YFP* lines (L5, L6, L7, L30, and L32).** Immune memory was determined by the level of E2-induced resistance against *Pseudomonas syringae* pv. *tomato* DC3000 *luxCDABE* (*Pst-Lux*). Bacterial colonisation was quantified by normalised bacterial bioluminescence intensity values[1]. **(a)** Five *XVE:ROS1-YFP* lines and untransformed wild-type plants (Col-0) were treated twice, first at 14 days after sowing (DAS) with 10 µM E2 or 0.02% DMSO, and again at 18 DAS with 25 µM E2 or 0.05% DMSO. Plants were then infected with *Pst-Lux* at 20 DAS, corresponding to 2 days post final treatment (2 dpt). **(b)** YFP fluorescence intensity in whole plants was measured 24 hours after each E2 treatment (15 and 19 DAS; n=7-17). Shown are YFP intensities normalised to the mean DMSO-treated YFP intensity within that group. Statistical analysis revealed significant effects of genotype on relative YFP fluorescence at both 15 DAS (Kruskal-Wallis: H=63.8, df=5, $p < 0.001$) and 19 DAS (H=68.3, df=5, $p < 0.001$). Groups that do not share the same letter are significantly different from each other, as determined by Conover-Iman post-hoc tests with FDR correction ($q < 0.05$). **(c)** At 20 DAS the same group of plants from (a) were inoculated with *Pst-Lux*, and colonisation was measured 3 days post infection (dpi)[1]. Genotype had a statistically significant effect on *Pst-Lux* colonisation in E2-treated plants (H=32.7, df=5, $p < 0.01$) but not in DMSO-treated plants (H=4.48, df=5, $p > 0.05$). For E2 treatment, genotypes with different letters indicate statistically significant differences (Conover-Iman, $q < 0.05$)[2]. **(d-e)** L5 and L7 plants were treated once at 14 DAS ('Single') or four times between 14 and 20 DAS ('Multi') with 10 µM E2. Corresponding controls for each treatment regime were treated with 0.02% DMSO. At 21 DAS, plants were spray-inoculated with *Pst-Lux*, and colonisation was quantified at 3 dpi. Treatment has a statistically significant effect on *Pst-Lux* colonisation in both L5 (ANOVA: F=13.7, df=3,120, $p < 0.001$) and L7 (ANOVA: F=10.2, df=3,118, $p < 0.001$). **(f-g)** Col-0, L5, and L7 plants were treated once with 25 µM E2 at 18 DAS ('Single') or twice at 14 DAS and 18 DAS with 10 µM E2 and 25 µM E2, respectively. Control plants were treated twice at 14 DAS and 18 DAS with 0.02% and 0.05% DMSO, respectively. While treatment had no statistically significant effect on *Pst-Lux* colonisation in Col-0 (F=1.53, df=2,183, $p > 0.05$), it had a significant effect on *Pst-Lux* colonisation in both L5 (F=25.8, df=2,194, $p < 0.001$) and L7 (F=17.4, df=2,175, $p < 0.001$). The experiment was repeated twice, yielding similar results (experiment x treatment x genotype interaction was not statistically significant; F=1.35, df=4,543, $p > 0.05$). For (e) and (g), different letters indicate statistically significant differences between treatments within each line (Tukey post-hoc, $p_{adj} < 0.05$). All chemical treatments were prepared in distilled water, supplemented with 0.015% v/v silwet L-77, and sprayed onto foliar tissue until saturation.

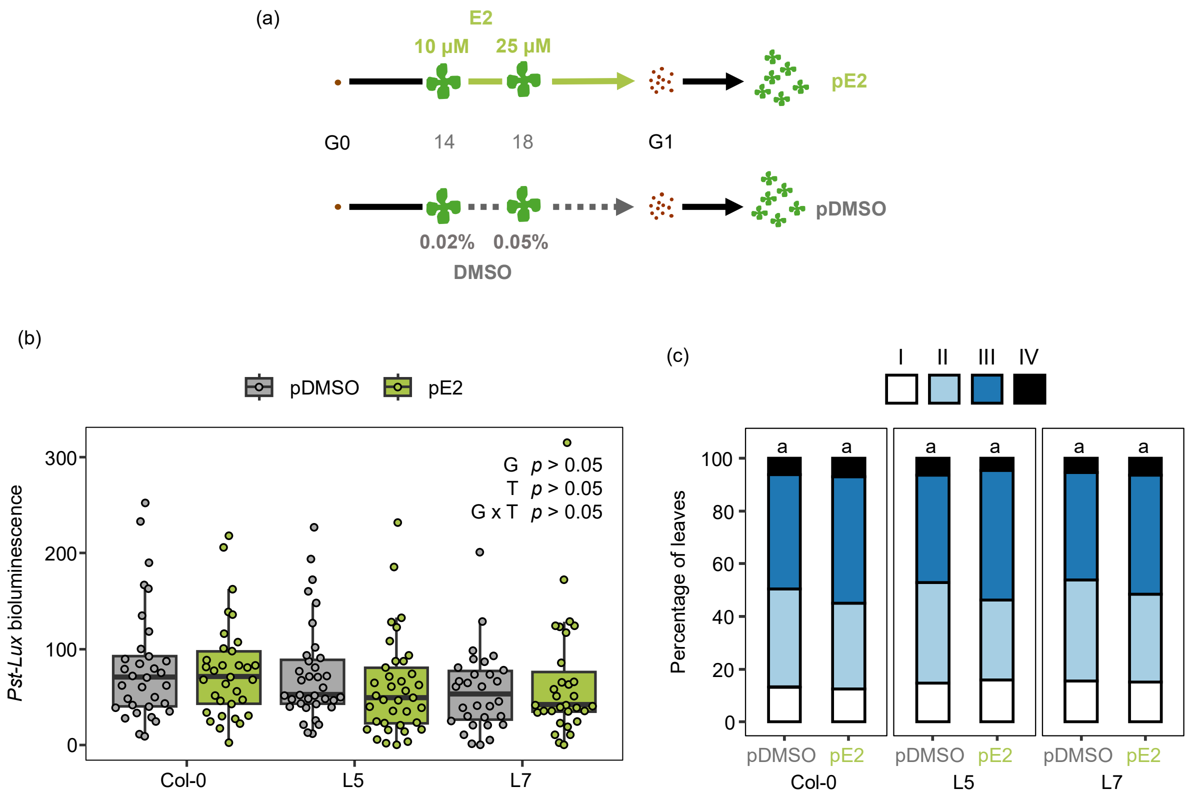


**Supplementary Figure 2. ROS1 induction does not result in heritable resistance against (hemi)biotrophic pathogens. (a)** Col-0, L5, and L7 plants (generation 0; G0) were treated twice with E2 or DMSO at 14 and 18 days after sowing (DAS), as described in the legend of Figure 2. Seeds (G1) were pooled from three individual plants per genotype for each treatment and sown before infecting with *Pst-Lux* (b) at 21 DAS or *Hpa* at 28 DAS (c). (**b**) *Pst-Lux* colonisation was quantified 2 days post infection (dpi) by bacterial bioluminescence[1]. Two-way ANOVA revealed no significant effects of genotype (G) (F=3.04, df=2,193, $p$=0.051), parental treatment (T) (F=0.596, df=1,193, $p$=0.441), or G x T interaction (F=0.997, df=2,193, $p$=0.371) on *Pst-Lux* colonisation. **(c)** *Hpa* colonisation was quantified at 5 dpi by scoring trypan blue-stained leaves. Individual leaves were microscopically assigned to four *Hpa* colonisation classes, ranging from severe (IV) to absent (I)[3]. Shown are relative leaf proportions (%) across the four *Hpa* colonisation classes, pooled by parental treatment and genotype (n = 123–132). Shared letters indicate statistically similar proportions (Fisher's exact test; FDR-correction; $q$ > 0.05).

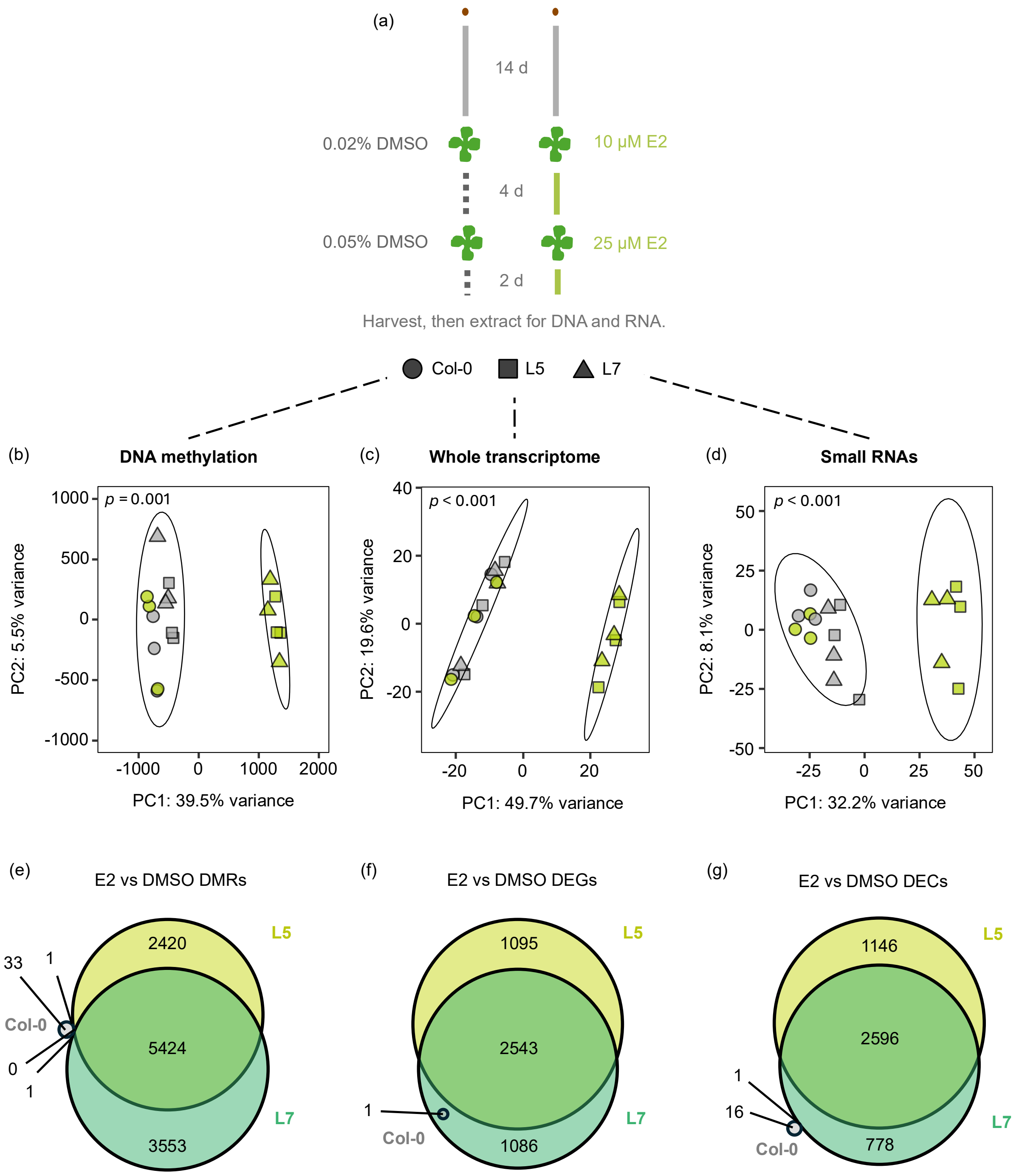


**Supplementary Figure 3. Global epigenomic and transcriptomic patterns in DMSO/E2-treated Col-0 and *XVE:ROS1-YFP* lines (a)** Experimental setup to profile the epigenome and transcriptome of IR by ectopically induced ROS1 (see also Figure 2). Col-0, L5, and L7 plants were treated twice with E2 (green) or DMSO (grey) at 14 days after sowing (DAS; 10 µM E2 or 0.02% DMSO) and 18 DAS (25 µM E2 or 0.05% DMSO). Three biologically replicated samples per genotype-treatment combination from aerial tissues were collected for DNA and RNA extractions at 48 hours after the second chemical treatment (20 DAS). **(b)** PCA plot displaying variation in global cytosine DNA methylation, based on 200 bp weighted-average windows[4], for E2 (green)- and DMSO (grey)-treated Col-0 (circle), L5 (square), and L7 (triangle) plants. **(c, d)** PCA plots displaying global variation in normalised transcript abundance of coding and non-coding RNA transcripts (c; whole transcriptome) and normalised abundance of small RNA clusters (d; small RNAs) in Col-0 (circle), L5 (square), and L7 (triangle) after treatment with either DMSO (grey) or E2 (green). Ellipses within PCA plots represent confidence intervals (95%) for ROS1-induced samples (E2-treated L5/L7) and non-induced samples (E2-treated Col-0 and DMSO-treated L5/L7). *P*-values indicate statistical significance between groups (PERMANOVA), using Euclidean distances and 999 permutations. **(e-g)** Venn diagrams showing the overlap of (e) context-merged differentially methylated regions (DMRs), (f) differentially expressed (non)coding genes (DEGs), and (g) differentially expressed sRNA clusters.

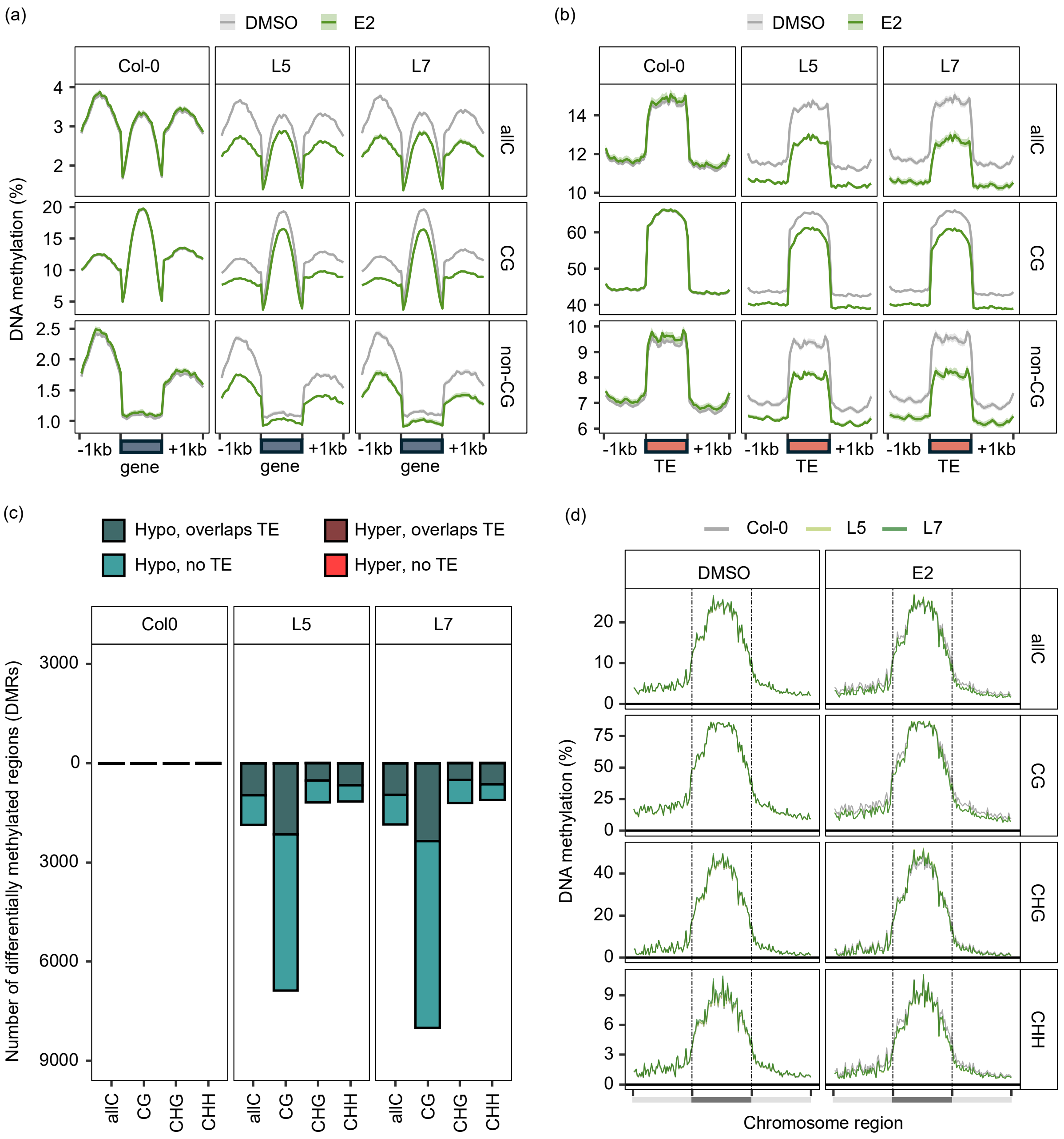


**Supplementary Figure 4. Global DNA methylation patterns after enhanced DNA demethylase activity by ROS1. (a-b)** Metaplots showing DNA methylation (%) in E2/DMSO-treated Col-0 and *XVE:ROS1-YFP* lines (L5 and L7) across allC-, CG-, and non-CG- (CHH/CHG) sequence contexts within (**a**) protein-coding genes ('gene') and (**b**) transposable elements ('TE'). For each gene or TE, DNA methylation was quantified in 20 equally sized bins across the feature body, as well as 20 bins each for the 1 kb regions upstream (-1 kb) and downstream (+1 kb) of the feature. **(c)** Differentially methylated regions (DMRs) between DMSO- and E2-treated Col-0, L5, and L7 for allC, CG, CHG, and CHH sequence contexts. Hypomethylated DMR (Hypo) counts are shown below the y=0 line, hypermethylated DMR (Hyper) counts are shown above the y=0 line. Counts of DMRs overlapping TEs are shown in dark-shaded colours. **(d)** Chromosome-level metaplots of DNA methylation in E2-treated and DMSO-treated Col-0, L5, and L7 plants across all cytosine contexts. For each chromosome, average DNA methylation was calculated for 50 equally sized bins within each of the left arm, the pericentromeric region, and right arm, and then averaged across replicates (n=3) and chromosomes. Dashed lines and dark grey bars represent pericentromeric regions of the genome.

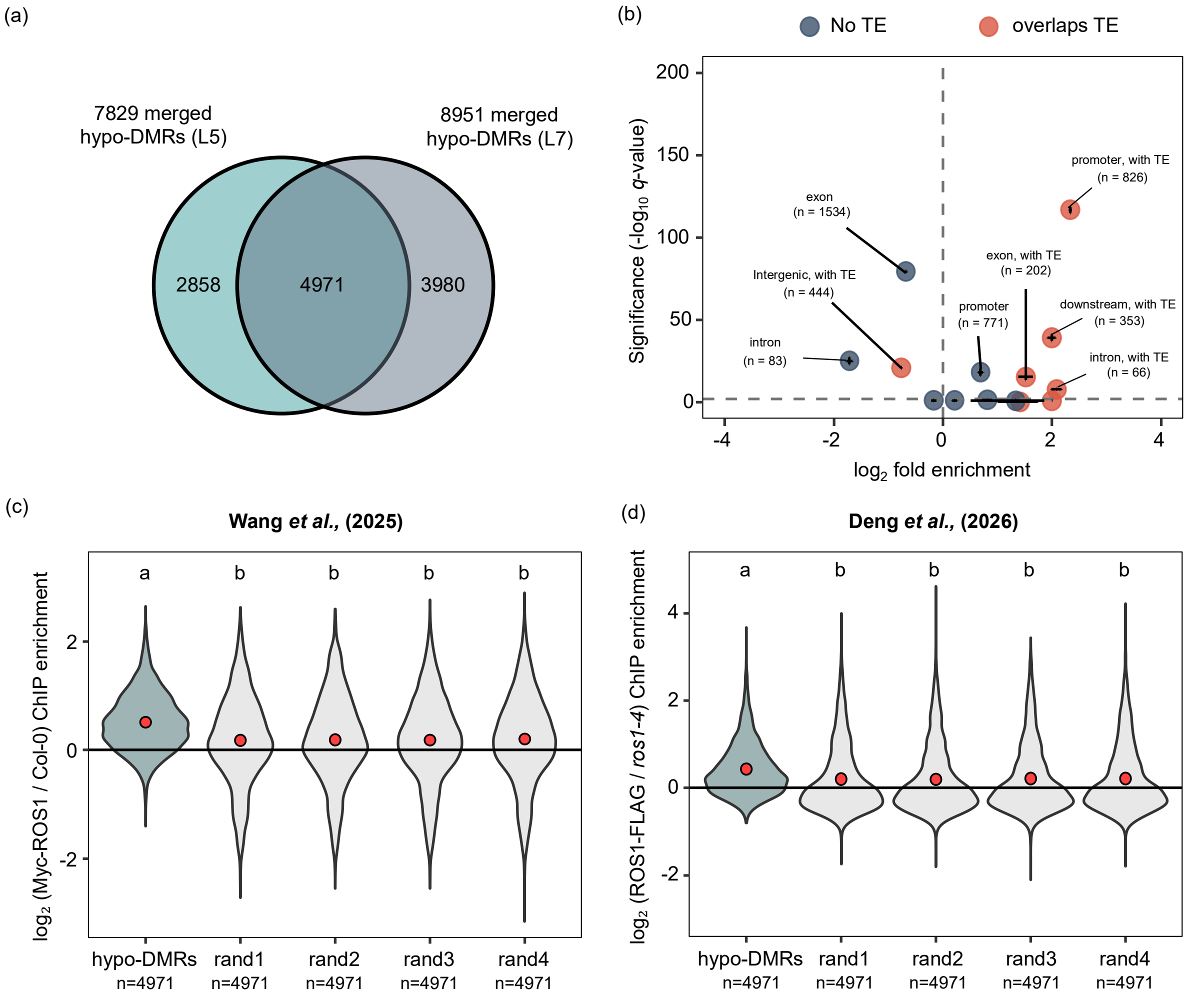


**Supplementary Figure 5. Genomic characterisation of *XVE:ROS1-YFP* hypomethylated DMRs and their overlap with endogenous ROS1 ChIP-seq signal. (a)** Hypomethylated DMRs (hypo-DMRs) from all cytosine context analyses were merged and simplified for L5 and L7. A total of 4,971 merged hypo-DMRs were shared between L5 and L7 ('consistent hypo-DMRs'). **(b)** Enrichment plot of consistent hypo-DMRs across different genomic features. DMRs were annotated with Araport11 [5] for overlap with exons, introns, promoters, downstream regions, non-coding RNA, intergenic regions. Overlaps with TEs are colour-coded (orange). To assess statistical enrichment of consistent hypo-DMRs with genomic features, four sets of randomly selected genomic regions of equal size and number were generated and annotated for comparison. Two-sided Fisher Exact Test's on the number of DMRs matching a given annotation feature were performed between selections of consistent hypo-DMRs and one of the four random sets. The average FDR-adjusted $p$-value ($q$-value) for these tests were plotted against the average fold change (real count / random count) on a $-\log_{10}$ and $\log_2$ scale, respectively. Error bars indicate the standard error of the mean, based on comparisons with four random DMR sets. The horizontal dashed line shows $q = 0.05$. Significantly under- or over-represented annotation features are labelled with the number of consistent hypo-DMRs mapping to the given annotation feature. **(c-d)** Quantification of endogenous ROS1 occupancy from two publicly-available independent ChIP-seq experiments[6,7] within our 4,971 ROS1-induced consistent hypo-DMRs (a). Shown are $\log_2$ fold-enrichment values of (c) Myc-ROS1 ChIP-seq signal relative to the negative Col-0 control or (d) ROS1-FLAG relative to the negative *ros1-4* control. Enrichment was also calculated for the four sets of randomly selected genomic regions (rand1-4) that had been used for the enrichment plot (b). Different letters indicate statistically significant differences between regions (Kruskal-Wallis, $p < 0.001$; Conover-Iman, $q < 0.05$). Red points indicate group means.

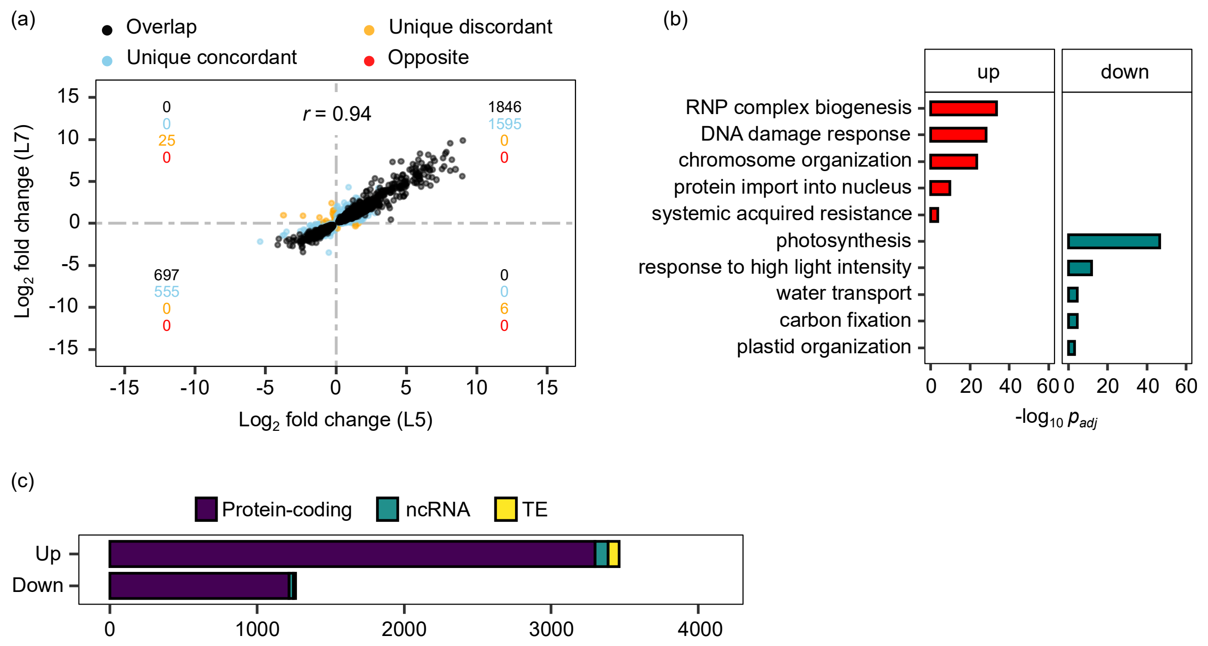


**Supplementary Figure 6. Transcriptome patterns (ribosome-depleted RNA-seq) after enhanced DNA demethylase activity by ROS1**. **(a)** Correlation between $\log_2$ fold-change ($\log_2$FC) values of DEGs (E2 versus DMSO) in two independent *XVE:ROS1-YFP* lines (L5 and L7), yielding a Pearson correlation coefficient ($r$) of 0.94 ($p < 0.001$). Each point represents a DEG identified in E2-treated L5 or L7 classified as 'overlap' (both $p_{adj} < 0.05$; both $\log_2$FC > 0 or both $\log_2$FC < 0), 'unique concordant' (blue) (only one $p_{adj} < 0.05$; both $\log_2$FC > 0 or both $\log_2$FC < 0), 'unique discordant' (orange) (only one $p_{adj} < 0.05$; one $\log_2$FC < 0 and one $\log_2$FC > 0), or 'opposite' (red) (both $p_{adj} < 0.05$; one $\log_2$FC < 0 and one $\log_2$FC > 0). Coloured inserted numbers (from top to bottom) represent the sums of DEGs classified as 'overlap', 'unique concordant, 'unique discordant', and 'opposite', respectively, in each quadrant of the graph (centred around 0, 0). **(b)** Selected simplified statistical gene ontology (GO) terms ($p_{adj} < 0.05$) for biological processes associated with protein-coding genes in the "Up" (red) and "Down" (blue) cluster from the heatmap depicted in Figure 3b. RNP = ribonucleoprotein. All GO terms are listed in Supplementary Data 3. **(c)** Frequency distribution of protein-coding mRNAs (purple), non-coding mRNAs (ncRNA; blue) and transposable elements RNAs (TEs; yellow) within up- and down-regulated RNA clusters (for details, see Figure 3b).

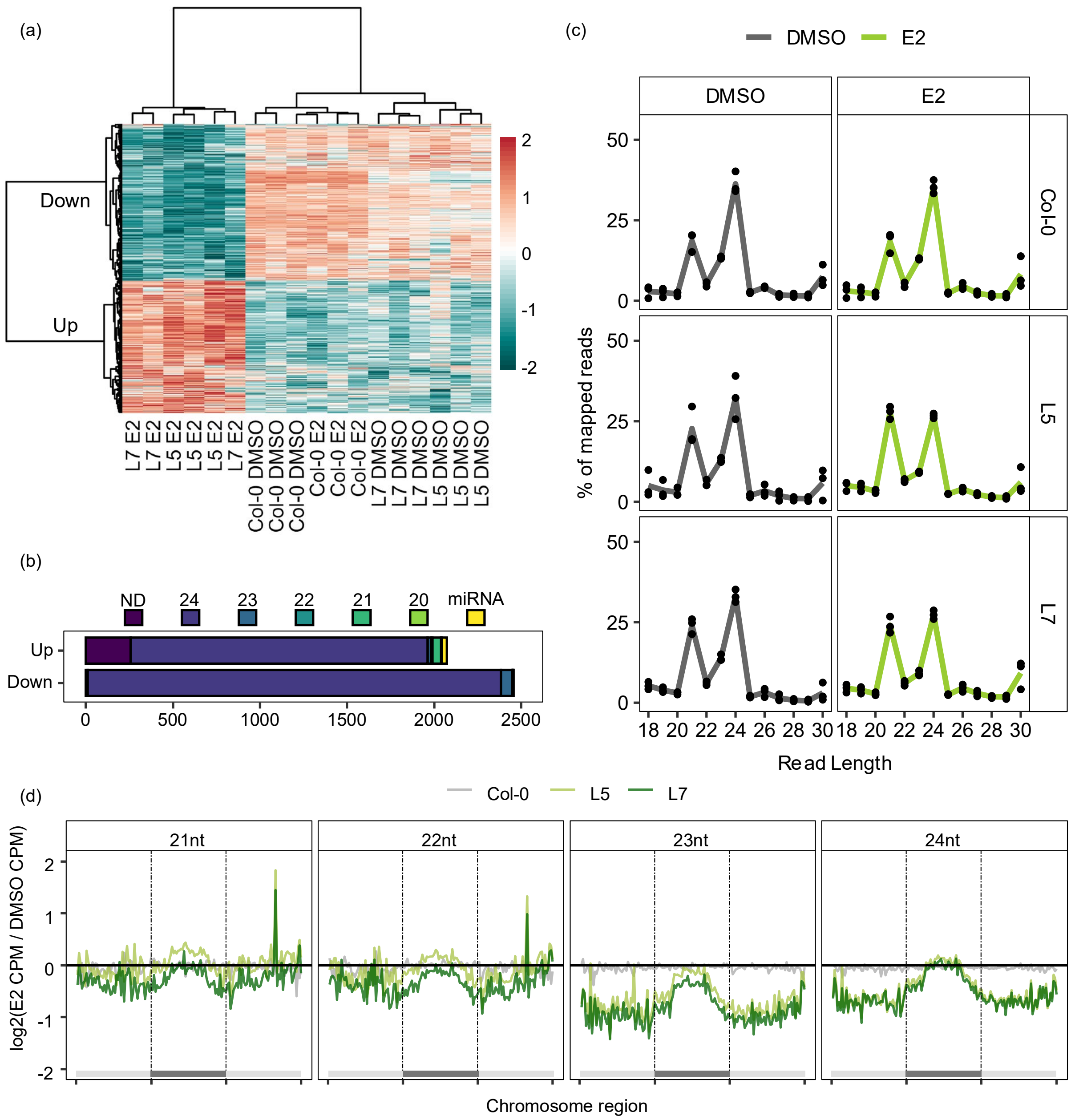


**Supplementary Figure 7. Global sRNA patterns after enhanced DNA demethylase activity by ROS1.** **(a)** Heatmap projection of sRNA abundance of all 4,528 DECs between E2- and DMSO-treated L5, L7, and Col-0 plants, using ShortStack and DESeq2[8,9]. Heatmap-projected values represent per cluster Z-scores of transformed and normalised read counts from 3 biological replicates for each genotype-treatment combination. Scores were clustered by DEC and sample (Euclidean, Ward). Two distinct clusters were identified, labelled “Down” and “Up”, reflecting expression patterns in E2-treated L5/L7 plants relative to all other genotype-treatment combinations. **(b)** Cluster type counts (ShortStack) of all DECs identified in the “Up” and “Down” cluster from the heatmap (a). Clusters where >80% of reads were of a single size, ranging from 20 to 24 nucleotides (nt), were assigned to that specific size category. Any clusters associated with predicted or known miRNAs were assigned as a miRNA cluster. Any other clusters with no dominant read size, or a dominant read size outside of the 20-24 nt range, were classified as not defined (ND). **(c)** The distribution of mapped sRNA read lengths for each genotype-treatment combination, aligned to the TAIR10 Arabidopsis genome by ShortStack. Lines represent the average percentage of mapped reads for each size in the 18-30 nt range from three biological replicates (each depicted as points). **(d)** Chromosome-level metaplots of sRNAs ranging from 21-24 nt in Col-0, L5, and L7 samples. Graphs show mean $\log_2$ fold-changes in counts per mapped-million (CPM), in E2-treated samples relative to DMSO-treated controls. Each chromosome in the TAIR10 assembly was split into 50 equally sized bins for the left arm, pericentromeric region, and right arm. Bins falling between the dashed lines represent pericentromeric regions.

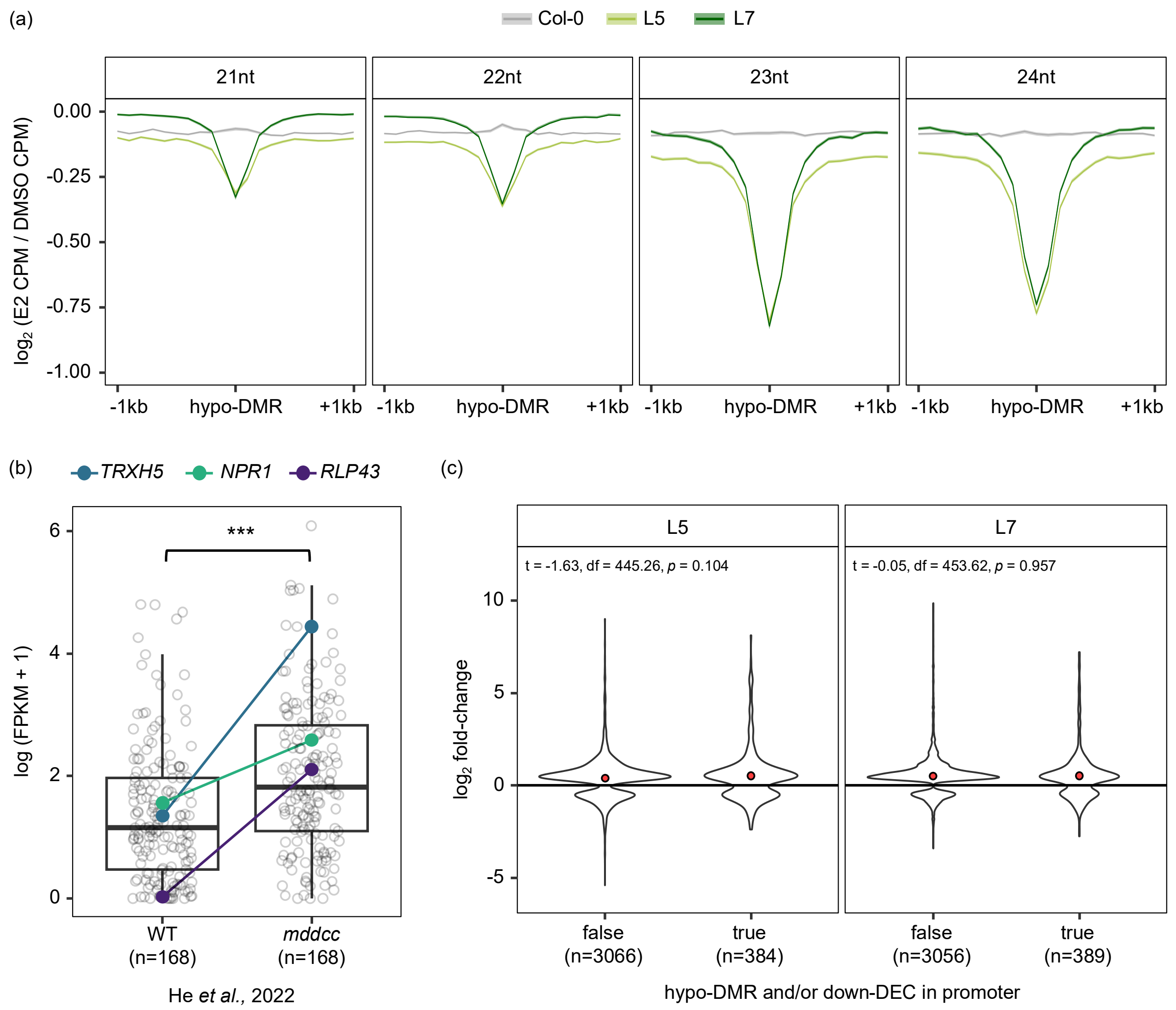


**Supplementary Figure 8. sRNA abundance within hypo-DMRs in the *XVE:ROS1-YFP* lines, transcriptional profiles of predicted *cis*-targets in the hypomethylated *mddcc* mutant, and the association between promoter-region epigenetic changes and differential gene expression in the *XVE:ROS1-YFP* lines. (a)** Metaplot of length-specific (21-24 nt) sRNA abundance within the 4,971 consistent *XVE:ROS1-YFP* hypo-DMRs (Supplementary Figure 5a). Each hypo-DMR was extended 1 kb upstream and downstream and divided into 21 equally sized bins. Solid lines represent the mean $\log_2$ ratio of counts per million mapped reads (CPM) in E2-treated vs DMSO-treated plants for Col-0 (grey), L5 (light green), and L7 (dark green) across all hypo-DMRs and biological replicates. Shaded ribbons represent standard error of the mean. **(b)** Expression of candidate ROS1-RdDM *cis*-regulated genes (defined in Figure 4) in Col-0 and *mddcc* mutants, using reported FPKM values from He *et al* (2022)[10]. Putative cis-regulated genes showed significantly higher expression in *mddcc* compared to Col-0 (Welch's t-test, $t = -5.00$, df = 329, ***$p < 0.001$). Highlighted are three immune-regulatory genes discussed in the main text: *TRXH5* (blue), *NPR1* (teal), and *RLP43* (purple). Five hypothetical-protein genes lacked FPKM values in He *et al.* (2022)[10] and were excluded. **(c)** DESeq2-reported $\log_2$ fold-change values of all differentially expressed genes (DEGs) in L5 and L7 whose promoters contain a hypomethylated DMR and/or a down-regulated 24 nt sRNA (‘true’) versus those that don’t (‘false’). Red points indicate group means. Welch’s t-tests were used to compare groups in L5 and L7, with non-significant ($p > 0.05$) statistics reported at the top of each panel.

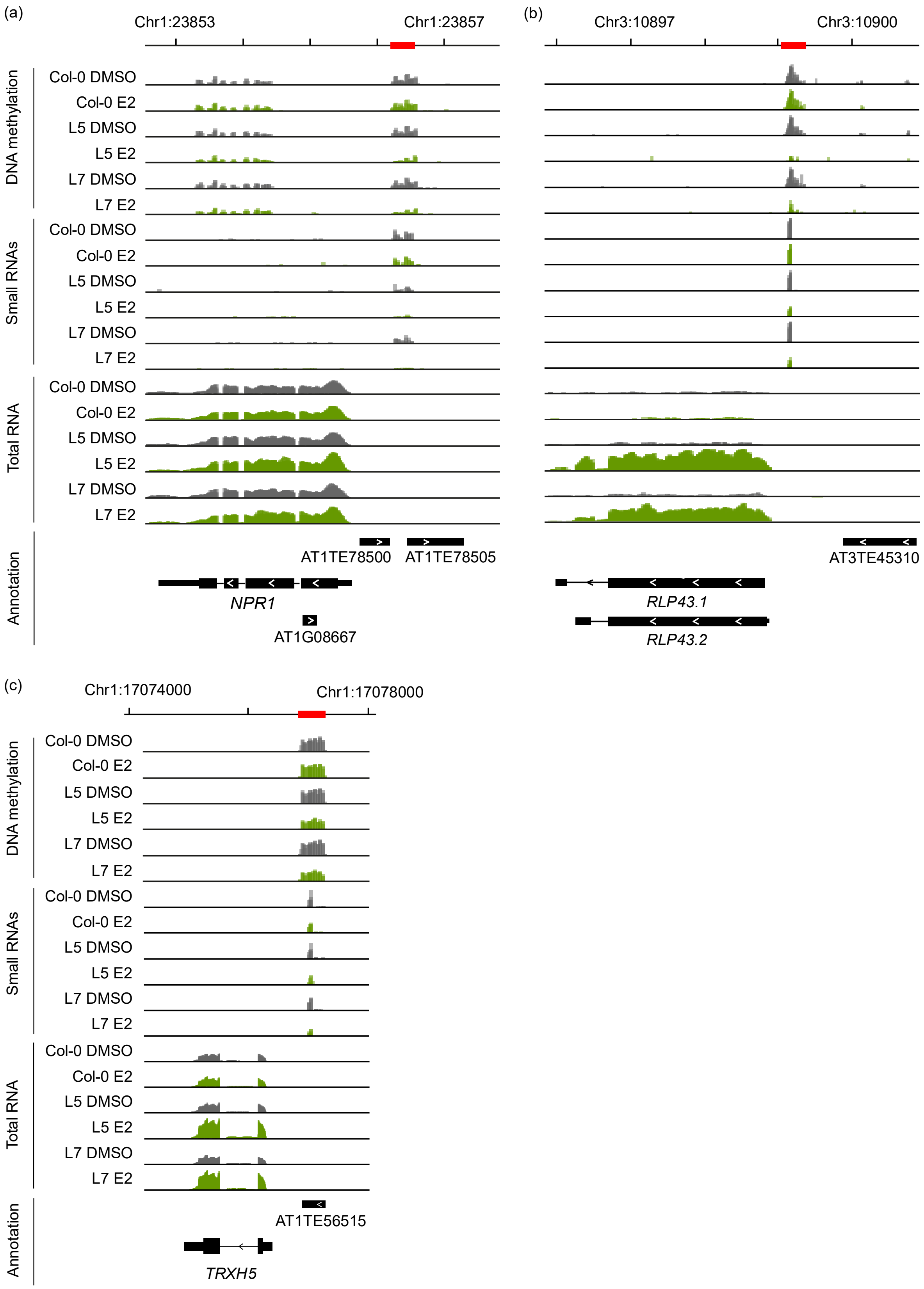


**Supplementary Figure 9. Genome browser snapshots of three *cis*-targeted gene promoters by ROS1.** Genome browser snapshots show DNA methylation (scale = 0-100%), small RNAs (read counts; scale variable, see below), and total ribosome-depleted RNA (read counts; scale variable, see below)

in DMSO- and E2-treated Col-0, L5, and L7. **(a)** *NPR1* promoter DMR and DEC overlapping a TE sequence of E2-treated *XVE:ROS1-YFP* lines (L5, L7). Small RNA count limits = 0-3; RNA count limits = 0-250. **(b)** *RLP43* promoter DMR and DEC in E2-treated *XVE:ROS1-YFP* lines (L5, L7). Small RNA count limits = 0-125; RNA count limits = 0-50. **(c)** *TRXH5* promoter DMR and DEC overlapping a TE in E2-treated *XVE:ROS1-YFP* lines (L5, L7). Small RNA count limits = 0-300; RNA count limits = 0-430. For details of all 173 *cis*-targeted loci, see Figure 4 and Supplementary Data 5. Annotations are based on Araport11[5] and TAIR10 TE annotation files. Each track represents an overlay of all three biological replicates. Figures were created by Integrative Genomics Viewer (IGV)[11]. Red bars at the top highlight DMRs and/or DECs.

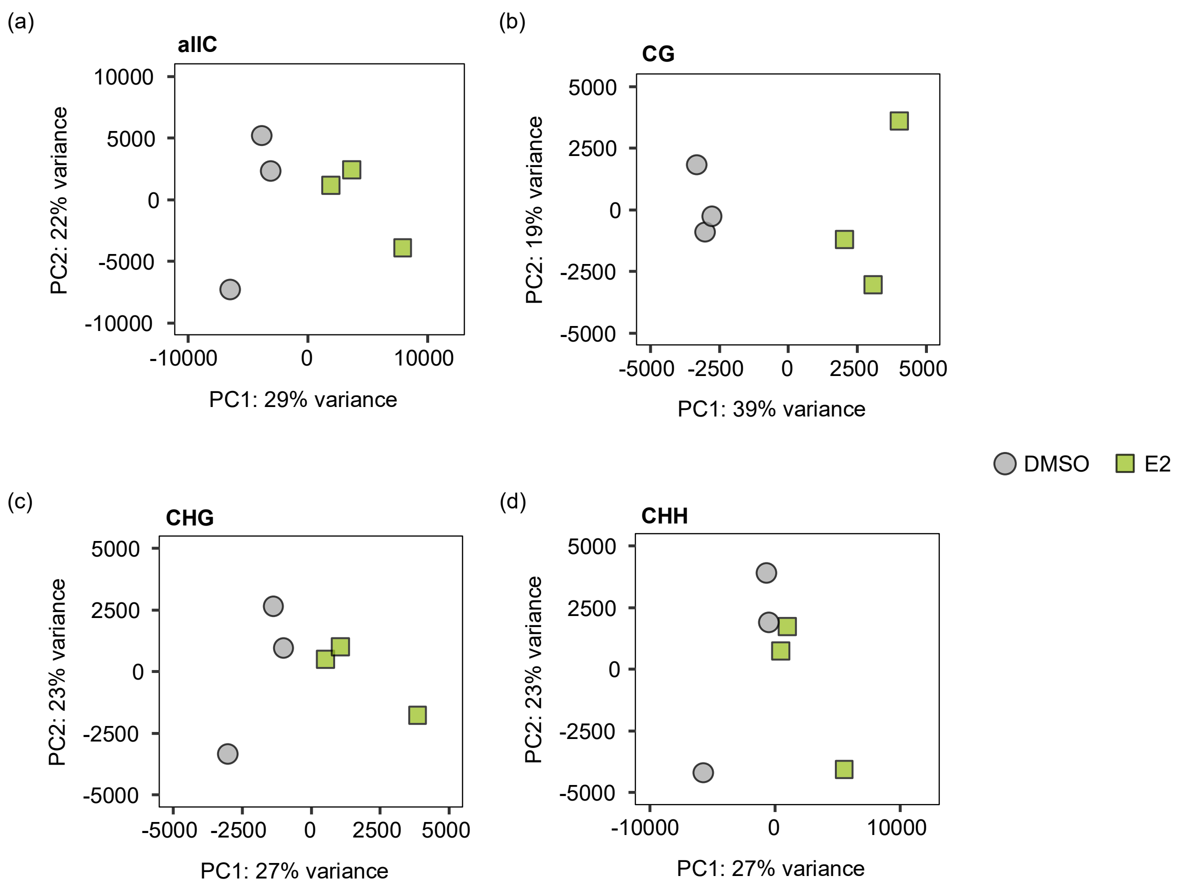


**Supplementary Figure 10. Global DNA methylome analysis by direct long-read Oxford Nanopore Technologies sequencing (ONT-seq) after ROS1 activity.** Principal component analysis (PCA) of cytosine methylation. Plots are based on 200 bp weighted-average windows[4] of cytosine DNA methylation (%) in DMSO-treated (grey circles) and E2-treated (green squares) *XVE:ROS1-YFP* L5 plants. Plants were sprayed twice with DMSO or E2 at 14 days after sowing (DAS) (10 µM E2 or 0.02% DMSO) and at 18 DAS (25 µM E2 or 0.05% DMSO). Cytosine methylation was determined with DeepSignal-Plant[12] and mapped to the Col-CEN reference genome assembly[13]. PCAs were performed for **(a)** all cytosines (allC), **(b)** CG, **(c)** CHG and **(d)** CHH contexts.

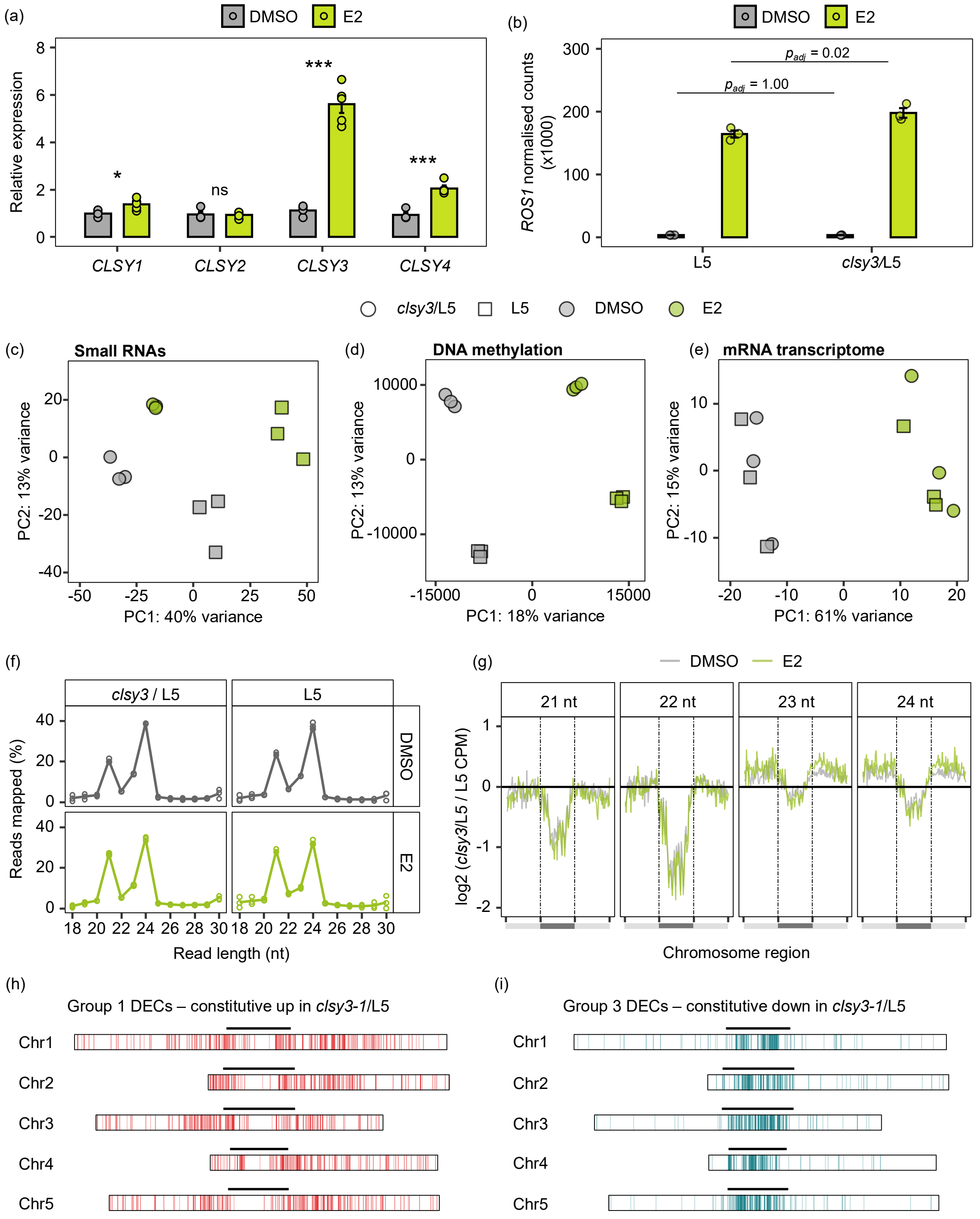


**Supplementary Figure 11. Role of CLSY3 in epigenomic and transcriptomic responses to ROS1 activity (a)** RT-qPCR analysis of samples from an independent experiment with L5 to verify *CLSY* gene expression patterns from the ribosome-depleted RNA-seq analysis (Figure 3b-c; Supplementary Figure 6). Data points represent single biological replicates (n=5 for E2-treated L5; n=4 for DMSO-treated L5), each comprising ~8 plants pooled from a single pot. Relative expression values were normalised to the mean relative expression value of *CLSY1* in DMSO-treated L5. Two-sided t-tests were performed for each *CLSY* gene to assess statistical differences between DMSO and E2 treatments (ns = $p > 0.05$; * $p < 0.05$; *** $p < 0.001$). **(b)** DESeq2 normalised read counts (x1,000) of *ROS1* in DMSO- and E2-treated *XVE:ROS1-YFP* lines in the wild-type background (L5) and in the *clsy3-1* mutant background (*clsy3*/L5). Adjusted *p*-values ($p_{adj}$) are derived from DESeq2 comparisons between genotypes (*clsy3*/L5 vs L5) within a treatment. **(c-e)** PCA plots displaying global variation in normalised abundance of small RNA clusters (c; small RNAs), weighted-average cytosine DNA methylation in 200 bp windows[4] (d), and normalised transcript abundance of coding RNA transcripts (e; mRNA-seq). Squares and circles represent samples from L5 and *clsy3*/L5 plants **(f)** Distribution of mapped sRNA read lengths for each genotype-treatment combination, aligned to the TAIR10 Arabidopsis genome by ShortStack. Lines represent the average percentage of mapped reads for each size in the 18-30 nt range from three biological replicates, which are depicted as points. **(g)** Chromosome-level metaplots of 21-24 nt sRNA profiles. Shown are $\log_2$ fold-changes in counts per mapped-million (CPM) in DMSO- or E2-treated *clsy3*/L5, relative to DMSO- or E2-treated L5, respectively. Each chromosome in the TAIR10 assembly was split into 50 equally sized bins for the left arm, pericentromeric region, and right arm. Bins falling between the dashed lines represent pericentromeric regions. **(h-i)** Genomic locations of differentially expressed sRNA clusters (DECs) characterised as being constitutively upregulated in *clsy3/*L5 (h; group 1 DECs, Figure 6b) or constitutively downregulated in *clsy3/*L5 (i; group 3 DECs, Figure 6b). Boxes represent the five nuclear chromosomes from the TAIR10 genome assembly. Black lines mark pericentromeric regions. For all experiments, plants were treated twice with E2 (green) or DMSO (grey) at 14 days after sowing (DAS; 10 µM E2 or 0.02% DMSO) and 18 DAS (25 µM E2 or 0.05% DMSO), before harvesting shoot material for DNA/RNA extraction at 20 DAS.

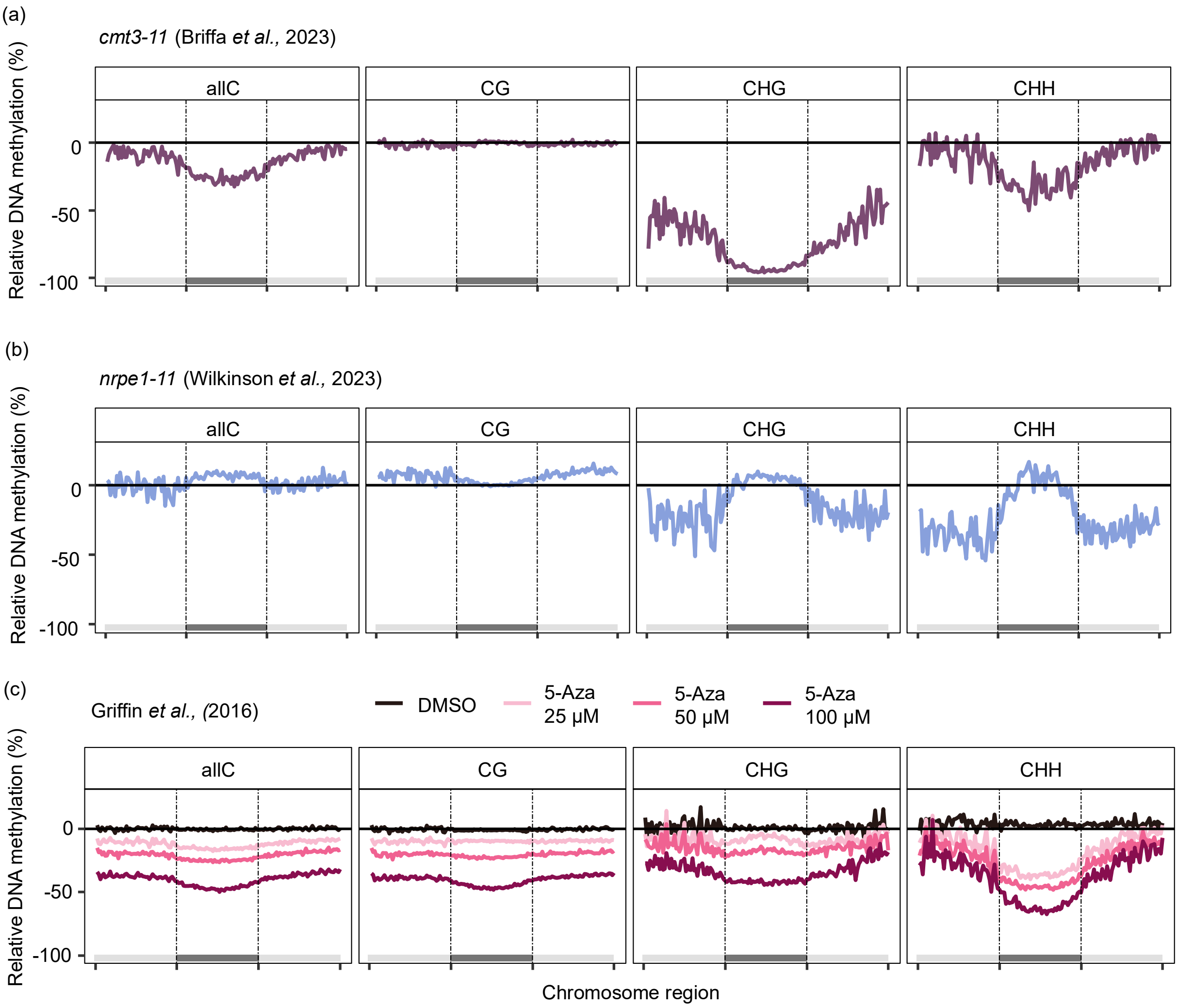


**Supplementary Figure 12. Impacts of the *cmt3-11* mutation, the *nrpe1-11* mutation, and the DNA methyltransferase inhibitor 5-azacytidine on global DNA methylation.** Chromosome-level metaplots of DNA methylation, derived from previously published whole genome bisulfite sequencing (WGBS) of **(a)** *cmt3-11*[14] and **(b)** *nrpe1-11*[15] mutants of Arabidopsis. Values shown are percentages relative to Col-0 control plants. **(c)** Chromosome-level metaplots of Arabidopsis seedlings grown for 10 days on agar plates (including a 2 day stratification period) with DMSO or increasing concentrations of 5-azacytidine (5-Aza)[16]. Values are percentages relative to untreated control plants. For each chromosome, average DNA methylation for a given context was calculated for 50 equally sized bins in the left arm, the pericentromeric region, and the right arm before averaging across replicates (n=3) and chromosomes. Dashed lines and dark grey bars represent pericentromeric bin limits.

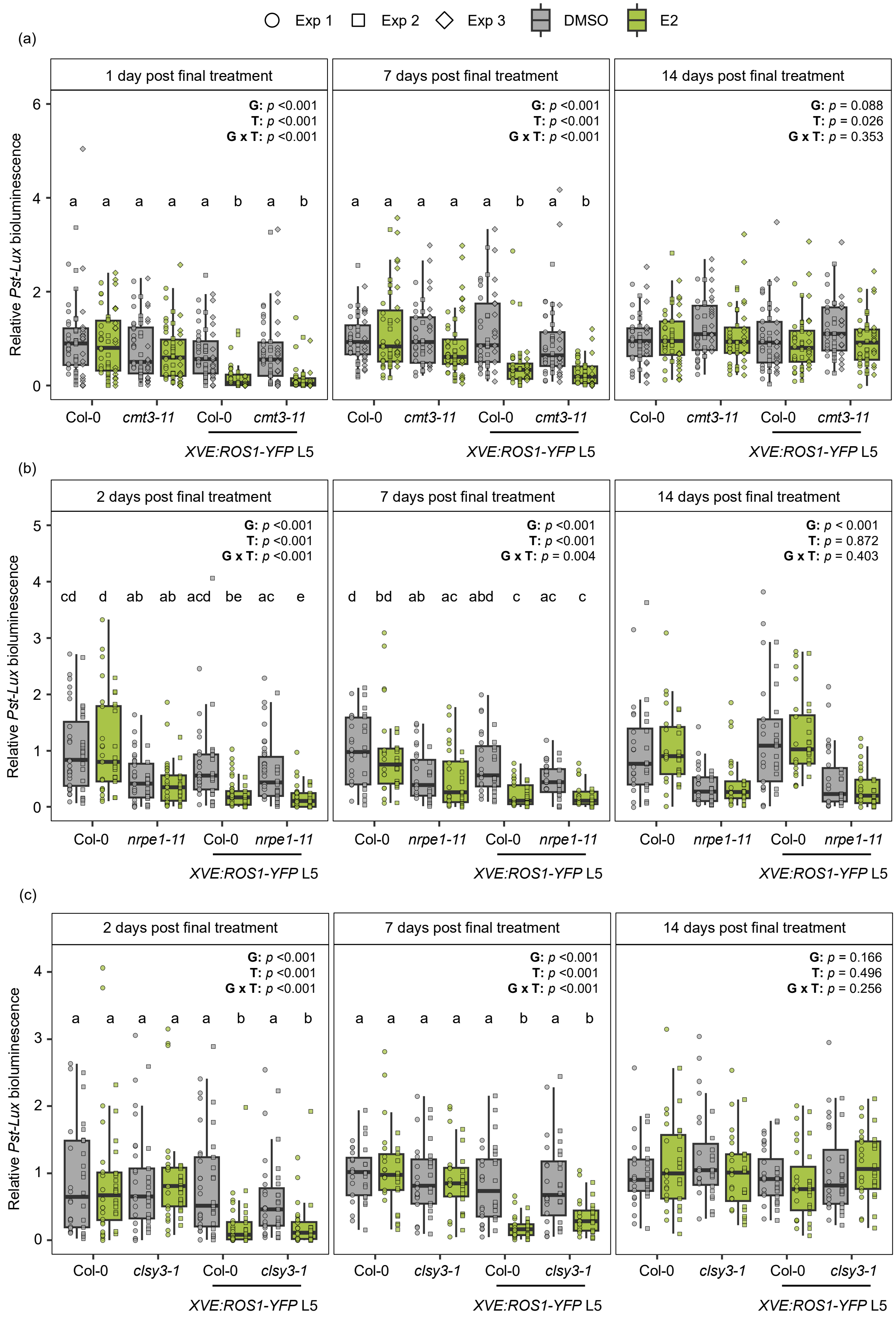

Exp 1
Exp 2
Exp 3
DMSO
E2
(a)
1 day post final treatment
7 days post final treatment
14 days post final treatment
G: p <0.001
T: p <0.001
G x T: p <0.001
G: p = 0.088
T: p = 0.026
G x T: p = 0.353
a a a a a b a b
Relative Pst-Lux bioluminescence
Col-0
cmt3-11
XVE:ROS1-YFP L5
(b)
2 days post final treatment
G x T: p = 0.004
G: p < 0.001
T: p = 0.872
G x T: p = 0.403
cd d ab ab acd be ac e
d bd ab ac abd c ac c
nrpe1-11
(c)
G: p = 0.166
T: p = 0.496
G x T: p = 0.256
clsy3-1


307

**Supplementary Figure 13. Impacts of mutations in DNA methylation machinery on the strength and duration of ROS1-induced resistance against *Pst-Lux*. (a-c)** Col-0 and mutant (*nrpe1-11, cmt3-11,* or *clsy3-1*) genotypes with or without the *XVE:ROS1-YFP* construct from L5 were sprayed twice with DMSO (0.02% and 0.05%) or E2 (10µM and 25 µM), at 14 and 18 days after sowing (DAS). All plants were challenged with *Pst-Lux* at 1-2 days, 7 days, or 14 days post the final E2/DMSO treatment. *Pst-Lux* colonisation was quantified by relative bioluminescence at 2-3 days post-inoculation (dpi)[1]. Data shown represent pooled data from two to three independent experiments for each mutant genotype (*cmt3-11*, *nrpe1-11*, or *clsy3-1*); before pooling, data from each experiment were normalised to the mean *Pst-Lux* bioluminescence values of DMSO-treated Col-0 plants at each given timepoint. Two-way ANOVA was used to assess statistical significance of genotype (G), treatment (T), and their interaction (G x T) on *Pst-Lux* colonisation. Tukey post-hoc analysis was conducted for timepoints at which there was a significant interaction term by the ANOVA ($p < 0.05$). Different letters indicate statistically significant differences between treatment-genotype combinations ($p_{adj} < 0.05$).

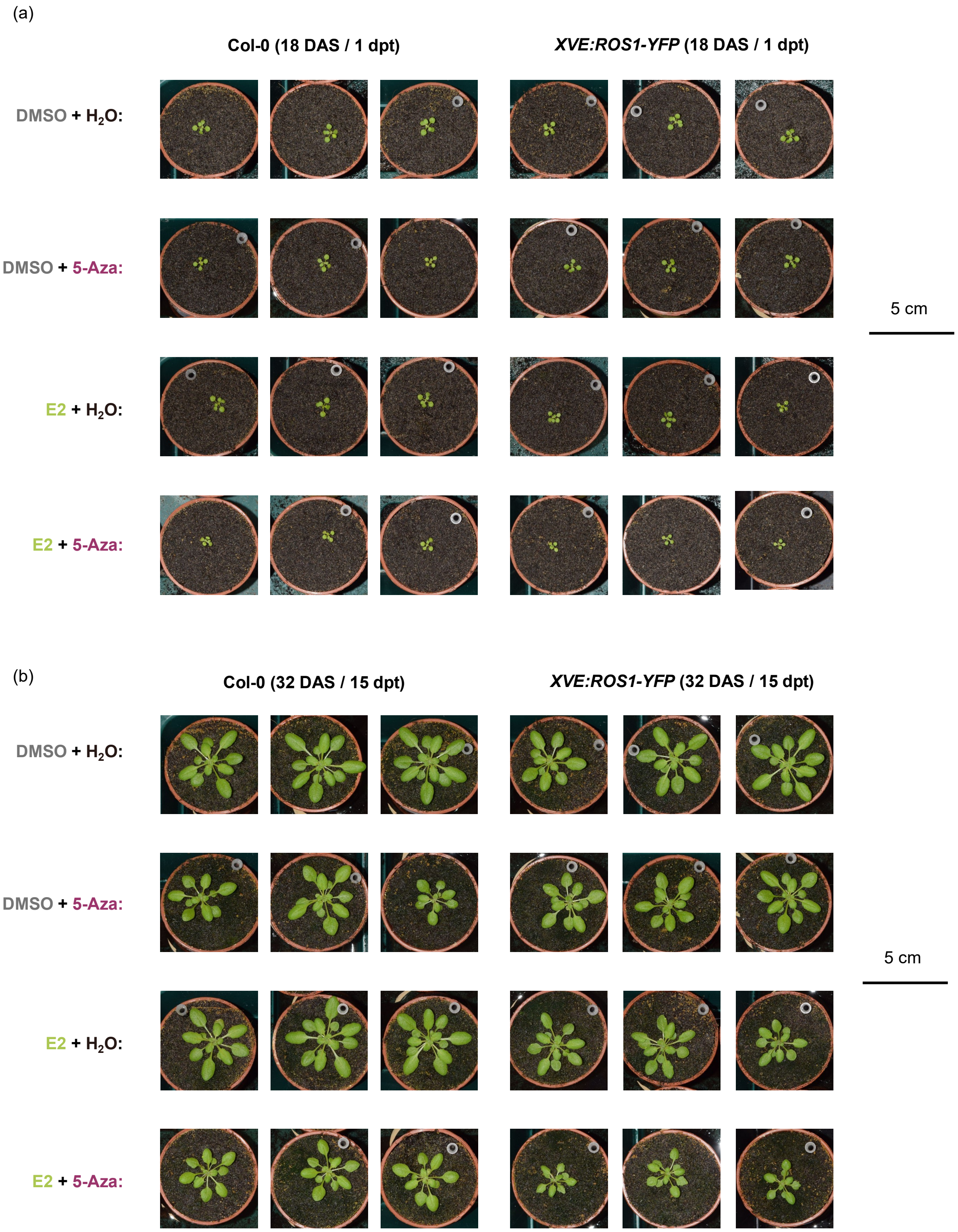


**Supplementary Figure 14: Representative growth phenotypes of Col-0 and *XVE:ROS1-YFP* L5 plants (n=3) upon different treatment combinations.** For details of treatments ($H_2O$/5-Azacytidine and DMSO/E2) see Figure 7a in the main text. **(a)** Top grid shows plants at 1 day after final E2 treatment (1 dpt), which corresponds to 18 days after sowing (18 DAS). **(b)** Bottom grid shows plants at 15 dpt / 32 DAS. The same set of plants are shown in panel (a) and (b). Scale bar = 5 cm.

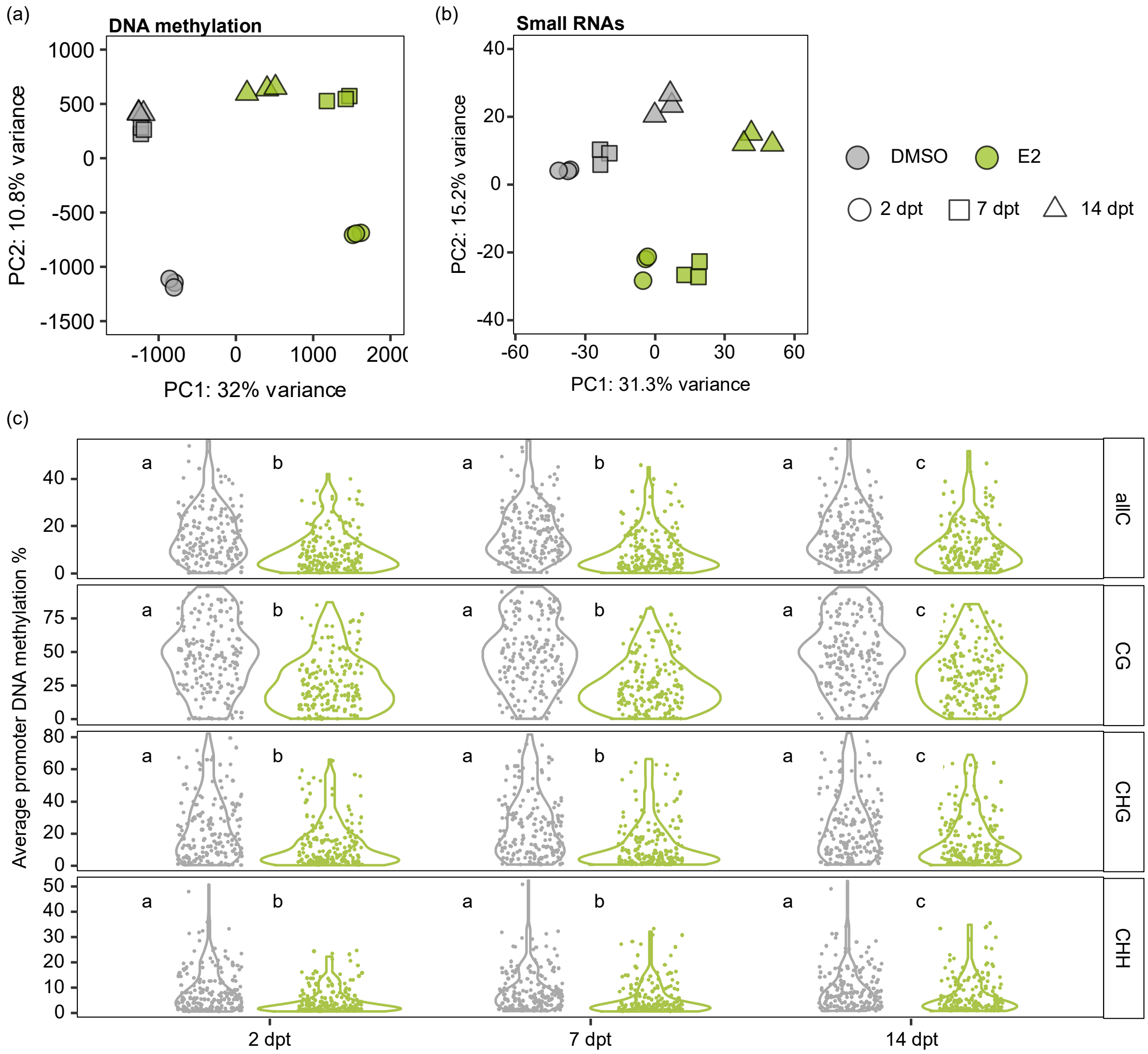


**Supplementary Figure 15: Temporal changes of DNA methylation and small RNAs following ROS1 activity.** *XVE:ROS1-YFP* plants (L5) were treated twice with DMSO (grey) or E2 (green) at 14 days after sowing (DAS; 10 µM E2 or 0.02% DMSO) and 18 DAS (25 µM E2 or 0.05% DMSO). **(a-b)** PCA plots displaying variation in global cytosine DNA methylation, based on 200 bp weighted-average windows[4], (a) and normalised abundance of small RNA clusters (b) at 2-, 7-, and 14-days post treatment (2 dpt, 7 dpt, 14 dpt). **(c)** Average DNA methylation levels in promoters of the 173 *cis*-regulated genes by ROS1 (Figure 4) for each cytosine context (allC, CG, CHG, CHH) at 2 dpt, 7 dpt and 14 dpt after DMSO (grey) or E2 (green) treatments. Mean cytosine methylation was calculated from the WGBS data for each gene and time point. A statistically significant interaction between genotype and treatment was identified by two-way ANOVA for each sequence context ($p < 0.05$). Different letters indicate statistically significant differences between treatment-time points combinations (Tukey $p_{adj} < 0.05$).

### Supplementary tables

**Supplementary Table 1:** Primers used in this study

| Purpose | Name | Nucleotide sequence | Notes |
|---|---|---|---|
| RT-qPCR of ROS1 | ROS1-F1 | AAAACTACCCCTCATCGCTG | Data presented in figure 2e |
| | ROS1-R1 | GTTAGTACGTGCATATTCCAAGC | |
| RT-qPCR of CLSY1 | CLSY1-F1 | GAGGAGGATGATGGCGAGAC | Data presented in Supplementary Figure 11 |
| | CLSY1-R1 | ACTCTCACCTGAAACTCATGGTC | |
| RT-qPCR of CLSY2 | CLSY2-F1 | AATCGGGATGTGTTGCAGGT | |
| | CLSY2-R1 | TCTGGGCTTCTTTGTGGCTC | |
| RT-qPCR of CLSY3 | CLSY3-F1 | TTCTTGACGAGGCACACACA | |
| | CLSY3-R1 | ACATTGCACAGCTCCAGGAA | |
| RT-qPCR of CLSY4 | CLSY4-F1 | AGTTTTGTCGCCCCTTTGGA | |
| | CLSY4-R1 | ACCCTTCTTGTTGGTGTGGG | |
| RT-qPCR of reference gene UBC21 | UBC21-F1 | CTGCGACTCAGGGAATCTTCTAA | Stably expressed gene used for RT-qPCR normalisation[17,18] |
| | UBC21-R1 | TTGTGCCATTGAATTGAACCC | |
| RT-qPCR of reference gene MON1 | MON1-F1 | AACTCTATGCAGCATTTGATCCACT | Stably expressed gene used for RT-qPCR normalisation[17,18] |
| | MON1-R1 | TGATTGCATATCTTTATCGCCATC | |
| Genotyping the T-DNA insertion in clsy3-1 (SALK_040366) | CLSY3 LP | GGATGGGTTTGACGACACCT | LP+RP amplifies wild-type allele. LBb1.3 + RP amplifies T-DNA insertion |
| | CLSY3 RP | AGACGAGTTTTTCCGGTCCC | |
| | LBb1.3 | ATTTTGCCGATTTCGGAAC | |
| Genotyping the T-DNA insertion in *cmt3-11* (SALK_148381) | CMT3 LP | CCCTCAACAATTAACTGACGC | LP+RP amplifies wild-type allele. LBb1.3 + RP amplifies T-DNA insertion |
| | CMT3 RP | ATAAGAGAAGGAGCTGCTGCC | |
| | LBb1.3 | ATTTTGCCGATTTCGGAAC | |
| Genotyping the T-DNA insertion in *nrpe1-11* (SALK_029919) | NRPE1 LP | CATCATTCTCCTAGCAAACAACATG | LP+RP amplifies wild-type allele. LBb1.3 + RP amplifies T-DNA insertion |
| | NRPE1 RP | GACATTTCTCAGCCGCTTTCTC | |
| | LBb1.3 | ATTTTGCCGATTTCGGAAC | |
| Genotyping the *XVE:ROS1-YFP L5* insertion | L5_LP | AAGTCTTTTGCAAGTTCTGAGGT | LP+RP amplifies wild-type allele. RB + RP amplifies *XVE:ROS1-YFP* L5 insertion |
| | L5_RP | TGACAGGTCGTTATGCTGGG | |
| | L5_RB | ATCATGGTGGAAAATGGCCG | |

### Supplementary methods

#### Image processing for *Pst-Lux* bioluminescence analysis

*Pseudomonas syringae* pv. *tomato* DC3000 luxCDABE (*Pst-Lux*) colonisation is proportional to the bioluminescence intensity observed in infected tissues[1,19]. Imaging of *Pst-Lux* colonisation was performed using a G:BOX Chemi XRQ gel doc system and GeneSys v1.8.0 software[1]. Plants (9-12 pots per image) were placed in the dark room of the gel doc for 8 minutes to quench all chlorophyll autofluorescence before measuring bacterial bioluminescence. Images were acquired with a 4-minute exposure time without an external light source, using "4 x 4 (0.25MP)" binning. To estimate rosette area, a matching bright field (BF) image was acquired immediately after, using the internal LED 'Upper White Light' source with a 25-30 ms exposure time. A 2 pence coin was placed in the gel doc for size reference. Images were saved as 16-bit TIFF files, and plants were returned to the climate chamber and maintained at 100% RH to ensure ongoing infection and allow for recurrent analyses at later timepoints. For image analysis, the brightfield image stack was imported (file > import > image sequence) into ImageJ[20] and converted to 8-bit format (image > type > 8-bit). The scale was set in mm per pixel using the diameter of a 2 pence coin (25.9 mm) as reference. Automatic local thresholding was applied to the stack (image > adjust > auto local threshold), using the 'Phansalkar' method with a radius of 30 and the option of "white objects on black background" selected. Plants that were connected after thresholding were separated using a black pencil tool, with a line width of 2, and disconnected leaves were reconnected using a white pencil tool with a line width of 2. Manual removal of non-plant selections (e.g., pot edge and soil debris) was done using the black pencil tool. Composite regions of interest (ROIs), corresponding to the outline of the plants (and thus, the area of the plant rosette), were generated (analyse > analyse particles > add to manager) and saved locally. The bioluminescent image stack was then imported, and the ROIs were overlayed to highlight plant regions. Five additional ROIs of varying size were drawn to select non-plant background regions, and the mean-grey value of these background ROIs were measured for every image in the stack (more > multi measure > untick one row per slice). Background noise levels were removed (process > math > subtract) by taking the mean background mean-grey value plus three times the mean standard deviation of the background mean-grey value across all images. Subsequently, the remaining pixels were used to calculate mean-grey values for each plant-derived ROI.

#### Analysis of *XVE:ROS1-YFP* WGBS data: read processing

Raw sequencing data (100 bp, paired end) were filtered by BGI Genomics using SOAPnuke[21] with the following parameters: '-n 0.001', '-l 20', '-q 0.4', '--adaMR 0.25', '--minReadLen 100'.

This filtering process was conducted to remove adapter sequences and low-quality reads. Reads were aligned to the TAIR10 reference genome (EnsemblPlants release v58) using Bismark's (v0.24.1) default parameters[22]. PCR duplicate reads were removed (function: 'deduplicate_bismark'; options: default parameters) and Bismark was used to generate and summarise binary scores for methylated cytosines and unmethylated cytosines in all contexts for each position (function: 'bismark_methylation_extractor'; options: '--comprehensive', '--bedGraph', '--CX', '--cytosine_report', '--ignore_r2 4'). The unmethylated plastid genome was used to calculate bisulfite non-conversion rates, and all samples from all experiments met the accepted threshold of <2%, with an average non-conversion rate of 0.41 %[23] (Supplementary Data 7).

**Analysis of *XVE:ROS1-YFP* WGBS data: principal component analysis (PCA)**

PCA was performed using weighted-average[4] cytosine methylation levels in 200 bp windows (%) for cytosines in any context (all cytosines; allC). Windows with <5 mapped reads were excluded from analyses. Furthermore, to focus the computational analysis on the most variable regions, windows with standard deviations that were less than the median of the standard deviation across all windows were removed. PCA was conducted in base R v4.2.1 (function: 'prcomp'; options: 'scale. = FALSE', 'center = TRUE') and plots were generated using the R package ggplot2 v3.5.0.

**Analysis of *XVE:ROS1-YFP* WGBS data: metaplots**

For each of the 5 nuclear chromosomes in the TAIR10 reference assembly, pericentromeric regions were defined as described in the main Methods section. For each chromosome and chromosomal region (left chromosome arm, pericentromeres and right chromosome arm), 50 equally-sized bins were created using bedtools v2.31.0 (function: 'bedtools makewindows'; options: '-n 50')[24]. Average cytosine methylation for each bin was then calculated for each context (allC, CG, CHG, CHH) using bedtools (function: 'bedtools map'; options: '-a TAIR10_bins', '-b input_methylation_file.bed', '-o mean,count'). Only cytosines with ≥5 mapped reads were included in the analysis, and any bins which had <60% of the expected number of cytosines for a given context were excluded. Furthermore, only bins that passed all filtering steps in all samples were retained; for analyses involving timepoints (Figure 7d), filtering was applied within each timepoint. Mean cytosine methylation was then calculated across all chromosomes grouping by bin, context, treatment, genotype, and timepoint. Averaged cytosine methylation levels were expressed relative to the average cytosine methylation levels of treatment- or wild-type controls.

For TE and gene metaplots (Supplementary Figure 4a), 20 equally sized bins were created across annotated transposable elements (TEs) and protein-coding genes (transcriptional start site to transcriptional termination site; TSS to TTS) using bedtools (function: 'bedtools makewindows'; options: '-n 20') and the Araport11 annotation[5]. Similarly, the 1 kb region upstream and downstream of the feature were divided into 20 equally sized bins, equating to 50 bp per bin. Average methylation was calculated for each bin and context using bedtools (function: 'bedtools map'; options: '-o mean,count'). Only cytosines with ≥5 mapped reads were included in the analysis.

**Analysis of *XVE:ROS1-YFP* WGBS data: analysis of differentially methylated regions (DMRs)**

The R package DSS v2.46.0[25,26] was used to identify DMRs between treatments or genotypes (Supplementary Data 1). As DSS accounts for read coverage, no prior data filtering was performed. Differentially methylated cytosines were called using the function 'DMLtest' with options 'equal.disp = FALSE' and 'smoothing = FALSE'. The resulting DMC object was subsequently used to call DMRs using DSS (function: 'callDMR'; options: 'minlen = 25', 'dis.merge = 50', 'minCG = 3', 'p.threshold = 0.05', 'data= 0.1', 'pct.sig = 0.5'). This pipeline was performed on cytosines in allC, CG, CHG, and CHH sequence contexts to generate context specific DMRs. Annotation of DMRs were based on Araport11[5] and TAIR10 transposable element annotation file (https://www.arabidopsis.org; accessed 10/02/2024) using the R packages genomation v1.34.0[27] and GenomicRanges v1.54.1[28]. In cases where a DMR overlapped with multiple features, a single annotation was assigned in the following order of preference: exon > intron > promoter > downstream > non-coding RNA > intergenic. Non-coding RNA included any defined transcribed locus in Araport11 that is not a protein-coding gene or a TE. Intergenic regions were defined as regions that do not overlap any of these above defined features. DMRs were annotated as overlapping/not overlapping TEs using the TAIR10 TE annotation file.

For annotation enrichment analyses (Supplementary Figure 5b), ROS1-hypomethylated DMRs (hypo-DMRs) were merged across contexts in E2-treated L5 and L7 plants to form set of non-duplicate hypo-DMRs using the function 'reduce' from the R package GenomicRanges v1.54.1. These merged DMRs were then cross-referenced between both lines and overlapping DMRs (identified with 'findOverlaps' from GenomicRanges) were extended such that the start coordinate was the minimum value between L5 and L7, and the end coordinate was the maximum value between L5 and L7. These so-called 'consistent hypo-DMRs' were then annotated using the Araport11 and TAIR10 TE annotations, as described above. Based on the consistent hypo-DMRs, four random control sets of genomic regions of equal size and number were generated by randomly selecting a chromosome start coordinate in the TAIR10 genome. To account for the varying size of chromosomes, proportional probability weights were assigned to each random region. Annotation of random regions was performed similarly as for the set of consistent hypo-DMRs. Counts of real DMRs mapping to a given annotation type was evaluated for enrichment using a two-sided Fisher's exact test with false discovery rate (FDR) correction, taking the counts from a single random genomic set as the expected count value. Four rounds of tests were performed (using a different random set each time) and averages were taken for the test statistics (fold change, *q*-value). Standard errors of the mean were calculated for the fold change and *q*-value and plotted as error bars. Random genomic regions were generated using base R v4.3.2 for selecting chromosomes (function: 'sample'; options: 'prob = chr_size_proportion') and coordinates (function: 'runif'; options: 'n = 1', 'min = 1', 'max = max_chr_size'). Fisher's exact test was performed using the R package rstatix v0.7.2 (function: 'pairwise_fisher_test'; options: 'alternative = "two.sided"', 'p.adjust.method = "fdr"').

Based on the same set of consistent hypo-DMRs, we quantified endogenous ROS1 protein enrichment within these DMRs and in our four sets of random regions, using two independent and publicly available ChIP-seq datasets[6,7]. $\log_2$ fold enrichment was calculated between Myc-ROS1 samples and corresponding Col-0 controls (Wang *et al.,* 2025)[6], or between ROS1-FLAG samples and corresponding *ros1-4* controls (Deng *et al.,* 2026)[7], using deeptools v3.5.4[29] (function: 'bigwigCompare'; options: '--operation log2'). Enrichment was calculated within each hypo-DMR or random region using 'multiBigwigSummary' from deeptools v3.5.4[29] and averaged across replicates.

**RNA purification for RNA-seq and sRNA-seq**

For all experiments, RNA was extracted from 3 biological replicates per genotype per treatment using ~100 mg aliquots of ground material and 1 mL of TRIzol Reagent (Thermo Fisher Scientific 15596026). Following chloroform-phenol separation, precipitation of RNA from the aqueous phase was performed using 0.8 M sodium citrate and isopropanol at a ratio of 2:1:1, respectively. Following centrifugation for 15 minutes at 12,000 × *g* at 4°C, the pellet was washed twice with 70% ethanol and resuspended in 50 µL nuclease-free water. To aid resuspension of the pellet, tubes were vortexed for 10-15 seconds, incubated at 60°C, and then cooled on ice. The quality of the extracted RNA was first assessed using a Nanodrop 8000 spectrophotometer. In cases where samples had low 260/280 ratios (< 1.7), RNA was reprecipitated overnight at -20°C by adding 0.1x volume of 3 M sodium acetate (pH 5.2-5.5 prepared in nuclease-free H2O) and 2.5x volume pre-cooled 100% ethanol. Centrifugation, wash steps, and resuspension were then repeated as before. Final RNA extracts were checked for quality and quantity using a Nanodrop 8000 spectrophotometer and an Agilent 2200 TapeStation (analysis type: 'Eukaryotic Total RNA QC'). Sequencing libraries for ribosomal RNA depletion (Figure 3) or polyA enrichment (Figure 6) were performed by BGI Genomics using their standard protocols. For sRNA-seq, a gel size-selection was performed for sRNAs 18-30 nt in length. All RNA samples across all experiments had an RNA integrity number (RIN) ≥7.5, as reported by BGI genomics (Supplementary Data 7).

##### Analysis of *XVE:ROS1-YFP* RNA-seq data: read alignment and counting

Raw sequencing data (stranded, 150 bp paired-end) were filtered by BGI Genomics using SOAPnuke[21] with the following parameters: '-n 0.001', '-l 20', '-q 0.4, '--adaMR 0.25', '--polyX 50', '--minReadLen 150'. This filtering process was conducted to remove adapter sequences and low-quality reads. A transcript index of the TAIR10 genome (EnsemblPlants release v58) was generated using STAR v2.7.9a (function: 'STAR'; options: '--runMode genomeGenerate' '--genomeSAindexNbases 12')[30] and a custom annotation file that merged Araport11 annotations[5] and a gene-like annotation file of transposable elements (TEs)[31]. TE loci from Araport11 were excluded from the merged file if they were also annotated by Panda and Slotkin (2020) to prevent multiple annotations of the same locus. Reads were aligned and numbers of reads mapping to each annotated gene were counted using STAR v2.7.9a (function: 'STAR'; options: '--quantMode GeneCounts' '--alignIntronMin 60' '--alignIntronMax 6000' '--outMultimapperOrder Random'). Since this is a reverse-stranded paired library, read counts from the fourth column of the output file were used, which represent the counts where the reverse strand aligned with the annotated locus.

##### Analysis of *XVE:ROS1-YFP* RNA-seq data: library normalisation

For ribosome-depleted RNA-seq (Figure 3), count tables were loaded into R v4.3.2 and filtered to retain only features where ≥6 samples had ≥10 read counts at the given locus. Counts were initially normalised using the default median-of-ratios method implemented by the R package DESeq2 v1.42.1[9,32]. This method assumes that most genes are not differentially expressed, leading to a normalisation factor, or size factor, that typically centres around 1. However, as discussed by Love et al. (2014), there exists a potential "pathological case", where size factors are confounded by extreme experimental conditions. In such cases, features that are truly unchanged in expression between conditions could be incorrectly identified as differentially expressed due to artifacts that are introduced during library normalisation. To investigate this, size factors were extracted in R using DEseq2 v1.42.1 and plotted for samples presented in Figure 3. This revealed larger size factors of E2-treated *XVE:ROS1-YFP* lines compared to all other sample groups (Supplementary Methods Figure 1a). To address issues where size factor is confounded with treatment, DESeq2 implemented the ability for users to generate size factors from a set of features that should not be changing with respect to treatment conditions[9] as is also recommended in the DESeq2 user guide on Bioconductor (guide version 1.42.0).

These features can be internal stably expressed reference genes, which can facilitate more representative library normalisation[33–39]. Accordingly, size factors were regenerated using 15 extremely stably expressed genes in Arabidopsis[17,18] using the R package DESeq2 (function: 'estimateSizeFactors'; options: 'controlGenes = "stableGenes_Czec_and_Hong"'). These 15 genes were selected based on the following criteria: (*i*) the genes were among the 22 most stably expressed genes identified from a dataset covering 323 experimental conditions[17]; and (*ii*) the genes were consistently experimentally verified as stably expressed in 3 independent biological replicates under at least 2 different experimental conditions out of the 6 tested by Hong et al. (2010). The experimental conditions assessed by Hong et al. (2010) included seedling age, photoperiod, ambient temperature, diurnal expression, distinct tissue types, and defined developmental stages. The 15 selected genes used for normalisation were: *AT2G28390*, *AT4G34270*, *AT4G26410*, *AT4G33380*, *AT5G46630*, *AT5G55840*, *AT5G08290*, *AT2G32170*, *AT5G25760*, *AT3G01150*, *AT5G12240*, *AT1G13320*, *AT1G59830*, *AT1G58050*, *AT1G62930*. This normalisation technique reduced the size factors of E2-treated *XVE:ROS1-YFP* lines, resulting in visually improved centring around 1 (Supplementary Methods Figure 1a). Plotting the mean relative expression of these 15 stably expressed genes, based on the change in normalised counts relative to DMSO-treated Col-0 plants, revealed statistically significant differences between treatment groups under the default DESeq2 normalisation method ($H$ = 30.6, df = 5, $p$ < 0.001; Supplementary Methods Figure 2a). Specifically, E2-treated *XVE:ROS1-YFP* lines exhibited a significant reduction in relative expression of these genes compared to all other treatment groups (Conover-Iman test, $p$ < 0.05; Supplementary Methods Figure 1b), confirming that the size factors derived from the default normalisation were systematically inflated, leading to overcorrection of gene expression data and causing artificially reduced normalised count values for E2-treated *XVE:ROS1-YFP* samples. By contrast, no significant differences in relative expression were detected for these 15 genes upon reference-guided normalisation ($H$ = 1.1, df = 5, $p$ > 0.05). Therefore, the control gene normalisation method was used instead. Kruskal-Wallis tests and Conover-Iman tests were performed using base R v4.3.2 (function: 'kruskal.test') and the R package DescTools v0.99.54 (function: 'ConoverTest'). For our mRNA-seq (Figure 6), count tables were filtered to retain only features where ≥3 samples had ≥10 read counts at the given locus before normalising using the same control gene method.

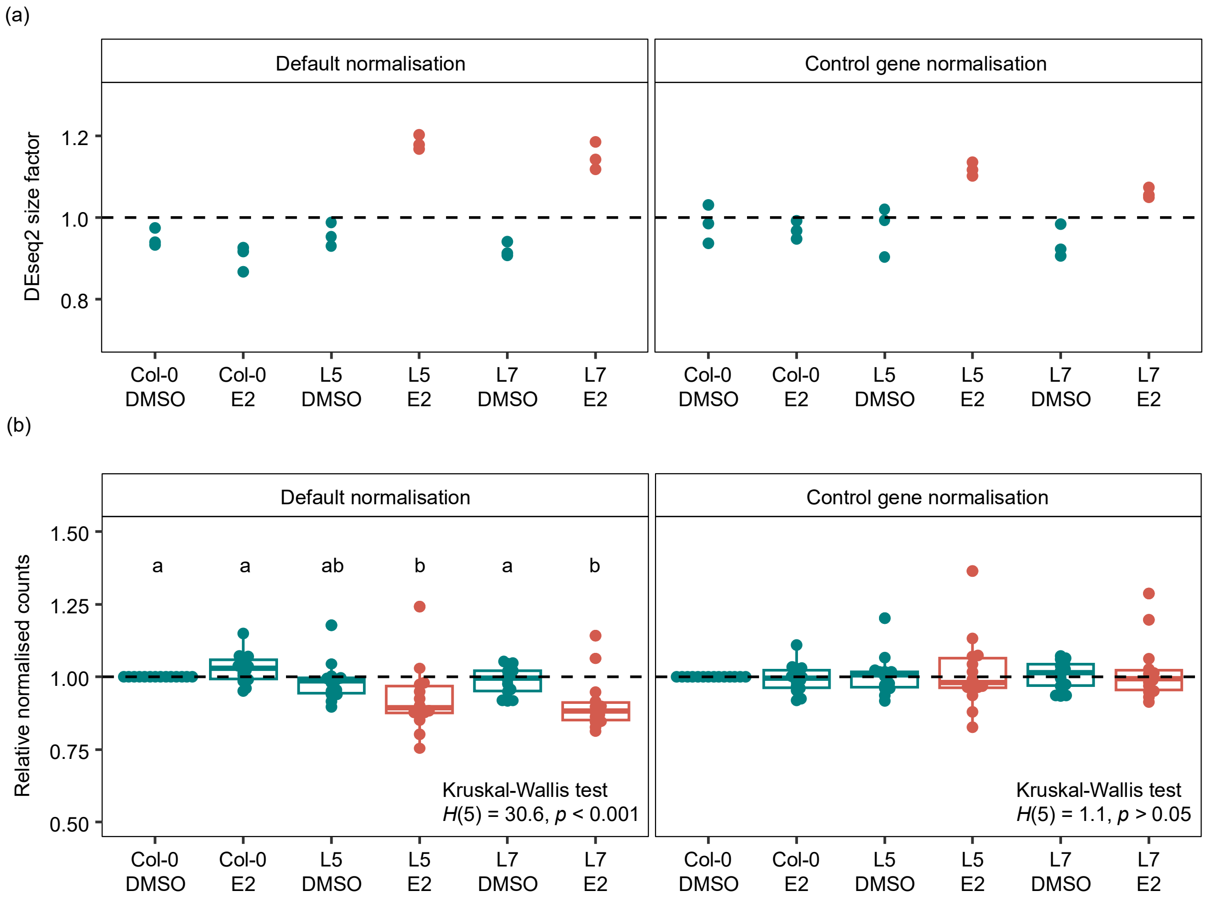


**Supplementary Methods Figure 1**: **(a)** Library normalisation weightings (“size factors”) by DESeq2, assigned to each sample and treatment, using the default median-of-ratios normalising method (“default normalisation”) or a normalisation to a defined set of 15 stably expressed genes in Arabidopsis (“control gene normalisation”). See text for more details. E2-treated *XVE:ROS1-YFP* lines (L5 and L7) are highlighted in red. **(b)** Relative normalised read counts of 15 stably expressed genes in Arabidopsis for each treatment group. Each point represents the mean normalised read count of a single gene, relative to the mean of DMSO-treated Col-0 samples. Conover-Iman post-hoc tests were performed only upon detecting a significant effect of treatment group on relative expression (Kruskal Wallis test, $p < 0.05$). Different letters indicate statistically significantly differences between gene expression values (Conover-Iman, $q < 0.05$).

#### Analysis of *XVE:ROS1-YFP* RNA-seq data: PCA

Normalised read counts were transformed with a regularised logarithm transformation using the R package DESeq2 v1.42.1 (function: ‘rlog’; options: ‘blind=FALSE’)[9]. PCA of the transformed data was conducted using base R v4.3.2 (function: ‘prcomp’; options: default parameters).

#### Analysis of *XVE:ROS1-YFP* RNA-seq data: differentially expressed loci

For our ribosome-depleted RNA-seq (Figure 3), differentially expressed genes (DEGs) ($p_{adj} <$ 0.05) associated with E2 treatment were identified by comparing DMSO-treated and E2-treated plants in Col-0, L5, and L7 using the R package DESeq2 v1.42.1 (model: ~

Genotype_Treatment_Group). For our mRNA-seq with the *clsy3/*L5 genotype (Figure 6e), DEGs were also called using DESeq2 v1.42.1 (~ genotype + treatment + genotype:treatment).

For hierarchical clustering analysis (HCA) of ribosome-depleted RNA-seq DEGs (Figure 3b), rlog-transformed expression values of all identified DEGs across all samples were visualised in a gene-wise and sample-wise clustered heatmap using the R package pheatmap v1.0.12 (function: ‘pheatmap’; options: ‘cluster_rows=TRUE’, ‘cluster_cols=TRUE’, ‘clustering_distance_cols = pearson_distance’, ‘clustering_distance_rows = "euclidean"’, ‘clustering_method = "ward.D2"’). Two distinct gene clusters emerged: one containing DEGs upregulated in E2-treated *XVE:ROS1-YFP* lines (the 'up' cluster), and another containing DEGs downregulated in these lines (the 'down' cluster). The DEGs associated with these clusters were isolated using the ‘cutree’ function from base R v4.3.2.

**Analysis of RNA-seq from *35S:TET1* Arabidopsis lines**

Raw RNA sequencing data from two *35S:TET1* lines and three corresponding Col-0 wild-type controls were obtained from Ji et al., (2018) under GEO accession GSE93024. Adapters and poor quality sequences were removed from reads using trimmomatic v0.39 (options: ‘SLIDINGWINDOW:4:25’, ‘MINLEN:36’, ‘ILLUMINACLIP:adapters/TruSeq3-SE.fa:2:30:10’)[41]. The absence of adapters and the overall quality of the trimmed reads were assessed using FastQC v0.12.1 and MultiQC v1.21[42]. Read alignment, read counting, library normalisation, calling of DEGs (*35S:TET1* vs Col-0), and GO term analysis were performed exactly as described above.

**Analysis of *XVE:ROS1-YFP* sRNA-seq data: read filtering**

Raw sequencing data (50 bp, single-end) were filtered by BGI Genomics using SOAPnuke[21] with the following parameters: ‘-n 0.001’, ‘-l 13’, ‘-q 0.1’, ‘--highA 1’, ‘--minReadLen 15’, ‘--maxReadLen 44’ ‘--ada_trim’. This filtering process was conducted to remove adapter sequences, low-quality reads, reads with 100% adenosine nucleotides, and any reads outside of the range of 15-44 nt. To match the original gel size selection, reads were further filtered for size using TrimGalore v0.6.10 (function: ‘trim_galore’; options: ‘--quality 0’, ‘--length 18’, ‘--max_length 30’, ‘-a X’), retaining only those between 18-30 nt. To remove rRNA reads from the data, reads were mapped to all known *Arabidopsis thaliana* rRNA sequences deposited on RNAcentral (accessed: 23/01/2024)[43] using Bowtie v1.3.1 (function: ‘bowtie’; options: ‘-q’, ‘--un ${input_file}.unmapped_reads.fq’, ‘--best’, ‘-v 0’)[44]. Reads that perfectly matched known rRNA sequences were removed and all unmapped reads were retained for further downstream analyses.

**Analysis of *XVE:ROS1-YFP* sRNA-seq data: identifying sRNA clusters**

ShortStack v4.0.3 software was used to align reads to the TAIR10 reference genome (EnsemblPlants release v58), identify and characterise de novo sRNA clusters, and quantify the number of reads in each sample mapping to these clusters[8,45,46]. ShortStack uses Bowtie v1.3.1[44] for genome indexing and read alignment. Known Arabidopsis miRNAs sequences were downloaded from miRBase (accessed 26/01/2024)[47] and ShortStack v4.0.3 was run with the following parameters: '--genomefile TAIR10.fa', '--known_miRNAs miRBase_ath_known_miRNAs.fasta', '--mmap u', '--dicermin 20', '--dicermax 24', '--pad 200', '--mincov 2'. Using these parameters, ShortStack first identified all predicted and/or known miRNA clusters. It then defined additional clusters based on the dominant read size, only considering those where ≥80% of the reads were of a single size, ranging from 20 to 24 nucleotides (nt). Clusters that did not meet these criteria were classified as "not defined" (ND) clusters.

**Analysis of *XVE:ROS1-YFP* sRNA-seq data: identifying differentially expressed sRNA clusters (DECs)**

Count tables produced by ShortStack were loaded into R v4.3.2 and clusters were filtered for low expression. Specifically, for our experiment using L5, L7, and Col-0 (Figure 3), clusters required ≥10 read counts in ≥6 samples, and for our time course experiment and *clsy3*/L5 experiments (Figure 7, Figure 6), clusters required ≥10 read counts in ≥3 samples. Counts were normalised using the default median-of-ratios method implemented by the R package DESeq2 v1.42.1[9,32]. Normalised read counts were then transformed with a regularised logarithm (rlog) transformation using the R package DESeq2 v1.42.1 (function: 'rlog'; options: 'blind=FALSE')[9]. PCA of the transformed data was conducted using base R v4.3.2 (function: 'prcomp'; default parameters). DECs associated with E2 treatment were identified by comparing DMSO-treated and E2-treated plants in Col-0, L5, and L7 using the R package DESeq2 v1.42.1 (model: ~ Genotype_Treatment_Group).

Hierarchical clustering analysis (HCA) of DECs (Supplementary Figure 7a, Figure 6b), rlog-transformed count values of all DECs were clustered row-wise in a heatmap using the R package pheatmap v1.0.12 (function: 'pheatmap'; options: 'cluster_rows=TRUE', 'clustering_distance_rows = "euclidean"', 'clustering_method = "ward.D2"'). For Supplementary Figure 7a, samples were also clustered column-wise (additional options: 'cluster_cols=TRUE', 'clustering_distance_cols = pearson_distance'). Cluster groups were visually identified and isolated using the 'cutree' function from base R v4.3.2.

**Analysis of *XVE:ROS1-YFP* sRNA-seq data: metaplots**

For each of the 5 nuclear chromosomes in the TAIR10 reference assembly, chromosomes bins for chromosome arms and pericentromeres were created as described for the WGBS data. Aligned sRNA reads were first separated by read size ranging from 21 to 24 nt using samtools v1.2[48] and then converted to '.bed' files using bedtools v2.31.0 (function: 'bedtools bamtobed')[24]. Total mapped sRNA counts for each bin was then calculated for each read-size category (21-24 nt) using bedtools (function: 'bedtools coverage'; options: '-a TAIR10_bins', '-b aligned_sRNA_Xnt.bed', '-counts'). Counts per mapped million (CPM) were calculated as the counts within a bin divided by the total number of mapped reads of that sample, irrespective of size. Mean CPM was then calculated across all chromosomes grouping by bin, read size, treatment, genotype, and timepoint. Averaged CPM values were expressed as a $\log_2$ fold-change relative to the average CPM values of treatment- or wild-type controls.

To generate sRNA metaplots across the 4,971 consistent hypo-DMRs, each hypo-DMR by 1 kb in both directions using 'bedtools slop' (options: '-g TAIR10', '-b 1000') and subsequently divided into 21 equally sized bins using 'bedtools makewindows' (options: '-n 21'). The number of mapped sRNA reads within each bin was quantified using 'bedtools coverage' (options: '-counts'). Mean CPM was calculated per bin, grouping by sRNA size class, treatment, and genotype across all hypo-DMRs and biological replicates. $\text{Log}_2$ ratios of mean CPM in E2-treated over DMSO-treated samples were then calculated for each bin to generate the final metaplot values.

**Analysis of *XVE:ROS1-YFP* L5 Oxford Nanopore Technology sequencing (ONT-seq): library preparation, sequencing, and base-calling**

ONT-seq was performed using ~400 ng of high molecular weight (HMW) DNA extracted from frozen tissue using the NucleoBond® HMW DNA kit (Macherey Nagel, 740160.20) according to the manufacturer's instructions. DNA quality, quantity and size were assessed using a Nanodrop 8000 spectrophotometer, a Qubit 3.0 Fluorometer and the Agilent Femto Pulse system (v1.0.0.32). Library preparation was conducted using the Rapid Barcoding Sequencing kit (ONT, SQK-RBK004) and sequencing performed on a MinION sequencer with FLO-MIN106 flow cells. After 2 days of sequencing, ~37% of FC pores were recovered using the Flow Cell Wash Kit (ONT, EXP-WSH004), and a fresh library preparation was loaded. The FC was then run to exhaustion (~2 days). After sequencing, raw '.fast5' files were base-called and filtered using Guppy v6.5.7 (function: 'guppy_basecaller'; options; '--config', 'dna_r9.4.1_450bps_hac.cfg'). Summary statistics were obtained using NanoPlot v1.39.0 [49].

Methylated cytosines were identified using DeepSignal-plant v0.1.4, as recommended in the GitHub user guide[12] (https://github.com/PengNi/deepsignal-plant). Base-called sequences were annotated onto raw '.fast5' files (function: 'tombo preprocess annotate_raw_with_fastqs'; options: '--basecall-group Basecall_1D_000') and raw signals were aligned to the Col-CEN (v1.2) Arabidopsis reference genome[13] using Tombo v1.5.1 (function: 'tombo resquiggle'; options: '--signal-length-range 0 10000000', '--sequence-length-range 0 1000000', '--basecall-group Basecall_1D_000')[12,50]. Methylation predictions for all cytosines in the genome were called, using DeepSignal-plant with the model 'model.dp2.CNN.arabnrice2-1_120m_R9.4plus_tem.bn13_sn16.both_bilstm.epoch6.ckpt' (function: 'deepsignal_plant call_mods'; options: '--corrected_group RawGenomeCorrected_000', '--motifs C')[12]. The frequency of methylation at each CG, CHG, and CHH site was then called using the 'deepsignal_plant call_freq' function from DeepSignal-plant. The unmethylated plastid genome was used for estimating false-positive methylation calls, with all samples showing an average plastid methylation percentage < 1% and an average methylation percentage across all samples of 0.74% (Supplementary Data 7), which, for reference, is below the accepted non-conversion rate of WGBS data (< 2%)[23].

PCA was performed using weighted-average[4] cytosine methylation levels in 200 bp windows for each context (allC, CG, CHG, CHH). Windows with <5 mapped reads were dropped from the analysis. Furthermore, to focus the analysis on the most variable regions, cytosines with standard deviations less than the median of the standard deviation across all windows were removed from the analysis. PCA was conducted in base R v4.2.1 (function: 'prcomp'; options: 'scale. = FALSE', 'center = TRUE').

**Analysis of *XVE:ROS1-YFP* L5 ONT-seq: chromosome-level metaplots**

For each of the 5 nuclear chromosomes in the Col-CEN reference assembly[13], pericentromeric regions were defined as described in the main text methods. Centromere coordinates were extracted directly from Naish et al (2021). Fifty equally sized bins were created for each chromosome arm flank (left or right relative to centromere), 20 equally sized bins for each pericentromeric flank (left or right relative to centromere), and 20 equally sized bins for each centromere, using bedtools v2.31.0 (function: 'bedtools makewindows'; options: '-n 20' or '-n 50')[24]. CMT2- and CMT3-efficient cytosine positions were defined as cytosines occurring in the sub-contexts CAA and CTA; and CAG and CTG, respectively[51]. Average cytosine methylation for each bin was then calculated for each context (allC, CG) or sub-context category (CMT3-efficient, CMT3-inefficient, CMT2-efficient, CMT2-inefficient), using bedtools (function: 'bedtools map'; options: '-a ColCEN_bins', '-b input_methylation_file.bed'; '-o mean,count'). Only cytosines with ≥5 mapped reads were included in the analysis, and any bins which had less than a third of the expected number of cytosines for a given context/category were excluded. Bins were only considered if they passed all filtering steps in at least 1 sample per treatment. Mean cytosine methylation was then calculated across all chromosomes grouping by bin, context, treatment, and genotype. Finally, these average cytosine methylation levels were expressed relative to the average cytosine methylation levels of DMSO-treated L5 plants.

**Analysis of *XVE:ROS1-YFP* L5 ONT-seq: calling differentially methylated cytosines (DMCs)**

The R package DSS v2.46.0[25,26] was used to identify DMCs between E2- and DMSO-treated L5 using the function 'DMLtest' with options 'equal.disp = FALSE' and 'smoothing = FALSE'. Cytosine methylation was considered significantly different with a $p$-value <0.05 and an absolute difference in methylation >10%. DMC density plots were generated using the R package KaryoploteR[52], using the function 'kpPlotDensity' with a coordinate bin size of 250,000 bp.

**Re-analysis of previously published WGBS data**

WGBS data of *nrpe1-11* and *cmt3-11* mutants were reanalysed from Wilkinson et al (2023)[15] (GSE199625) and Briffa et al (2023)[14] (GSE204837; generation 0 data), respectively. WGBS of 5-Aza-treated Arabidopsis are from Griffin et al (2016)[16] (GSE80300). Analysis of these data were performed as close as possible to the WGBS data from *XVE:ROS1-YFP* lines. However, slight differences in read processing are outlined below, as determined by the method and quality of sequencing.

For *nrpe1-11* data, reads were trimmed for quality using Trimmomatic v0.39[41] (options: 'SLIDINGWINDOW:4:25', 'HEADCROP:6', 'MINLEN:36'). For *cmt3-11* and 5-Aza data, reads were trimmed for quality and adapters using Trimmomatic v0.39[41] (options: 'SE', 'SLIDINGWINDOW:4:25', 'MINLEN:36', 'ILLUMINACLIP:TruSeq3-SE.fa:2:30:10').

For chromosome-level metaplots, filtering steps were performed as described for the *XVE:ROS1-YFP* WGBS data. However, due to lower coverage in some *nrpe1-11* samples, bins were kept given they contained >25% of expected cytosines (rather than 60%) for a given bin and context. Furthermore, for *cmt3-11* (n = 2) and *nrpe1-11* (n = 4) data, bins were kept so long as they passed filtering steps in at least half of the replicates per genotype. For 5-Aza-treated Col-0 data, bins were only kept if they passed filtering for all treatment doses.
